# A dynamic encoder-like protein hub reconciles specialization and plasticity through subcellular mobility

**DOI:** 10.64898/2026.05.23.727322

**Authors:** Carmen Sánchez Moreno, Renée A. Duckworth, Alexander V. Badyaev

## Abstract

How does a conserved regulatory network produce locally specific outcomes across many developmental contexts? Flexible subcellular compartmentalization of proteins can be a powerful source of dynamic modularity, in which proteins mobility links parts of a network, opening alternative regulatory states. Using a comprehensive protein-expression atlas of avian beak morphogenesis, we show that a craniofacial network resolves into a stable core and a hub of multivalent proteins whose links to the core depend on where in the cell they colocalize. We show that this bow-tie architecture doubles the number of regulatory states the network can express at any given stage and, across development, allows nearly complete exploration of its regulatory potential without adding or removing elements. These proteins behave as dynamic encoder-like hubs whose localization-dependent interactions convert diverse mechanical and biochemical inputs into specific outputs, such that specialization and changeability increase together rather than trade off as is commonly assumed. This organization lets a compact, conserved network sustain continuous and diversifying development, exemplifying a mechanistic reconciliation of ecological precision and the evolutionary lability of avian beaks.

## Introduction

Evolution simultaneously favors the robustness needed for the maintenance of current functionality and the flexibility needed to accommodate change. The architectural requirements and origins of complex systems that reconcile these dual demands are debated [1, 2]. A potential empirical resolution can be inferred from developmental multiplexing, in which a conserved regulatory network concurrently orchestrates alternative coexpression states, such as during tissue diversification, while preserving the overall stability of continuous development [3–5]. In these networks, a combination of the diversity and variability of inputs with the specificity required of the outputs favors evolution of an hourglass or bow-tie organization [6–8], where diverse inputs are converted into a narrower set of common rules at the regulatory waist before being routed to downstream pathways [9, 10]. Both the inputs and outputs may be structurally modular, whereas the middle layer, comprising either conserved elements (in a traditional bow-tie) or context-sensing ones (in a “rheostat” bow-tie), orchestrates the dynamic recombination of subcircuits while maintaining network cohesion [11–13]. The resulting dynamic modularity reconciles the accommodation of novel inputs with continuity of development and evolution [2, 14, 15], but the central question remains as to the mechanisms by which elements at the regulatory waist sense, convert, and propagate diverse and often novel inputs [16].

The cell is a fundamental unit in which a multitude of biological, chemical and physical inputs are converted into discrete decisions to differentiate, proliferate or migrate, and subcellular compartmentalization is a powerful organizer of regulatory networks that guide these decisions [17–20]. Context-sensing links that connect or disconnect subcellular neighborhoods can transmit inputs and switch between a network’s regulatory states without gain, loss, or rewiring of its elements [21–25]. Because such bidirectional switches are reversible and combinable, they can form bow-tie type networks that unlock latent regulatory subcircuits that can be explored simultaneously, greatly increasing the range of regulatory states even a small network can achieve; the same core group of regulators can generate a multitude of outcomes [26, 27].

Because subcellular compartmentalization is linked to cell shape and mechanical state [28, 29], the regulatory networks bridging subcellular compartments are closely tuned to the physical inputs that a cell experiences [30]. Thus, for these context-sensing bridges, evolution should favor multivalent molecular sensors able to interchangeably accommodate mechanical and biological signals and convert them into precise molecular interactions within a cell [31]. Understanding the rules by which such sensors compress and convert inputs across physical and biological domains is a central question in biology [10, 16] and is key to reconciling specialization with flexibility in morphogenesis.

Here we examine these rules by investigating the flexible subcellular compartmentalization of proteins in a deeply conserved regulatory network that underlies avian beak evolution [32] (Fig. 1a). Modulation of the key signaling pathways in this network drives avian beak diversification on time scales ranging from ecological plasticity to species divergence [33–38]. Although subcellular compartmentalization is structurally poised to modulate the regulatory interactions of these proteins (Fig. 1a), its role in these diversifications has not been explicitly established.

**Fig 1.**
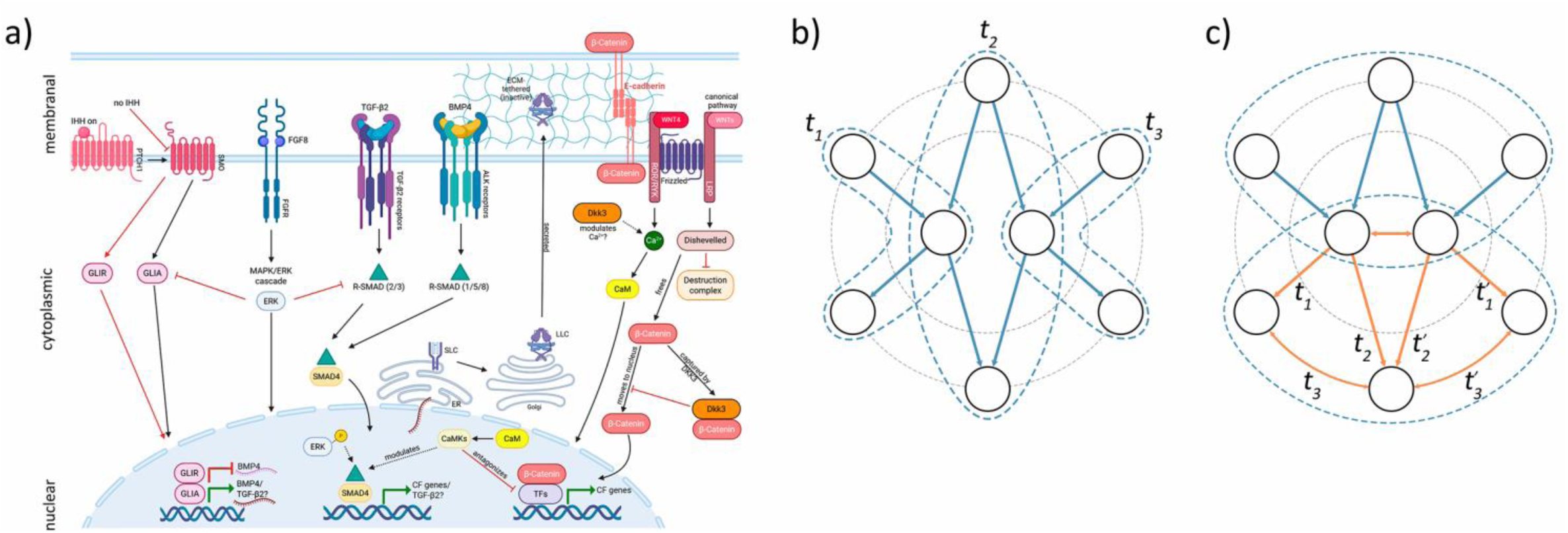
Subcellular compartmentalization of proteins unlocks alternative regulatory states in a core craniofacial network. **(a)** Functional interactions among proteins of the network are mediated by their subcellular colocalization (nuclear, cytoplasmic, or membranal) and by location-specific interactions with shared co-factors [39]. Ihh, Bmp4, Tgfβ2, and Fgf8 converge on transcription factors GLI and R-SMADs whose nuclear translocation is regulated in the cytoplasm. Fgf8-activated ERK intercepts GLI-A and R-SMADs before nuclear entry; once in the nucleus, ERK can phosphorylate R-SMADs at linker sites, altering their co-factor recruitment and redirecting transcriptional output to different target genes, making cytoplasmic–nuclear partitioning the key regulatory step. Wnt4/Ca²⁺ signaling activates CaM and CaMKs that modulate SMADs and other transcription factors in either compartment, antagonizing the parallel canonical Wnt axis in which β-catenin must escape the cytoplasmic destruction complex to enter the nucleus and partner with TCF/LEF at craniofacial target genes. Dkk3 tunes this balance by capturing cytoplasmic β-catenin and modulating Ca²⁺ production, while a membrane-bound β-catenin sequestered at E-cadherin adherens junctions further limits its nuclear availability. Tgfβ2 adds a distinct spatial layer: processed through the ER and Golgi into a latent complex tethered to the ECM, its signaling activity depends on extracellular release before any intracellular cascade can be initiated (see text) [39]. **(b)** Sequential (*t*_1_-*t*_3_) subsampling of a regulatory network allows tissue specialization (blue dashed lines) but compromises long-term network stability and developmental potential. **(c)** Bow-tie architecture where context-dependent outputs (orange edges) allow simultaneous access to alternative regulatory states (*t_n_ vs t_n_’*) allows sustained developmental diversification and expansion. This architecture predicts that context-sensing proteins should concentrate in a narrow layer linking context-invariant layers and the regulatory plasticity of the sensing layer should remain constant while the periphery diverges in development. We specifically test whether context-sensitivity covaries with subcellular colocalization of proteins.

We first construct a comprehensive atlas of protein expression during avian beak development, documenting the anatomical, histological, and subcellular expression of eight core regulatory proteins throughout the development of the upper and lower beak of a passerine bird (Figs. S1-21, Tables S1 and 2), and use it to examine how subcellular compartmentalization modulates the network across developmental contexts (combinations of ontogenetic stages, locations, and tissue types, Table S1). We then contrast three architectures that could account for the network’s ecological responsiveness despite its conservation: progressive subsampling of the network, recombination of fixed tissue-specific modules, and dynamic modularity generated by context-sensing relocation (Fig. 1b,c). The dynamic modularity model predicts that proteins with the most variable subcellular compartmentalization would form a context-sensing hub in bow-tie linkage with the compartment-stable proteins, and that the regulatory plasticity of this hub would be maintained throughout morphogenesis while the peripheral subcircuits accumulate tissue-specificity (Fig. 1c).

We find that the subcellular colocalization of proteins acts as a context-sensing switch that unlocks regulatory states sustaining the network’s dynamic modularity throughout developmental expansion. This organization enables the network to simultaneously express alternative regulatory states without loss of stability while also potentially priming developmental transitions. In such an architecture, specialization and robustness do not trade off with changeability and continuity: a mechanistic reconciliation of the ecological precision with the evolutionary lability evident in avian beak diversification.

## Results

### Context-sensitivity of protein coexpression covaries with their subcellular compartmentalization

In both upper and lower beaks, the protein network consists of a stable core of proteins whose coexpression is invariant across tissue types, jaw locations, and developmental stages (*n* = 11 protein pairs in the upper beak and *n* = 10 in the lower) and a group of proteins whose coexpression sign switches across developmental contexts (upper: *n* =17 protein pairs, lower: *n* =18; Fig. 2a,b). For example, all protein pairs involving CaM and Tgfβ2, and most pairs involving Fgf8 alternate between positive and negative coexpression depending on developmental contexts, whereas interactions among Ihh, Bmp4, Dkk3, and Wnt4 were consistent across all tissues and locations throughout development (Fig. 2a, b; Table S3). Context-invariant and context-sensing protein sets were similar between the upper and lower beak except for Dkk3-Wnt4 and Dkk3-Fgf8 (context-sensing only in the lower beak) and Fgf8-Ihh (context-sensing only in the upper beak). The two groups differ in subcellular colocalization (Fig. 2c, d): context-invariant pairs predominantly involved proteins colocalized in the cytoplasm (e.g., Bmp4-Dkk3, Bmp4-Ihh, Dkk3-Ihh, and Ihh-Wnt4) in both jaws, whereas context-sensing associations involved proteins with variable localization across three to four subcellular compartments (Fig. 2c, d).

**Fig 2.**
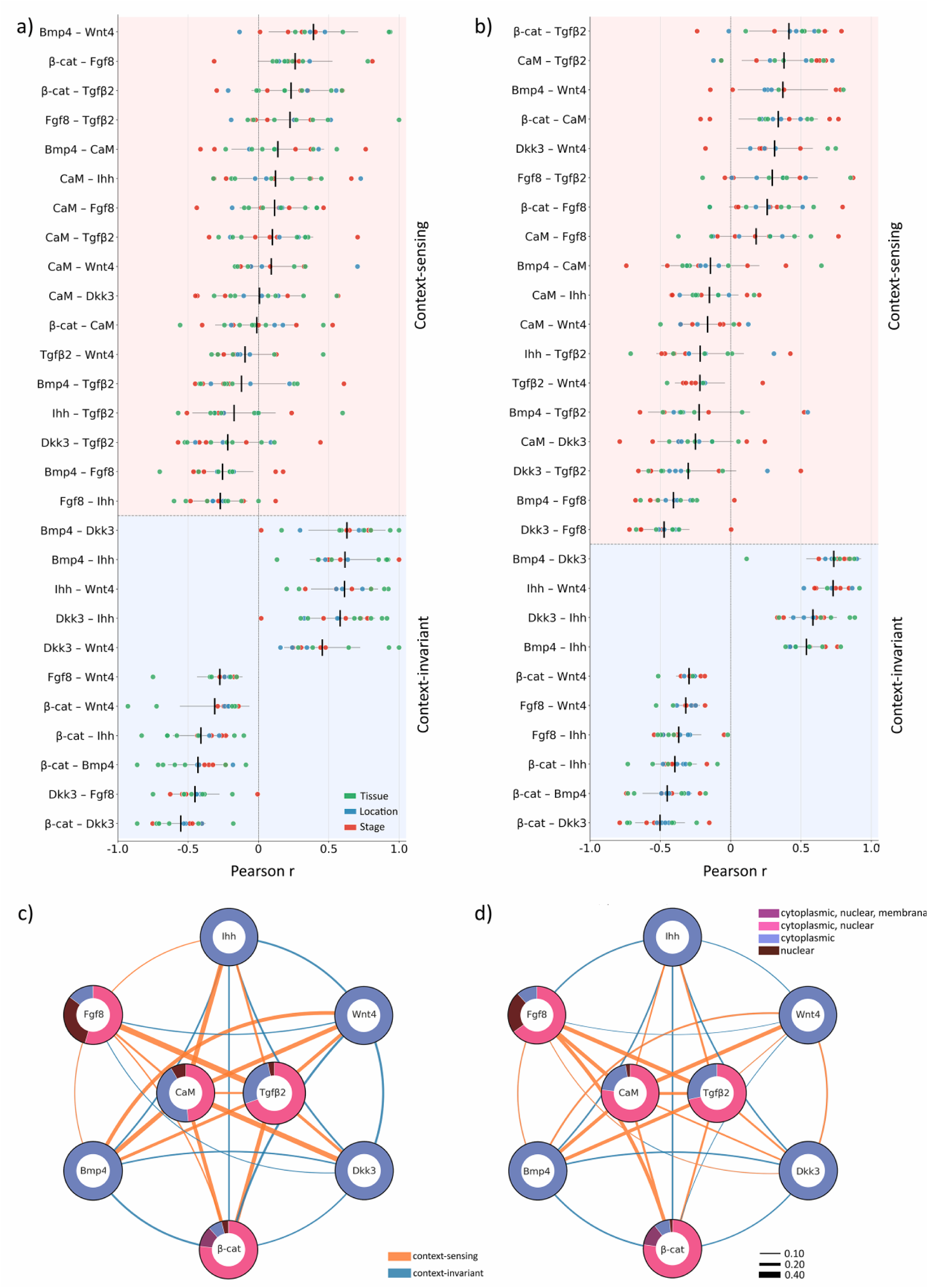
Developmental context of protein colocalization is associated with their subcellular colocalization. Regulatory network comprises proteins with context-invariant coexpression and proteins whose correlations switch signs across contexts in **(a)** upper and **(b)** lower beak. Compartment colocalization of proteins covary with developmental context. Shown are mean Fisher’s *z-*transformed Pearson *r*’s (Table S3) for tissues (green), locations (blue), and developmental stages (red; combining within-stage tissues and locations) for each protein pair. Vertical black lines show mean *r* across all contexts and stages, horizontal gray line spans ±1 s.d. Edge thickness in protein networks for **(c)** upper and **(d)** lower beaks shows variation (s.d.) of correlations across all developmental contexts and stages. Context-invariant correlations (from panels a and b) are shown with blue lines, context-sensing correlations – in orange. Nodes show proportion of protein subcellular compartmentalization (perinuclear and nuclear combined) across samples.

### Context-sensing proteins bridge regulatory states

Within a developmental stage, tissue- and location-specific subnetworks have highly divergent correlational structures (Figs. S22-S23, Table S4), such that averaging often orthogonal protein coexpression either at the whole-cell level or within a developmental stage masks lower-level patterns. For example, the upper beak shows largely discontinuous and sparse stage-specific networks linked across stages by only two shared protein links, whereas the densely interconnected stage-specific networks of the lower beak are linked by only three shared links (Fig. 3). Yet, both patterns, although reflecting genuine temporal differences in morphogenesis (see below), are the result of stage-pooling of orthogonal and dynamic context-specific networks (Fig. 4a): context-sensing pairs of proteins drive switching between the tissue subnetworks within each stage (Fig. S22, Table S4). In contrast, context-invariant protein pairs act as “developmental scaffolds” – their coexpression is similar across tissues and stages forming diagonals in Fig. 4 a, b. Centroids of both context-invariant and context-sensing protein groups are displaced from (0.5, 0.5) midpoint in both jaws (Fig. 4a, b), confirming that the stage-pooled network is an aggregate of distinct tissue-specific subcircuits.

**Fig 3.**
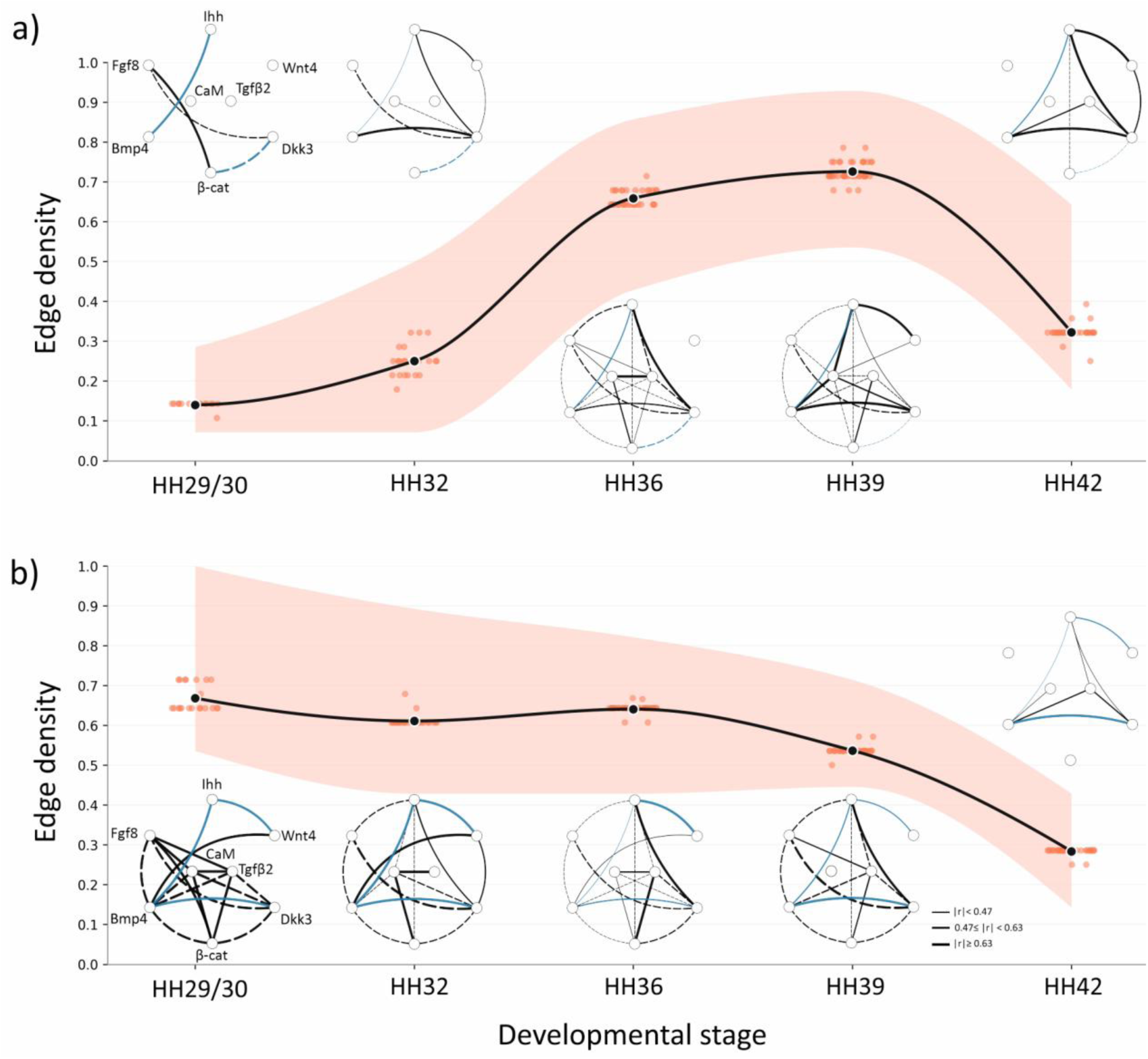
Stage-pooled protein associations mask within-stage dynamic modularity. In both **(a)** upper and **(b)** lower beaks, blue edges show protein associations that persist throughout the developmental sequence; black edge thickness indicates significant within-stage absolute means of correlations. Dashed lines indicate negative correlation, solid lines – positive correlation. Developmental multiplexing within each stage accounts for missing (masked) edges (Fig. 4). Black dots are means of within-stage bootstrap resamplings shown as red dots. Confidence intervals (CI) are 95% bootstrap resampling with replacement. Edge density is the ratio (out of 28 total in the network) of significant edges (*P* < 0.05; Benjamini-Hochberg multiple comparison correction).

**Fig 4.**
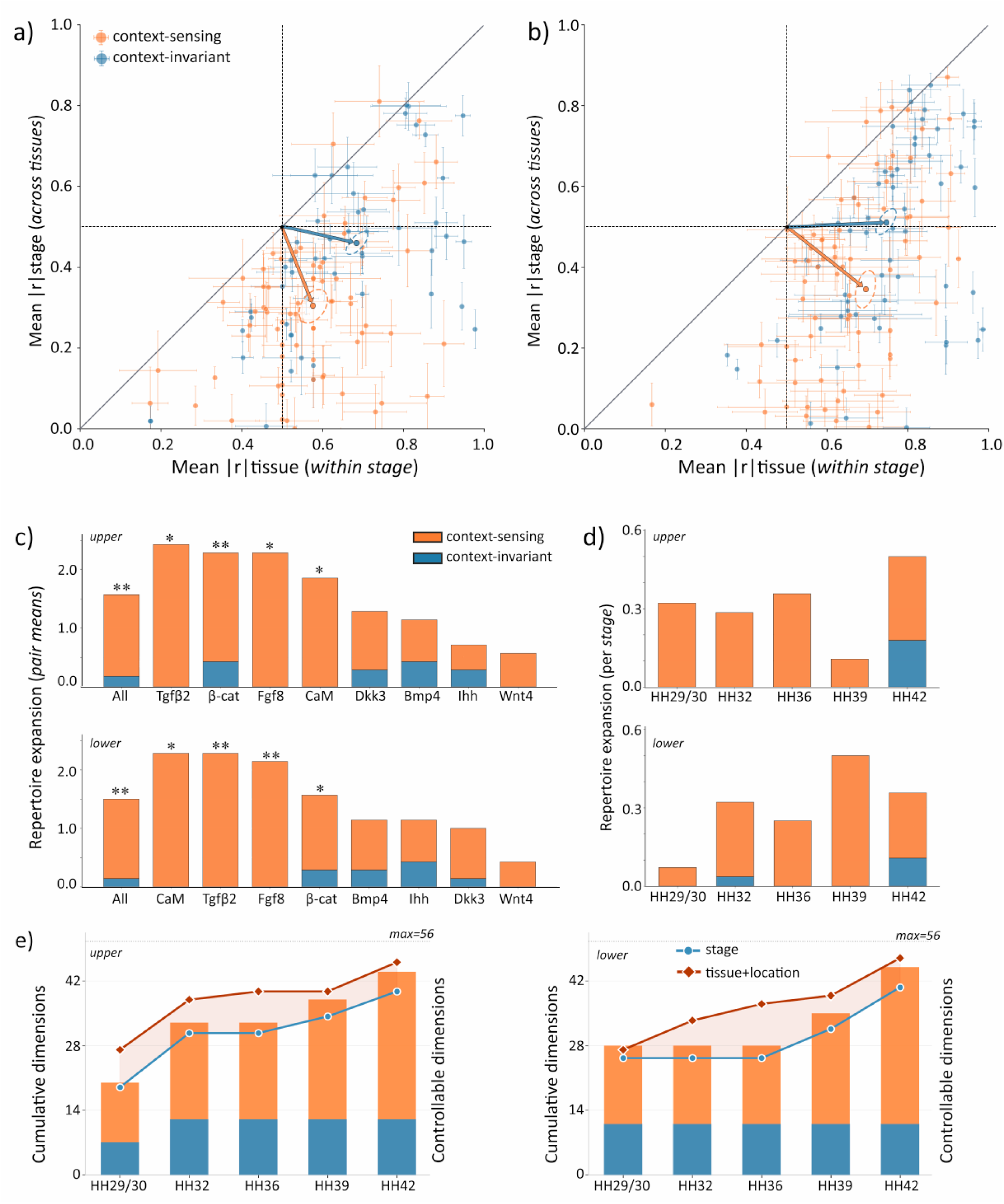
Bow-tie organization expands the network regulatory repertoire. Stage-pooled coexpression masks orthogonal tissue-level subcircuits in **(a)** upper (left column) and **(b)** lower (right column) beak. Context-sensing pairs (orange) cluster in the bottom-right quadrant, where stage-pooled coexpression is weak but tissue coexpression is strong. Context-invariant pairs are shown in blue (Fig. 2a, b). Dashed centroids (mean ±1 s.e.) differ between the two classes (centroid-distance permutation test; upper: *D* = 0.166, *P* < 0.001; lower: *D* = 0.133, *P* = 0.005) and all centroids displaced from midpoint (0.5, 0.5; Hotelling’s T², all *P’s* < 0.05). **(c)** Regulatory repertoire expansion (in regulatory modes averaged across developmental stages, Methods) is driven largely by CaM-, Fgf8-, or Tgfβ2-involving pairs. * denotes values exceeding the random-triad null at *P* < 0.05 (data in Table S5). **(d)** Regulatory repertoire expansion per stage averaged across proteins, for upper and lower beaks. **(e)** Flexible protein colocalization unlocks nearly full regulatory potential. The cumulative (left axis, bars) and controllable (right axis, lines) dimensions contributed at each stage by context-sensing and context-invariant groups of proteins in upper (left) and lower (right) beak. Edge-repertoire size dimensionality is plotted as a proxy of controllability. Lines show total reachable space (in absolute dimensions, out of 56 maximum) under stage-summed inputs (blue circles) versus added by all tissue plus location contexts (red diamonds). The shaded region between the lines is the reachable space added by within-stage contexts’ regulatory states, it is driven largely by context-sensing protein pairs, while context-invariant pairs contribute few non-redundant dimensions across the entire developmental sequence (see Fig. S24).

### Context-sensitivity of hub proteins is associated with repertoire expansion

We quantified the extent to which specific protein pairs expand the network regulatory repertoire (the number of signed correlations between proteins that become masked in the pooled aggregation, see Methods) and found that the network harbors substantially greater regulatory diversity than can be inferred from its stage-summed states (Fig. 3). Repertoire expansion was driven by three proteins: CaM, Tgfβ2, and Fgf8 (hereafter, the triad) that form the hub bridging developmental contexts (Figs. 1c; 4c; S24). In the upper beak, the triad-associated protein pairs gained 3.5 regulatory modes (signed correlations between proteins, summed over stages, see Methods), compared with 0.90 modes for pairs not involving the triad (hereafter, the periphery, *P* < 0.05). In the lower beak, the difference was more pronounced: 6.17 modes/pair vs 0.80 modes/pair (*P* < 0.05). The triad accounted for 63 out of 72 (88%) expansion modes in the upper beak and 111 out of 119 (93%) in the lower beak.

To evaluate whether repertoire expansion corresponds to expansion of the network’s accessible regulatory space, we applied a network controllability framework in which the same set of inputs can drive a network into a larger set of reachable configurations (see Methods). We find that at HH29/30, the network occupied 35.7% (upper) and 50.0% (lower) of theoretical maximum dimensions (here approximated by *n* = 56 reachable signed edges; Fig. S24) and, by HH42, had expanded to 91 (upper) and 93% (lower; Fig. 4e). This expansion occurs without gain or loss of network elements and is driven almost exclusively by context-sensing protein pairs: in both jaws, the flexible subcellular compartmentalization of the triad proteins (CaM, Tgfβ2, Fgf8) and their pairwise interactions contributed the majority of new dimensions at each stage (Figs. 4e and S24). The largest single-stage expansion of controllable space in both jaws, occurred with the simultaneous sign reversals of Bmp4-Tgfβ2 and β-cat-Tgfβ2 (Fig. 2a, b), marking the transition from chondrogenic to osteogenic signaling programs in this species [40]. Together, these results indicate that the repertoire expansion (Fig. 4c, d) and the growth of the network’s reachable space (Fig. 4e) reflect the same process – context-sensing switching enabled by flexible subcellular compartmentalization within an otherwise constant network (Figs. 2c, d; S24).

Interestingly, β-catenin’s contribution is bimodal: its three pairs with the triad proteins averaged 5.0 (upper) and 5.7 (lower) modes/pair, similar to the triad proteins themselves, whereas its four peripheral-partner pairs averaged 0.75 modes/pair in both jaws – also matching the contributions of the periphery pairs. β-catenin therefore behaves as an upstream external input transduced through the triad, consistent with its cellular placement at the convergence of the canonical Wnt and Wnt/Ca²⁺ signaling routes (Fig. 1a).

### Regulatory plasticity despite developmental expansion

We next examine whether the capacity of the regulatory network to flexibly alter its expression (regulatory plasticity) is reduced with developmental expansion (increase in complexity as tissues differentiate), as expected under sequential subsampling, or remains undiminished regardless of how many regulatory modules are added (Fig. 1b *vs* c). We specifically compared variability in the regulatory triad proteins with that in peripheral protein pairs. We find that the triad protein pairs maintained their variability despite developmental expansion (Fig. 5); this organization can forestall regulatory commitment that would otherwise narrow subsequent expansion. In contrast, the plasticity of peripheral linkages remained low but increased with developmental expansion: as more tissues accumulated, these proteins took on increasingly tissue-specific roles (Fig. 5), although the pattern was mostly driven by the upper jaw. Overall, the combination of a flexible waist and a progressively differentiating periphery (Fig. 1c) is the pattern expected from a bow-tie architecture in which the regulatory waist resists commitment while sustaining peripheral specialization.

**Fig 5.**
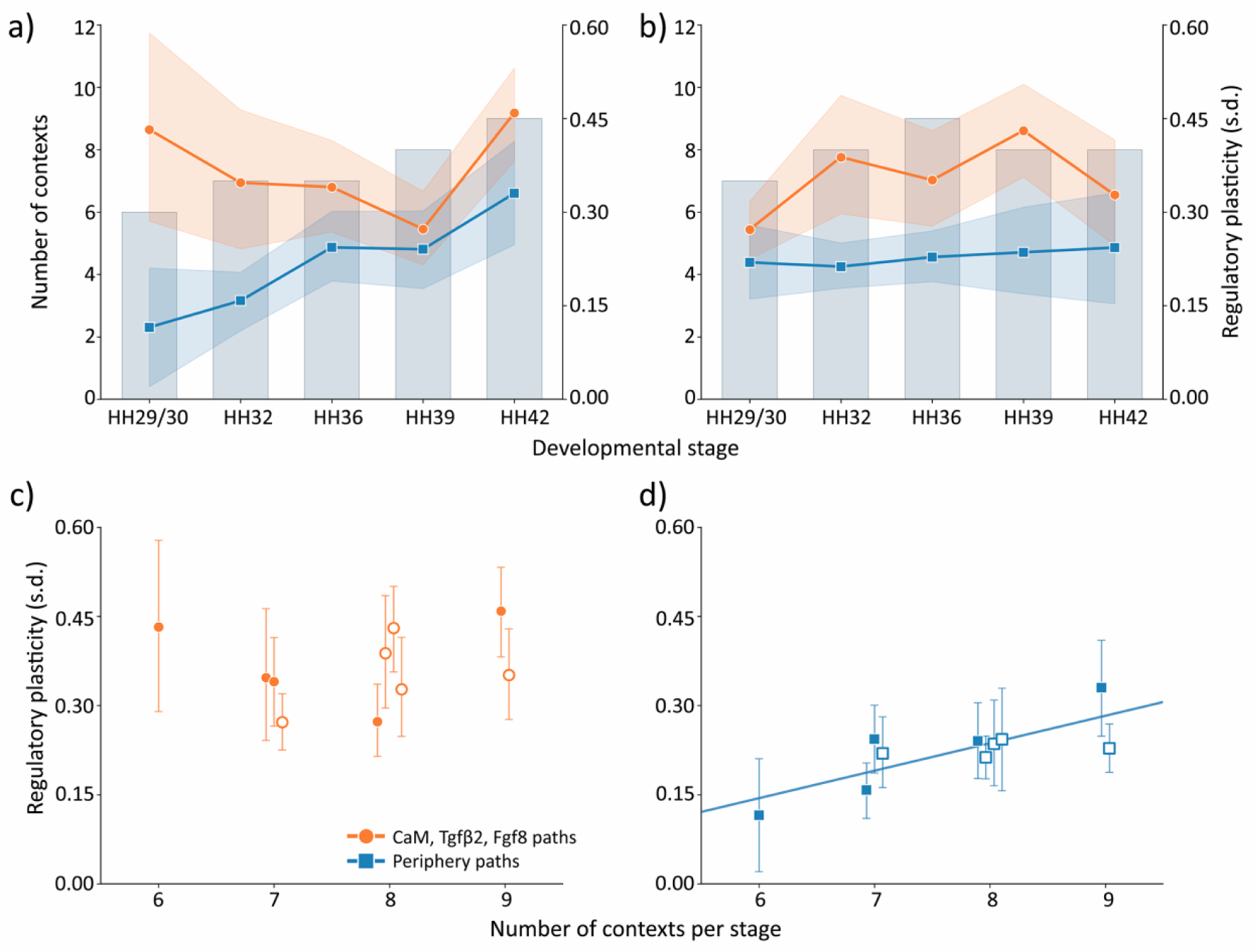
Regulatory plasticity is maintained despite developmental expansion. The number of developmental contexts (tissue types and locations; bars, left axis) across developmental stages for **(a)** upper and **(b)** lower beak and associated variation in regulatory plasticity (across-contexts, within-stage s.d. of Pearson *r’s*; right axis) for proteins in the waist triad of the bow-tie (paths involving CaM, Fgf8, or Tgfβ2, *n* =18 pairs) and proteins at the periphery of the bow-tie organization (paths among β-catenin, Bmp4, Dkk3, Ihh, and Wnt4, *n* =10 pairs). Shaded bands are 95% CIs from bootstrap resampling of the protein group mean (data in Table S6). **(c-d)** Means (±95% CI) of regulatory plasticity in relation to number of within-stage developmental contexts for (c) regulatory waist triad proteins and (d) peripheral proteins. Plasticity of peripheral pairs increased with the number of developmental contexts per stage (*b*_ST_ = 0.81, *P* = 0.004), whereas plasticity of triad-involving pairs showed no trend (*b*_ST_ = 0.01, *P* = 0.99) but was consistently higher (*b*_ST_ = 0.67, P < 0.001; upper beak: closed symbols, lower beak: open symbols; data in Table S7).

## Discussion

Using a newly constructed atlas of avian beak morphogenesis, we find that across multiple developmental contexts in both jaws, a small set of multivalent proteins (CaM, Tgfβ2, and Fgf8) forms a context-sensing hub whose connections to an otherwise stable protein core depend on where in the cell the proteins colocalize. Through this hub, a compact network can regulate multiple tissue-specific states concurrently: it nearly doubles the coexpression repertoire within the same stage and, over development, reaches more than 90% of the signed coexpression states theoretically available to it – without gain, loss, or rewiring of its elements. Although developmental multiplexing in regulatory networks is frequently noted [41–44], its subcellular colocalization basis has not, to our knowledge, been established.

Our results highlight the fundamental advantages of this organization: it would retain undiminished flexibility despite increased complexity (Fig. 5) and is well placed to underpin the robustness of morphogenesis under a wide range of perturbations (Fig. 6). These findings raise two general questions. First, what mechanisms link subcellular distribution and regulatory versatility in the triad proteins? Second, how does the regulatory multiplexing we observe within a stage correspond to repeatability of these states across developmental stages? Can regulatory states prime or bias transitions, thereby allowing within-stage diversification to pull development forward, contributing to its directionality and continuity?

**Fig 6.**
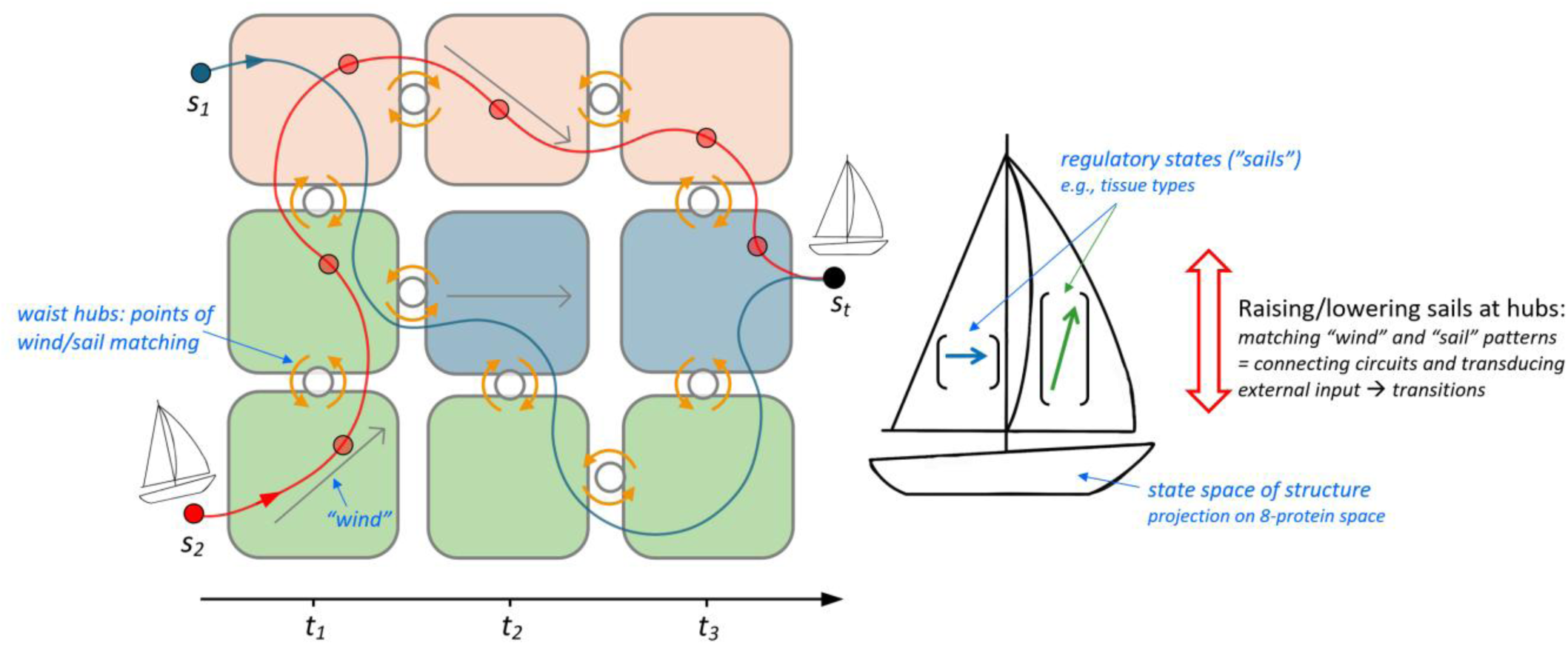
Conceptual illustration of flexible hub proteins reconciling specialization and plasticity on a shared regulatory landscape. Developmental states (e.g., tissue types) are shown as squares of different colors at different temporal stages (x-axis). Red and blue developmental trajectories are temporal projections of a developing structure on the full 8-protein expression state space from distinct starting points, s_1_ and s_2_, toward a common target morphology s_t_. Each tissue/location state has a correlational structure of proteins – shown by vectors within squares. Extending the analogy of [51] we can view these correlations as a background “wind”. Some protein associations are constant across all states and stages (Fig. 4a,b diagonals), providing directionality toward *s*_t_ without prescribing a specific trajectory. We can envision a developing structure as a sailboat with several sails – each harboring a regulatory state, such as tissue-specific protein correlations (Fig. 2a,b). Stages and states (colored squares) are connected by hubs (circles) at which correlational patterns in background wind and those on available sails are compared. They match when sails are “raised”, meaning that a regulatory circuit of a sail is capable of transducing the external input (wind), resulting in state transitions of a developing structure. The mechanism for dynamic raising and lowering of sails is flexibility of subcellular colocalization of proteins (Fig. 2c,d). Thus, dynamic matching at lower levels of organization produces specific and robust outcomes at higher levels despite a fully shared regulatory landscape across all states and stages: dynamic specialization arises without specific regulators. An alternative organization (permanently raised or lowered sails) requires additional regulators to navigate a structure to s_t_ despite unfavorable local background winds. An additional advantage of this organization is that the transduction of external input itself pulls development forward; within-stage diversification and across-stage transitions are produced by the same mechanism.

The first question highlights striking regulatory autonomy of the triad proteins – their capacity to self-generate mechanisms that drive *both* their subcellular distribution and the ability to convert a wide range of non-specific inputs into specific molecular interactions. In each of these proteins, subcellular distribution and regulatory versatility are linked, yet the mechanisms that produce this coupling are distinct. For example, CaM’s distribution among compartments follows a continuous readout of the local calcium environment [45]: changes in substrate stiffness or cell shape alter actomyosin tension, shift calcium gradients [46], and redistribute CaM between the nucleus and cytoplasm [47]. Because CaM’s binding affinity and propensity to form condensates are dosage-dependent [48, 49], this redistribution, in turn, shapes its regulatory interactions [50]. Thus, the entire sequence from physical impact on the cell to locally specific transcriptional output and back to the mechanical state of cell and long-range cell coordination [30], can be routed through CaM without any new protein synthesis, external regulators, or network rewiring.

The other two hub proteins show similar linkage of subcellular distribution and regulatory versatility. Fgf8’s regulatory output and subcellular distribution are tuned by the global ratio of its isoforms [52, 53]; graded, compartment-specific isoform accumulation lets the Fgf8 complex finely adjust its local signaling [54–56]. Tgfβ2, tethered to the extracellular matrix in a latent complex (Fig. 1a), is activated when cytoskeletal tension – itself set by substrate stiffness, cell shape, and contacts with neighboring cells – releases it [57, 58]. Crucially, Tgfβ2 also feeds back on the matrix it senses by upregulating collagen and fibronectin, such that force-dependent release stiffens the matrix and progressively lowers the threshold for subsequent release [59], a self-reinforcing, time-keeping loop that could let the same Tgfβ2 signal yield locally distinct outcomes tuned to tissue mechanics, maturation rate, allometric scaling, and other predictable modulators [60].

Taken together, the diversity of these autoregulatory mechanisms and the breadth of non-specific inputs they accommodate suggest that these proteins are not passive “flexible links” in regulatory networks, but rather encoder-like hubs whose interchangeable physical and biological sensitivities enable them to recognize, compress and translate a wide range of inputs into specific regulatory outputs. Notably, the inputs the hub proteins convert into subcellular circuits come from cell collectives. The common denominator of the hub protein activation is their transduction of cell tension – a collective measure of cell accumulation, uniformity and cooperation [30]. These hubs, therefore, reset local specialization by accessing rules of higher or later levels of organization [e.g., 61], enabling both assembly of and transition between specialized states, ultimately underpinning cell and tissue competence in navigating diverse developmental contexts (Fig. 6).

Because the hub proteins behave as bidirectional switches, the expansion of regulatory repertoire (Fig. 4c,d) does not require additional inputs or topological change – the same upstream signals can be redirected into new regulatory states through colocalization-dependent interactions among hub proteins (Figs. 1c, S24) as is illustrated by the bimodal contribution of β-catenin to the triad and the periphery proteins (Fig. 1a). More generally, this architecture realizes the advantages of temporal controllability of a network: the controllable space of a network is strongly expanded by sequential bidirectional sign reversals at otherwise fixed nodes (Figs. 4e, S24) [51, 62]. Our results identify the flexible subcellular compartmentalization of multivalent hub proteins as a candidate molecular substrate for these sign reversals (Figs. 1a; 2c, d).

The second question concerns how within-stage developmental multiplexing relates to development over time. Regulatory states established within a stage commonly persist into later ones, whether through memory in the network’s switches [63, 64] or through the persistence of physical states and their associated signaling [65, 66]. Such recurrence might facilitate “competence to transition” [67] – the temporal priming of a tissue for subsequent regulation and differentiation. So-called “sensitization” is ubiquitous in development [5, 68, 69] and the bidirectional switches and interchangeability of physical and biological inputs we describe could be one of its mechanisms [70], illustrating how diversification itself can pull development forward by priming sequential transitions (Fig. 6). This is particularly relevant in craniofacial morphogenesis, where diverse outcomes arise from a few conserved mechanisms such as epithelial–mesenchymal interaction [71]: in avian beaks, Shh/Fgf8 boundary signaling presages the frontonasal ectodermal zone [72], and the placement of mesenchymal prominences is primed by regulatory boundaries induced by earlier differentiating epithelium [73].

Our inferences are based on the analyses of protein coexpression, assessed by consensus immunohistochemistry scores and comprehensively measured across numerous contexts. Because the network’s regulatory states depend on where its proteins colocalize, mapping which interactions are context-sensing and which are context-invariant is a useful step before experimental manipulation: perturbations that do not account for this structure may shift the network into an alternative configuration rather than isolate a causal mechanism. Our developmental atlas supplies that map, identifying targets of experimental manipulation. For example, the mechanisms we invoke for the three hub proteins under this study – calcium-gated relocation of CaM, isoform partitioning of Fgf8, and tension-based activation of Tgfβ2 – identify the subcellular localization as one such target: its perturbation in any hub protein should collapse within-stage regulatory diversity without altering network topology.

If confirmed experimentally, an architecture in which a few dynamic, self-organizing encoder-like hubs convert diverse inputs may help explain how development reaches a target morphology across a wide range of conditions (Fig. 6) and thus provide a mechanistic framework that could reconcile the ecological precision and the evolutionary lability evident in avian beak diversification.

## Materials and methods

### Data collection and sample sizes

We measured protein expression and its subcellular, histological and anatomical distribution in upper and lower beaks of *n* = 322 embryos house finches (*Haemorhous mexicanus*) across five Hamburger-Hamilton (HH) developmental stages (HH29/30: *n* = 62, HH32: *n* = 122; HH36: *n* = 47; HH39: *n* = 53, and HH42: *n* = 38). Protocols for egg collection, field storage of samples, clutch randomization, and incubation to the required developmental stage are in [30]. Briefly, upper and lower beaks were cryosectioned at midline at 8 µm and stored at −80°C. Thirteen sections per individual were obtained at beak midline: one section was stained with Alcian blue hematoxylin and eosin (H&E) to delineate the histological area of interest and twelve were used in immunohistochemical (IHC) analyses (see below for combination with negative and isotype controls). Only complete, undamaged midline sections of both jaws were used to construct a developmental atlas for this work (Figs. S1-21). Upper and lower beaks were divided into three equal anterior-posterior spans [74] (hereafter, locations): base, mid, and tip; tongue was recorded as a separate location in the lower beak (Tables S1-2).

### Immunohistochemistry assays of protein expression

We used IHC to measure expression of core craniofacial proteins: β-catenin, bone morphogenetic protein 4 (Bmp4), calmodulin 1 (CaM), dickkopf homolog 3 (Dkk3), fibroblast growth factor 8 (Fgf8), Indian hedgehog (Ihh), transforming growth factor beta 2 (Tgfβ2), and Wnt family member 4 (Wnt4). Summary of involvement of these proteins in major developmental pathways (TGF-β, Wnt, FGF, Hedgehog, and calcium signaling) is in [39] and Fig 1a. For immunostaining, we used anti-β-catenin (610153, 1:16,000, BD Transduction Laboratories), anti-CaM (sc-137079, 1:15, Santa Cruz Biotechnology), anti-Wnt4 (ab91226, 1:800; Abcam), anti-Tgfβ2 (ab36495, 1:800, Abcam), anti-Bmp4 (ab118867, 1:100, Abcam), anti-Ihh (ab184624, 1:100, Abcam), anti-Dkk3 (ab214360, 1:100, Abcam), and anti-Fgf8 (ab89550, 1:50, Abcam) antibodies. Reactions were visualized with either diaminobenzidine (DAB, Elite ABC HRP Kit, PK-6100, Vector Labs) or Vector Red Alkaline Phosphatase substrate and Vectastain ABC-AP Kit (AK-5000, Vector Labs) and nuclei were counterstained with hematoxylin. Protein expression intensity was recorded for seven tissues (Tables S1-S2) and scored within each section as shown in Fig. S25b by observers blind to the hypotheses of this study. Automated scoring [74] and manual scoring produced identical results in this dataset, however, because in this study we were particularly interested in confirmation of subcellular locations of proteins that our staining allowed, manual scoring with two addition confirmation steps was used. First, comparison of protein intensity among regions of the beak was conducted within the same section of an individual to ensure that staining variability was not recorded as expression difference. Second, the protein expression patterns were further evaluated for consistency across individuals. Low magnifications (4× and 10×) were used to confirm locations and consistency of expression across the beak.

We classified subcellular compartments of proteins under high magnification (20× and 40×; Fig. S25 a,b) as nuclear if protein expression overlapped with the stained nucleus, perinuclear if protein expression was lining the nucleus, cytoplasmic if protein expression was in cytoplasm, or membranal if protein binding was expressed in the cell membrane. Our assessment of subcellular location of protein (Fig. S25a, b) corroborates previously reported rare colocalization of some proteins. For example, we report perinuclear expression of Tgfβ2; this pattern is consistent with its secretion to the ECM that is routed through the ER/Golgi [39, 60, 75]. Endogenous Fgf8 was also reported in the nucleus of mouse embryonic neural tube cells [76, 77]. Similarly, nuclear expression of Fgf8 in chondrogenic regions [40] is consistent with its role in chondrocyte differentiation and osteogenic gene downregulation [77, 78]. Protocols, validation assays, verification of isotype controls for avian specificity of stains, and interobserver repeatability are given in [30, 39, 74, 79], respectively.

### Statistical analysis

#### Protein correlations

To quantify protein coexpression across locations and tissue types for each developmental stage and separately for upper and lower beak, we recoded protein expression (Tables S1-2, Figs. S25-S26) as integers (0 = absent, 1 = low, 2 = medium, 3 = high); all four levels were retained in the correlation analysis. Excluding absent values did not change the context-sensing/context-invariant classification (below). We calculated Pearson *r* for all 28 protein pairs across all contexts. Means across contexts were taken on Fisher *z-*transformed values [80] to correct for skew of correlations near ±1, then back-transformed for presentation (Table S3). For each protein pair, overall variation (Fig. 2c, d) was quantified as the Bessel-corrected (*n* −1) standard deviation (s.d.) of the Fisher *z-*transformed correlations across all stages and contexts. Group-level s.d. were estimated with 95% CI from B = 2,000 within-group resampling protein pairs with replacement. Because Pearson *r* and Fisher *z* transformations are sensitive to the assumption of the continuous distribution, we repeated all analyses with Spearman ρ and Kendall τ; the classification of pairs as context-sensing or context-invariant did not change across all three methods.

#### Classification of protein pairs

A pair was classified as context-sensing (sign-switching) if the sign of any correlation switched in any context; and classified as context-invariant (sign-constant) if otherwise. We used the permutation null model to establish significance of context-specific correlations. Each pair was also summarized by an index, *I* = max[0, min(*r*max −rmin)], where rmax and rmin are the largest and smallest context-specific *r*’s. Observed *I* was compared with 2,000 datasets in which tissue and location labels were permuted among records within each stage, leaving the stage-pooled data unchanged. A pair was classified as context-sensing if its observed *I* exceeded 95% of its null values.

#### Repertoire expansion and regulatory dimensionality

Repertoire expansion was the number of signed correlations between proteins (*regulatory modes* hereafter) that were significant within a developmental context (e.g., tissue type) but become nonsignificant (masked) in the stage-pooled expression. In this metric, each protein pair has four possible regulatory modes in each stage-specific repertoire: two in each tissue type and two in each location on each jaw. Because this metric does not account for the magnitude of switches, it conservatively estimates the lowest bound of repertoire expansion. We followed [81] to test this metric against two null models to ensure that the observed repertoire expansion exceeds that expected from sampling variance alone and does not arise from aggregation (Simpson’s paradox). First, for each developmental stage, the joint association between contexts and protein expression vectors was reshuffled, such that each (stage; context) array in the shuffled dataset became a uniform random draw from the homogeneous within-stage pool. Repertoire expansion was then recomputed on the reshuffled dataset, yielding the per-pair expansion expected under the null hypothesis that within-stage regulatory structure is homogeneous and that all observed expansion is sampling variance from tissue or location contexts. We report the observed per-pair mean against B = 500 random resamplings per jaw. Second, we test whether CaM-, Tgfβ2-, and Fgf8-involved pairs harbor expansion above what is expected for a randomly chosen three-protein subset of the network (Figs. 4 and 5). The test statistic was the difference in mean per-pair expansion between pairs involving at least one triad protein (*n* = 18) and the remaining pairs (*n* = 10). Since 56 triads are possible, we retained those with *P* < 0.02 (1/56).

Motivated by work on the controllability of temporal networks [51], in which sign reversals of fixed links enlarge the set of reachable configurations without any change in network composition, we summarized the range of expressed coexpression states as signed-edge coverage -- a combinatorial proxy of a controllability measure. This is the cumulative number of distinct signed pairwise associations (max=56 (28×2)) that were significant (*P* < 0.05) in at least one context at or before a given stage (Fig. S24). Reachable space was computed from stage-pooled correlations alone and from all within-stage tissue and location contexts; the difference is added by context-specific coexpression (Fig. 4e).

**Supplementary Materials:** Figures S1 to S26; Tables S1 to S7.

## Supplementary information

This file includes:

1) **Supplementary Table S1:** Hyperlinked summary of protein expression across developmental stages, axial locations, and subcellular contexts for the upper beak.

2) **Supplementary Table S2:** Hyperlinked summary of protein expression across developmental stages, axial locations, and subcellular contexts for the lower beak

3) **Supplementary Table S3:** Context-specific Fisher’s Z-normalized Pearson r’s (data for Fig. 2)

4) **Supplementary Table S4:** Protein-pair tissue switches associated with subcellular relocation or compartment changes

5) **Supplementary Table S5:** Stage vs tissue |r’s| and associated statistics (data for Fig. 4)

6) **Supplementary Table S6:** Raw data for regulatory plasticity and developmental expansion figure (data for Fig. 5)

7) **Supplementary Table S7:** Bootstrap intervals for Fig. 5

8) **Supplementary Figure S1-S21:** Complete developmental atlas of house finch beak morphogenesis (Supplementary Tables S1, S2)

9) **Supplementary Figure S22:** Non-overlapping tissue-specific regulatory states of the core network

10) **Supplementary Figure S23:** Non-overlapping location-specific regulatory states of the core network

11) **Supplementary Figure S24:** Context-sensing proteins drive non-redundant controllable space expansion

12) **Supplementary Figure S25:** Examples of protein categorization to different subcellular locations

13) **Supplementary Figure S26:** Example illustrating protein expression intensity measures

*Tables S3 and S5-7 are included in Table S3-7.zip

---

**TABLE S1.**
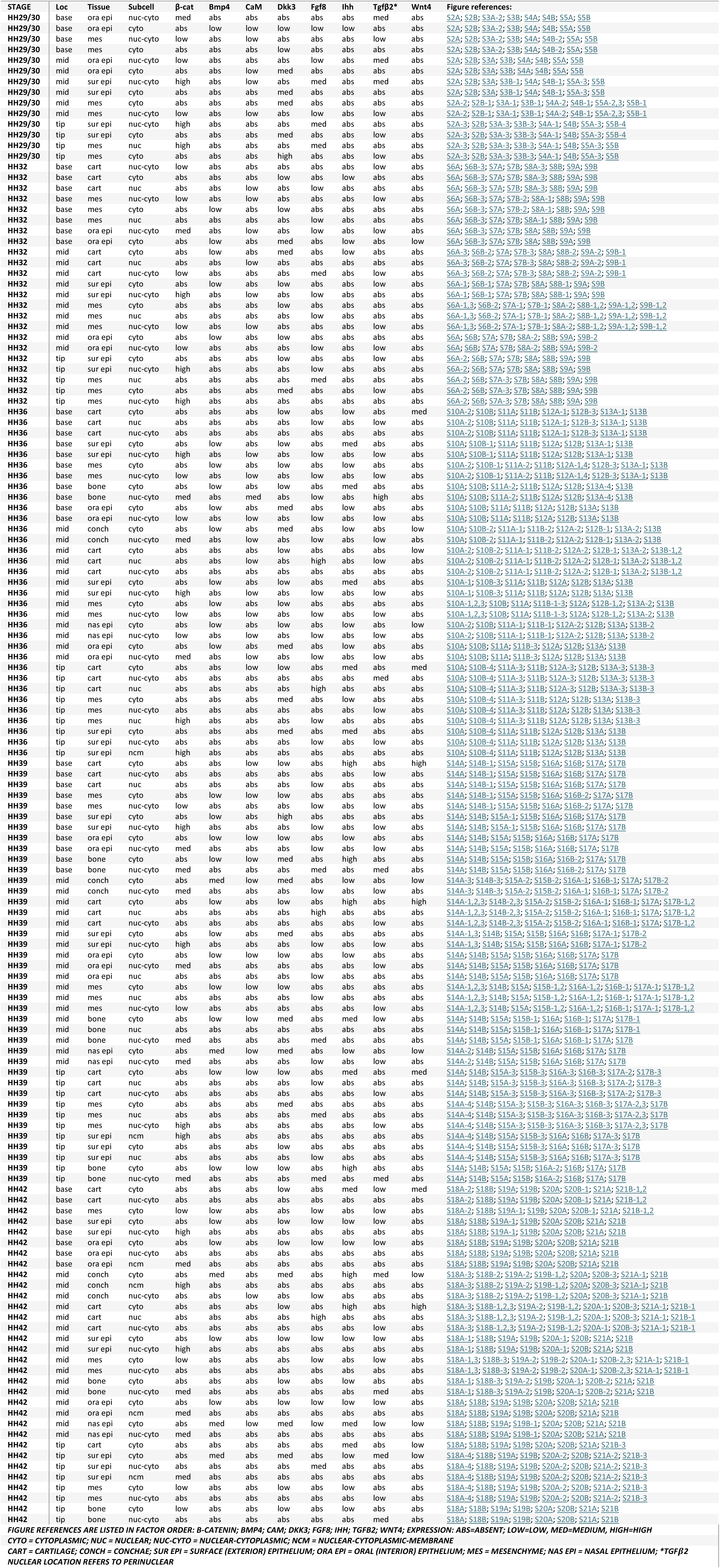
SUMMARY OF PROTEIN EXPRESSION ACROSS DEVELOPMENTAL STAGES, AXIAL LOCATIONS, AND SUBCELLULAR CONTEXTS FOR UPPER BEAK.

| **STAGE** | **Loc** | **Tissue** | **Subcell** | **β-cat** | **Bmp4** | **CaM** | **Dkk3** | **Fgf8** | **Ihh** | **Tgfβ2\*** | **Wnt4** | **Figure references:** |
|---|---|---|---|---|---|---|---|---|---|---|---|---|
| **HH29/30** | base | ora epi | nuc-cyto | med | abs | abs | abs | abs | abs | med | abs | S2A; S2B; S3A-2; S3B; S4A; S4B; S5A; S5B |
| **HH29/30** | base | ora epi | cyto | abs | low | abs | low | low | low | abs | abs | S2A; S2B; S3A-2; S3B; S4A; S4B; S5A; S5B |
| **HH29/30** | base | mes | nuc-cyto | low | abs | low | abs | abs | abs | abs | abs | S2A; S2B; S3A-2; S3B; S4A-2; S4B-2; S5A; S5B |
| **HH29/30** | base | mes | cyto | abs | abs | abs | low | abs | abs | abs | abs | S2A; S2B; S3A-2; S3B; S4A; S4B-2; S5A; S5B |
| **HH29/30** | mid | ora epi | nuc-cyto | low | abs | low | abs | abs | abs | med | abs | S2A; S2B; S3A; S3B; S4A; S4B; S5A; S5B |
| **HH29/30** | mid | ora epi | cyto | abs | abs | low | med | abs | abs | abs | abs | S2A; S2B; S3A; S3B; S4A; S4B; S5A; S5B |
| **HH29/30** | mid | sur epi | nuc-cyto | high | abs | low | abs | med | abs | med | abs | S2A; S2B; S3A; S3B-1; S4A; S4B-1; S5A-3; S5B |
| **HH29/30** | mid | sur epi | cyto | abs | abs | abs | med | abs | abs | abs | abs | S2A; S2B; S3A; S3B-1; S4A; S4B-1; S5A-3; S5B |
| **HH29/30** | mid | mes | cyto | abs | abs | abs | med | abs | abs | abs | abs | S2A-2; S2B-1; S3A-1; S3B-1; S4A-2; S4B-1; S5A-2,3; S5B-1 |
| **HH29/30** | mid | mes | nuc-cyto | low | abs | low | abs | low | abs | abs | low | S2A-2; S2B-1; S3A-1; S3B-1; S4A-2; S4B-1; S5A-2,3; S5B-1 |
| **HH29/30** | tip | sur epi | nuc-cyto | high | abs | low | abs | med | abs | abs | abs | S2A-3; S2B; S3A-3; S3B-3; S4A-1; S4B; S5A-3; S5B-4 |
| **HH29/30** | tip | sur epi | cyto | abs | abs | abs | med | abs | abs | abs | low | S2A-3; S2B; S3A-3; S3B-3; S4A-1; S4B; S5A-3; S5B-4 |
| **HH29/30** | tip | mes | nuc | high | abs | abs | abs | med | abs | abs | abs | S2A-3; S2B; S3A-3; S3B-3; S4A-1; S4B; S5A-3; S5B |
| **HH29/30** | tip | mes | cyto | abs | abs | abs | high | abs | abs | abs | abs | S2A-3; S2B; S3A-3; S3B-3; S4A-1; S4B; S5A-3; S5B |
| **HH32** | base | cart | nuc-cyto | low | abs | abs | abs | abs | abs | abs | low | S6A; S6B-3; S7A; S7B; S8A-3; S8B; S9A; S9B |
| **HH32** | base | cart | cyto | abs | abs | abs | low | abs | low | abs | abs | S6A; S6B-3; S7A; S7B; S8A-3; S8B; S9A; S9B |
| **HH32** | base | cart | nuc | abs | abs | low | abs | low | abs | abs | abs | S6A; S6B-3; S7A; S7B; S8A-3; S8B; S9A; S9B |
| **HH32** | base | mes | nuc-cyto | low | abs | abs | abs | low | abs | abs | low | S6A; S6B-3; S7A; S7B-2; S8A-1; S8B; S9A; S9B |
| **HH32** | base | mes | cyto | abs | low | abs | low | low | abs | abs | abs | S6A; S6B-3; S7A; S7B; S8A-1; S8B; S9A; S9B |
| **HH32** | base | mes | nuc | abs | abs | abs | abs | low | abs | abs | abs | S6A; S6B-3; S7A; S7B; S8A-1; S8B; S9A; S9B |
| **HH32** | base | ora epi | nuc-cyto | med | abs | low | abs | low | abs | abs | abs | S6A; S6B-3; S7A; S7B; S8A; S8B; S9A; S9B |
| **HH32** | base | ora epi | cyto | abs | low | abs | abs | abs | abs | abs | low | S6A; S6B-3; S7A; S7B; S8A; S8B; S9A; S9B |
| **HH32** | mid | cart | cyto | abs | abs | abs | med | abs | low | abs | abs | S6A-3; S6B-2; S7A; S7B-3; S8A; S8B-2; S9A-2; S9B-1 |
| **HH32** | mid | cart | nuc | abs | abs | low | abs | abs | abs | abs | abs | S6A-3; S6B-2; S7A; S7B-3; S8A; S8B-2; S9A-2; S9B-1 |
| **HH32** | mid | cart | nuc-cyto | low | abs | abs | abs | med | abs | abs | low | S6A-3; S6B-2; S7A; S7B-3; S8A; S8B-2; S9A-2; S9B-1 |
| **HH32** | mid | sur epi | cyto | abs | abs | abs | abs | abs | abs | abs | low | S6A-1; S6B-1; S7A; S7B; S8A; S8B-1; S9A; S9B |
| **HH32** | mid | sur epi | nuc-cyto | high | abs | abs | abs | low | abs | abs | low | S6A-1; S6B-1; S7A; S7B; S8A; S8B-1; S9A; S9B |
| **HH32** | mid | mes | cyto | abs | abs | abs | low | abs | abs | abs | abs | S6A-1,3; S6B-2; S7A-1; S7B-1; S8A-2; S8B-1,2; S9A-1,2; S9B-1,2 |
| **HH32** | mid | mes | nuc | abs | abs | abs | abs | low | abs | abs | abs | S6A-1,3; S6B-2; S7A-1; S7B-1; S8A-2; S8B-1,2; S9A-1,2; S9B-1,2 |
| **HH32** | mid | mes | nuc-cyto | low | abs | low | abs | low | abs | abs | low | S6A-1,3; S6B-2; S7A-1; S7B-1; S8A-2; S8B-1,2; S9A-1,2; S9B-1,2 |
| **HH32** | mid | ora epi | cyto | abs | low | abs | low | abs | abs | abs | abs | S6A; S6B; S7A; S7B; S8A-2; S8B; S9A; S9B-2 |
| **HH32** | mid | ora epi | nuc-cyto | low | abs | abs | abs | low | abs | abs | abs | S6A; S6B; S7A; S7B; S8A-2; S8B; S9A; S9B-2 |
| **HH32** | tip | sur epi | cyto | abs | abs | abs | low | abs | abs | abs | low | S6A-2; S6B; S7A; S7B; S8A; S8B; S9A; S9B |
| **HH32** | tip | sur epi | nuc-cyto | high | abs | abs | abs | low | abs | abs | low | S6A-2; S6B; S7A; S7B; S8A; S8B; S9A; S9B |
| **HH32** | tip | mes | nuc | abs | abs | abs | abs | med | abs | abs | abs | S6A-2; S6B; S7A-3; S7B; S8A; S8B; S9A; S9B |
| **HH32** | tip | mes | cyto | abs | abs | abs | med | abs | abs | abs | abs | S6A-2; S6B; S7A-3; S7B; S8A; S8B; S9A; S9B |
| **HH32** | tip | mes | nuc-cyto | high | abs | abs | abs | abs | abs | abs | abs | S6A-2; S6B; S7A-3; S7B; S8A; S8B; S9A; S9B |
| **HH36** | base | cart | cyto | abs | abs | abs | low | abs | low | abs | med | S10A-2; S10B; S11A; S11B; S12A-1; S12B-3; S13A-1; S13B |
| **HH36** | base | cart | nuc | abs | abs | abs | abs | low | abs | abs | low | S10A-2; S10B; S11A; S11B; S12A-1; S12B-3; S13A-1; S13B |
| **HH36** | base | cart | nuc-cyto | low | abs | abs | abs | abs | abs | abs | abs | S10A-2; S10B; S11A; S11B; S12A-1; S12B-3; S13A-1; S13B |
| **HH36** | base | sur epi | cyto | abs | low | abs | low | abs | med | abs | abs | S10A; S10B-1; S11A; S11B; S12A; S12B; S13A-1; S13B |
| **HH36** | base | sur epi | nuc-cyto | high | abs | low | abs | low | abs | abs | low | S10A; S10B-1; S11A; S11B; S12A; S12B; S13A-1; S13B |
| **HH36** | base | mes | cyto | abs | low | abs | low | low | abs | abs | abs | S10A-2; S10B-1; S11A-2; S11B; S12A-1,4; S12B-3; S13A-1; S13B |
| **HH36** | base | mes | nuc-cyto | low | abs | abs | abs | low | abs | abs | low | S10A-2; S10B-1; S11A-2; S11B; S12A-1,4; S12B-3; S13A-1; S13B |
| **HH36** | base | bone | cyto | abs | low | abs | low | abs | med | abs | abs | S10A; S10B; S11A-2; S11B; S12A; S12B; S13A-4; S13B |
| **HH36** | base | bone | nuc-cyto | med | abs | med | abs | abs | abs | high | abs | S10A; S10B; S11A-2; S11B; S12A; S12B; S13A-4; S13B |
| **HH36** | base | ora epi | cyto | abs | low | abs | med | abs | low | abs | abs | S10A; S10B; S11A; S11B; S12A; S12B; S13A; S13B |
| **HH36** | base | ora epi | nuc-cyto | low | abs | low | abs | low | abs | abs | abs | S10A; S10B; S11A; S11B; S12A; S12B; S13A; S13B |
| **HH36** | mid | conch | cyto | abs | low | abs | med | abs | low | abs | low | S10A; S10B-2; S11A-1; S11B-2; S12A-1; S12B-1; S13A-2; S13B |
| **HH36** | mid | conch | nuc-cyto | med | abs | low | abs | low | abs | abs | low | S10A; S10B-2; S11A-1; S11B-2; S12A-2; S12B-1; S13A-2; S13B |
| **HH36** | mid | cart | cyto | abs | abs | abs | low | abs | low | abs | low | S10A-2; S10B-2; S11A-1; S11B-2; S12A-2; S12B-1; S13A-2; S13B-1,2 |
| **HH36** | mid | cart | nuc | abs | abs | low | abs | high | abs | abs | abs | S10A-2; S10B-2; S11A-1; S11B-2; S12A-2; S12B-1; S13A-2; S13B-1,2 |
| **HH36** | mid | cart | nuc-cyto | low | abs | abs | abs | abs | abs | abs | abs | S10A-2; S10B-2; S11A-1; S11B-2; S12A-2; S12B-1; S13A-2; S13B-1,2 |
| **HH36** | mid | sur epi | cyto | abs | low | abs | low | abs | med | abs | abs | S10A-1; S10B-3; S11A; S11B; S12A; S12B; S13A; S13B |
| **HH36** | mid | sur epi | nuc-cyto | high | abs | low | abs | low | abs | abs | low | S10A-1; S10B-3; S11A; S11B; S12A; S12B; S13A; S13B |
| **HH36** | mid | mes | cyto | abs | low | abs | low | low | abs | abs | abs | S10A-1,2,3; S10B; S11A; S11B-1-3; S12A; S12B-1,2; S13A-2; S13B |
| **HH36** | mid | mes | nuc-cyto | low | abs | abs | abs | low | abs | abs | low | S10A-1,2,3; S10B; S11A; S11B-1-3; S12A; S12B-1,2; S13A-2; S13B |
| **HH36** | mid | nas epi | cyto | abs | low | abs | low | abs | low | abs | low | S10A-2; S10B; S11A-1; S11B-1; S12A-2; S12B; S13A; S13B-2 |
| **HH36** | mid | nas epi | nuc-cyto | low | abs | abs | abs | abs | abs | abs | abs | S10A-2; S10B; S11A-1; S11B-1; S12A-2; S12B; S13A; S13B-2 |
| **HH36** | mid | ora epi | cyto | abs | low | abs | med | abs | low | abs | abs | S10A; S10B; S11A; S11B-3; S12A; S12B; S13A; S13B |
| **HH36** | mid | ora epi | nuc-cyto | med | abs | low | abs | low | abs | abs | abs | S10A; S10B; S11A; S11B-3; S12A; S12B; S13A; S13B |
| **HH36** | tip | cart | cyto | abs | abs | abs | low | abs | med | abs | med | S10A; S10B-4; S11A-3; S11B; S12A-3; S12B; S13A-3; S13B-3 |
| **HH36** | tip | cart | nuc-cyto | low | abs | abs | abs | abs | abs | med | abs | S10A; S10B-4; S11A-3; S11B; S12A-3; S12B; S13A-3; S13B-3 |
| **HH36** | tip | cart | nuc | abs | abs | abs | abs | high | abs | abs | abs | S10A; S10B-4; S11A-3; S11B; S12A-3; S12B; S13A-3; S13B-3 |
| **HH36** | tip | mes | cyto | abs | abs | abs | med | abs | low | abs | abs | S10A; S10B-4; S11A-3; S11B; S12A; S12B; S13A; S13B-3 |
| **HH36** | tip | mes | nuc-cyto | low | abs | abs | abs | low | abs | abs | low | S10A; S10B-4; S11A-3; S11B; S12A; S12B; S13A; S13B-3 |
| **HH36** | tip | mes | nuc | high | abs | abs | abs | low | abs | abs | abs | S10A; S10B-4; S11A-3; S11B; S12A; S12B; S13A-3; S13B-3 |
| **HH36** | tip | sur epi | cyto | abs | abs | abs | med | abs | med | abs | abs | S10A; S10B-4; S11A; S11B; S12A; S12B; S13A; S13B |
| **HH36** | tip | sur epi | nuc-cyto | low | abs | abs | abs | low | abs | abs | abs | S10A; S10B-4; S11A; S11B; S12A; S12B; S13A; S13B |
| **HH36** | tip | sur epi | ncm | high | abs | abs | abs | low | abs | abs | abs | S10A; S10B-4; S11A; S11B; S12A; S12B; S13A; S13B |
| **HH39** | base | cart | cyto | abs | abs | abs | low | low | high | abs | high | S14A; S14B-1; S15A; S15B; S16A; S16B; S17A; S17B |
| **HH39** | base | cart | nuc-cyto | low | abs | abs | abs | abs | abs | abs | low | S14A; S14B-1; S15A; S15B; S16A; S16B; S17A; S17B |
| **HH39** | base | cart | nuc | abs | abs | abs | abs | low | abs | abs | abs | S14A; S14B-1; S15A; S15B; S16A; S16B; S17A; S17B |
| **HH39** | base | mes | cyto | abs | low | abs | low | low | abs | abs | abs | S14A; S14B-1; S15A; S15B; S16A; S16B-2; S17A; S17B |
| **HH39** | base | mes | nuc-cyto | low | abs | abs | abs | low | abs | abs | low | S14A; S14B-1; S15A; S15B; S16A; S16B-2; S17A; S17B |
| **HH39** | base | sur epi | cyto | abs | low | abs | high | abs | abs | abs | abs | S14A; S14B; S15A-1; S15B; S16A; S16B; S17A; S17B |
| **HH39** | base | sur epi | nuc-cyto | high | abs | abs | abs | low | abs | abs | low | S14A; S14B; S15A-1; S15B; S16A; S16B; S17A; S17B |
| **HH39** | base | ora epi | cyto | abs | low | abs | low | abs | abs | abs | abs | S14A; S14B; S15A; S15B; S16A; S16B; S17A; S17B |
| **HH39** | base | ora epi | nuc-cyto | med | abs | abs | abs | low | abs | abs | abs | S14A; S14B; S15A; S15B; S16A; S16B; S17A; S17B |
| **HH39** | base | bone | cyto | abs | low | abs | low | med | high | abs | abs | S14A; S14B; S15A; S15B; S16A; S16B-2; S17A; S17B |
| **HH39** | base | bone | nuc-cyto | med | abs | abs | abs | med | abs | med | abs | S14A; S14B; S15A; S15B; S16A; S16B-2; S17A; S17B |
| **HH39** | mid | conch | cyto | abs | med | abs | med | abs | low | abs | low | S14A-3; S14B-3; S15A-2; S15B-2; S16A-1; S16B-1; S17A; S17B-2 |
| **HH39** | mid | conch | nuc-cyto | med | abs | abs | abs | low | abs | abs | low | S14A-3; S14B-3; S15A-2; S15B-2; S16A-1; S16B-1; S17A; S17B-2 |
| **HH39** | mid | cart | cyto | abs | abs | abs | low | abs | high | abs | high | S14A-1,2,3; S14B-2,3; S15A-2; S15B-2; S16A-1; S16B-1; S17A; S17B-1,2 |
| **HH39** | mid | cart | nuc | abs | abs | abs | abs | high | abs | abs | abs | S14A-1,2,3; S14B-2,3; S15A-2; S15B-2; S16A-1; S16B-1; S17A; S17B-1,2 |
| **HH39** | mid | cart | nuc-cyto | low | abs | abs | abs | abs | abs | abs | low | S14A-1,2,3; S14B-2,3; S15A-2; S15B-2; S16A-1; S16B-1; S17A; S17B-1,2 |
| **HH39** | mid | sur epi | cyto | abs | low | abs | med | abs | abs | abs | abs | S14A-1,3; S14B; S15A; S15B; S16A; S16B; S17A-1; S17B-2 |
| **HH39** | mid | sur epi | nuc-cyto | high | abs | abs | abs | low | abs | abs | low | S14A-1,3; S14B; S15A; S15B; S16A; S16B; S17A-1; S17B-2 |
| **HH39** | mid | ora epi | cyto | abs | low | abs | low | abs | abs | abs | abs | S14A; S14B; S15A; S15B; S16A; S16B; S17A; S17B |
| **HH39** | mid | ora epi | nuc-cyto | med | abs | abs | abs | low | abs | abs | abs | S14A; S14B; S15A; S15B; S16A; S16B; S17A; S17B |
| **HH39** | mid | ora epi | nuc | abs | abs | abs | abs | low | abs | abs | abs | S14A; S14B; S15A; S15B; S16A; S16B; S17A; S17B |
| **HH39** | mid | mes | cyto | abs | low | abs | low | low | abs | abs | abs | S14A-1,2,3; S14B; S15A; S15B-1,2; S16A-1,2; S16B-1; S17A-1; S17B-1,2 |
| **HH39** | mid | mes | nuc | abs | abs | abs | abs | low | abs | abs | abs | S14A-1,2,3; S14B; S15A; S15B-1,2; S16A-1,2; S16B-1; S17A-1; S17B-1,2 |
| **HH39** | mid | mes | nuc-cyto | low | abs | abs | abs | low | abs | abs | low | S14A-1,2,3; S14B; S15A; S15B-1,2; S16A-1,2; S16B-1; S17A-1; S17B-1,2 |
| **HH39** | mid | bone | cyto | abs | low | abs | low | med | abs | med | low | S14A; S14B; S15A; S15B-1; S16A; S16B-1; S17A; S17B-1 |
| **HH39** | mid | bone | nuc | abs | abs | abs | abs | low | abs | abs | abs | S14A; S14B; S15A; S15B-1; S16A; S16B-1; S17A; S17B-1 |
| **HH39** | mid | bone | nuc-cyto | med | abs | abs | abs | med | abs | abs | abs | S14A; S14B; S15A; S15B-1; S16A; S16B-1; S17A; S17B |
| **HH39** | mid | nas epi | cyto | abs | med | low | med | abs | low | abs | low | S14A-2; S14B; S15A; S15B; S16A; S16B; S17A; S17B |
| **HH39** | mid | nas epi | nuc-cyto | med | abs | abs | abs | low | abs | abs | low | S14A-2; S14B; S15A; S15B; S16A; S16B; S17A; S17B |
| **HH39** | tip | cart | cyto | abs | low | abs | low | abs | med | abs | med | S14A; S14B; S15A-3; S15B-3; S16A-3; S16B-3; S17A-2; S17B-3 |
| **HH39** | tip | cart | nuc | abs | abs | abs | abs | low | abs | abs | abs | S14A; S14B; S15A-3; S15B-3; S16A-3; S16B-3; S17A-2; S17B-3 |
| **HH39** | tip | cart | nuc-cyto | low | abs | abs | abs | abs | abs | abs | low | S14A; S14B; S15A-3; S15B-3; S16A-3; S16B-3; S17A-2; S17B-3 |
| **HH39** | tip | mes | cyto | abs | abs | abs | med | abs | abs | abs | abs | S14A-4; S14B; S15A-3; S15B-3; S16A-3; S16B-3; S17A-2,3; S17B |
| **HH39** | tip | mes | nuc | abs | abs | abs | abs | med | abs | abs | abs | S14A-4; S14B; S15A-3; S15B-3; S16A-3; S16B-3; S17A-2,3; S17B |
| **HH39** | tip | mes | nuc-cyto | high | abs | abs | abs | low | abs | abs | low | S14A-4; S14B; S15A-3; S15B-3; S16A-3; S16B-3; S17A-2,3; S17B |
| **HH39** | tip | sur epi | ncm | high | abs | abs | abs | low | abs | abs | abs | S14A-4; S14B; S15A; S15B-3; S16A; S16B; S17A-3; S17B |
| **HH39** | tip | sur epi | cyto | abs | abs | abs | low | abs | abs | abs | abs | S14A-4; S14B; S15A; S15B-3; S16A; S16B; S17A-3; S17B |
| **HH39** | tip | sur epi | nuc | abs | abs | abs | abs | low | abs | abs | abs | S14A-4; S14B; S15A; S15B-3; S16A; S16B; S17A-3; S17B |
| **HH39** | tip | bone | cyto | abs | low | abs | low | abs | high | abs | abs | S14A; S14B; S15A; S15B; S16A-2; S16B; S17A; S17B |
| **HH39** | tip | bone | nuc-cyto | med | abs | abs | abs | med | abs | abs | abs | S14A; S14B; S15A; S15B; S16A-2; S16B; S17A; S17B |
| **HH42** | base | cart | cyto | abs | low | abs | low | abs | med | abs | med | S18A-2; S18B; S19A; S19B; S20A; S20B-1; S21A; S21B-1,2 |
| **HH42** | base | cart | nuc-cyto | low | abs | abs | abs | abs | abs | abs | low | S18A-2; S18B; S19A; S19B; S20A; S20B-1; S21A; S21B-1,2 |
| **HH42** | base | mes | cyto | abs | low | abs | low | low | abs | abs | abs | S18A-2; S18B; S19A-1; S19B; S20A; S20B-1; S21A; S21B-1,2 |
| **HH42** | base | sur epi | cyto | abs | low | abs | low | abs | low | abs | low | S18A; S18B; S19A-1; S19B; S20A; S20B; S21A; S21B |
| **HH42** | base | sur epi | nuc-cyto | high | abs | abs | abs | low | abs | abs | low | S18A; S18B; S19A-1; S19B; S20A; S20B; S21A; S21B |
| **HH42** | base | ora epi | cyto | abs | low | abs | low | abs | abs | abs | low | S18A; S18B; S19A; S19B; S20A; S20B; S21A; S21B |
| **HH42** | base | ora epi | nuc-cyto | abs | abs | abs | abs | low | abs | abs | abs | S18A; S18B; S19A; S19B; S20A; S20B; S21A; S21B |
| **HH42** | base | ora epi | ncm | med | abs | abs | abs | low | abs | abs | abs | S18A; S18B; S19A; S19B; S20A; S20B; S21A; S21B |
| **HH42** | mid | conch | cyto | abs | med | abs | med | abs | high | med | low | S18A-3; S18B-2; S19A-2; S19B-1,2; S20A; S20B-3; S21A-1; S21B |
| **HH42** | mid | conch | ncm | high | abs | abs | abs | low | abs | abs | abs | S18A-3; S18B-2; S19A-2; S19B-1,2; S20A; S20B-3; S21A-1; S21B |
| **HH42** | mid | conch | nuc-cyto | abs | abs | abs | abs | low | abs | abs | low | S18A-3; S18B-2; S19A-2; S19B-1,2; S20A; S20B-3; S21A-1; S21B |
| **HH42** | mid | cart | cyto | abs | abs | abs | low | abs | high | abs | high | S18A-3; S18B-1,2,3; S19A-2; S19B-1,2; S20A-1; S20B-3; S21A-1; S21B-1 |
| **HH42** | mid | cart | nuc | abs | abs | abs | abs | high | abs | abs | abs | S18A-3; S18B-1,2,3; S19A-2; S19B-1,2; S20A-1; S20B-3; S21A-1; S21B-1 |
| **HH42** | mid | cart | nuc-cyto | low | abs | abs | abs | abs | abs | abs | low | S18A-3; S18B-1,2,3; S19A-2; S19B-1,2; S20A-1; S20B-3; S21A-1; S21B-1 |
| **HH42** | mid | sur epi | cyto | abs | low | abs | low | abs | low | abs | low | S18A-1; S18B; S19A; S19B; S20A-1; S20B; S21A; S21B |
| **HH42** | mid | sur epi | nuc-cyto | high | abs | abs | abs | low | abs | abs | low | S18A-1; S18B; S19A; S19B; S20A-1; S20B; S21A; S21B |
| **HH42** | mid | mes | cyto | abs | low | abs | low | low | abs | abs | low | S18A-1,3; S18B-3; S19A-2; S19B-2; S20A-1; S20B-2,3; S21A-1; S21B-1 |
| **HH42** | mid | mes | nuc-cyto | low | abs | abs | abs | low | abs | abs | low | S18A-1,3; S18B-3; S19A-2; S19B-2; S20A-1; S20B-2,3; S21A-1; S21B-1 |
| **HH42** | mid | bone | cyto | abs | low | abs | low | med | abs | abs | abs | S18A-1; S18B-3; S19A-2; S19B; S20A-1; S20B-2; S21A; S21B |
| **HH42** | mid | bone | nuc-cyto | med | abs | abs | abs | med | abs | med | abs | S18A-1; S18B-3; S19A-2; S19B; S20A-1; S20B-2; S21A; S21B |
| **HH42** | mid | ora epi | cyto | abs | low | abs | low | abs | abs | abs | low | S18A; S18B; S19A; S19B; S20A; S20B; S21A; S21B |
| **HH42** | mid | ora epi | ncm | med | abs | abs | abs | low | abs | abs | abs | S18A; S18B; S19A; S19B; S20A; S20B; S21A; S21B |
| **HH42** | mid | nas epi | cyto | abs | med | low | med | abs | med | abs | low | S18A; S18B; S19A; S19B-1; S20A; S20B; S21A; S21B |
| **HH42** | mid | nas epi | nuc-cyto | med | abs | abs | abs | low | abs | abs | low | S18A; S18B; S19A; S19B-1; S20A; S20B; S21A; S21B |
| **HH42** | tip | cart | cyto | abs | low | abs | low | abs | med | abs | low | S18A; S18B; S19A; S19B; S20A; S20B; S21A; S21B-3 |
| **HH42** | tip | sur epi | cyto | abs | med | abs | med | abs | low | med | low | S18A-4; S18B; S19A; S19B; S20A-2; S20B; S21A-2; S21B-3 |
| **HH42** | tip | sur epi | nuc-cyto | abs | abs | abs | abs | med | abs | abs | abs | S18A-4; S18B; S19A; S19B; S20A-2; S20B; S21A-2; S21B-3 |
| **HH42** | tip | sur epi | ncm | med | abs | abs | abs | low | abs | abs | abs | S18A-4; S18B; S19A; S19B; S20A-2; S20B; S21A-2; S21B-3 |
| **HH42** | tip | mes | cyto | abs | low | abs | low | low | abs | abs | abs | S18A-4; S18B; S19A; S19B; S20A-2; S20B; S21A-2; S21B-3 |
| **HH42** | tip | mes | nuc-cyto | low | abs | abs | abs | low | abs | abs | low | S18A-4; S18B; S19A; S19B; S20A-2; S20B; S21A-2; S21B-3 |
| **HH42** | tip | bone | cyto | abs | low | abs | low | abs | low | abs | abs | S18A; S18B; S19A; S19B; S20A; S20B; S21A; S21B |
| **HH42** | tip | bone | nuc-cyto | med | abs | abs | abs | med | abs | med | abs | S18A; S18B; S19A; S19B; S20A; S20B; S21A; S21B |

*FIGURE REFERENCES ARE LISTED IN FACTOR ORDER: B-CATENIN; BMP4; CAM; DKK3; FGF8; IHH; TGFB2; WNT4; EXPRESSION: ABS=ABSENT; LOW=LOW, MED=MEDIUM, HIGH=HIGH*
*CYTO = CYTOPLASMIC; NUC = NUCLEAR; NUC-CYTO = NUCLEAR-CYTOPLASMIC; NCM = NUCLEAR-CYTOPLASMIC-MEMBRANE*
*CART = CARTILAGE; CONCH = CONCHAE; SUR EPI = SURFACE (EXTERIOR) EPITHELIUM; ORA EPI = ORAL (INTERIOR) EPITHELIUM; MES = MESENCHYME; NAS EPI = NASAL EPITHELIUM; \*TGFβ2 NUCLEAR LOCATION REFERS TO PERINUCLEAR*

**TABLE S2.**
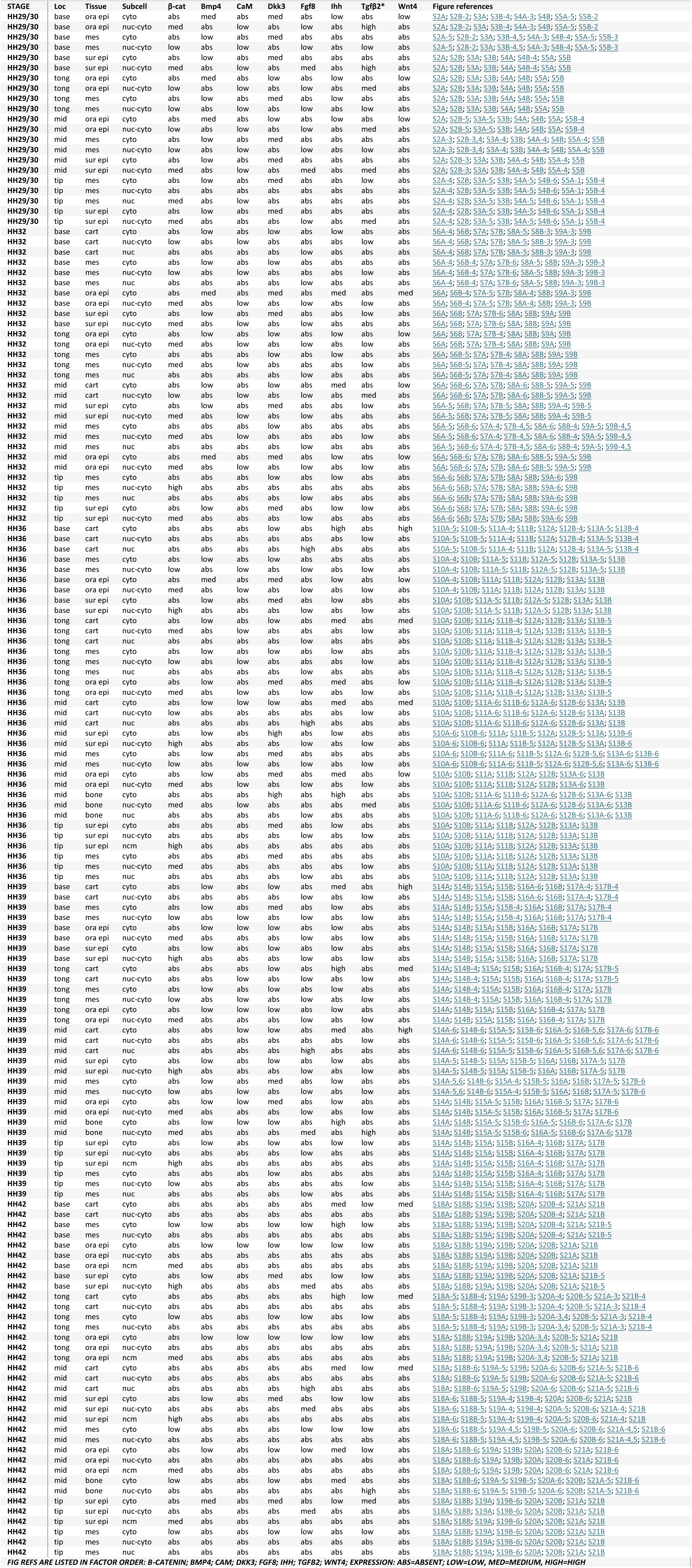
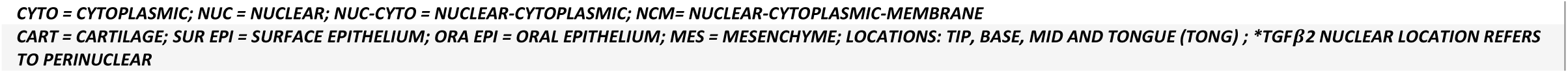
SUMMARY OF PROTEIN EXPRESSION ACROSS DEVELOPMENTAL STAGES, AXIAL LOCATIONS, AND SUBCELLULAR CONTEXTS FOR LOWER BEAK.

**TABLE S4.** Protein-pair tissue switches associated with subcellular relocation.

| **Protein pair** | **Beak / stage** | **Tissue shift (Pearson *r*’s)** | **Subcellular relocation / compartment switch** |
|---|---|---|---|
| β-catenin–Fgf8 | Upper HH32 | surface epithelium (1.00), oral epithelium (0.90), cartilage (0.46) → mesenchyme (-0.38) | Fgf8: nuclear-cytoplasmic → nuclear |
| CaM–Fgf8 | Lower HH39 | mesenchyme (0.79) → oral epithelium (-1.00) | CaM: nuclear-cytoplasmic → cytoplasmic;<br>Fgf8 → nuclear in cartilage |
| β-catenin–CaM | Upper HH29/30 | surface epithelium (0.58) → oral epithelium (-0.90) | CaM: nuclear-cytoplasmic → cytoplasmic |
| Bmp4–Tgfβ2 | Lower HH32 | surface epithelium (1.00) → oral epithelium (-0.93), cartilage (-0.61) | Tgfβ2: cytoplasmic →perinuclear-cytoplasmic |
| CaM–Tgfβ2 | Lower HH32 | oral epithelium (1.00), mesenchyme (1.00), cartilage (0.92) → surface epithelium (-0.71) | Tgfβ2: perinuclear-cytoplasmic → cytoplasmic |
| Dkk3–Tgfβ2 | Lower HH32 | surface epithelium (1.00) → oral epithelium (-0.93), cartilage (-0.61) | Tgfβ2: cytoplasmic → perinuclear-cytoplasmic |
| Fgf8–Tgfβ2 | Lower HH32 | oral epithelium (1.00) → surface epithelium (-1.00) | Tgfβ2: perinuclear-cytoplasmic → cytoplasmic |
| β-catenin–Tgfβ2 | Lower HH32 | oral epithelium (1.00), mesenchyme (0.56), cartilage (0.92) → surface epithelium (-1.00) | Tgfβ2: perinuclear-cytoplasmic → cytoplasmic |
| Bmp4–CaM | Lower HH39 | oral epithelium (1.00) → mesenchyme (-0.50) | CaM: cytoplasmic → nuclear-cytoplasmic |
| CaM–Dkk3 | Lower HH39 | oral epithelium (0.89) → mesenchyme (-0.58) | CaM: cytoplasmic → nuclear-cytoplasmic |
| CaM–Ihh | Lower HH39 | oral epithelium (1.00) → mesenchyme (-0.50) | CaM: cytoplasmic → nuclear-cytoplasmic |
| CaM–Tgfβ2 | Lower HH39 | mesenchyme (0.79) → oral epithelium (-1.00) | CaM: nuclear-cytoplasmic → cytoplasmic |
| β-catenin–CaM | Lower HH39 | mesenchyme (0.79) → oral epithelium (-1.00) | CaM: nuclear-cytoplasmic → cytoplasmic |
| Bmp4–Fgf8 | Lower HH42 | oral epithelium (1.00) → surface epithelium (-0.55) | Fgf8: cytoplasmic → nuclear-cytoplasmic |
| Dkk3–Fgf8 | Lower HH42 | oral epithelium (1.00), mesenchyme (0.79) → surface epithelium (-0.60) | Fgf8: cytoplasmic → nuclear-cytoplasmic |
| Fgf8–Ihh | Lower HH42 | oral epithelium (0.92), mesenchyme (0.52) → surface epithelium (-0.60) | Fgf8: cytoplasmic → nuclear-cytoplasmic |
| Fgf8–Tgfβ2 | Lower HH42 | oral epithelium (1.00), mesenchyme (1.00) → surface epithelium (-0.55) | Fgf8: cytoplasmic → nuclear-cytoplasmic |
| β-catenin–Dkk3 | Lower HH42 | mesenchyme (0.79) → surface epithelium (-0.59), oral epithelium (-0.50) | β-catenin: cytoplasmic → nuclear-cytoplasmic-membranal |
| β-catenin–Fgf8 | Lower HH42 | mesenchyme (1.00) → oral epithelium (-0.50) | β-catenin: cytoplasmic → nuclear-cytoplasmic-membranal |
| β-catenin–Tgfβ2 | Lower HH42 | mesenchyme (1.00) → surface epithelium (-0.53), oral epithelium (-0.50) | β-catenin: cytoplasmic → nuclear-cytoplasmic-membranal |
| CaM–Dkk3 | Upper HH29/30 | oral epithelium (0.90) → surface epithelium (-0.58), mesenchyme (-0.61) | CaM: cytoplasmic → nuclear-cytoplasmic |
| CaM–Tgfβ2 | Upper HH29/30 | surface epithelium (0.87), mesenchyme (0.71) → oral epithelium (-1.00) | CaM: nuclear-cytoplasmic → cytoplasmic |
| Bmp4–Tgfβ2 | Upper HH32 | surface epithelium (0.58) → oral epithelium (-1.00) | Tgfβ2: cytoplasmic → perinuclear-cytoplasmic |
| CaM–Tgfβ2 | Upper HH32 | oral epithelium (0.60), mesenchyme (0.76) → surface epithelium (-0.57), cartilage (-0.50) | CaM: nuclear-cytoplasmic → nuclear;<br>Tgfβ2: perinuclear-cytoplasmic → cytoplasmic |
| Dkk3–Tgfβ2 | Upper HH32 | surface epithelium (1.00) → oral epithelium (-0.90) | Tgfβ2: cytoplasmic → perinuclear-cytoplasmic |
| Fgf8–Tgfβ2 | Upper HH32 | oral epithelium (1.00) → surface epithelium (-1.00) | Tgfβ2: perinuclear-cytoplasmic → cytoplasmic |
| β-catenin–Tgfβ2 | Upper HH32 | oral epithelium (0.90), cartilage (1.00) → surface epithelium (-1.00) | Tgfβ2: perinuclear-cytoplasmic → cytoplasmic |
| CaM–Ihh | Upper HH36 | cartilage (0.53) → surface epithelium (-0.55), oral epithelium (-1.00) | CaM: cytoplasmic/nuclear → nuclear-cytoplasmic |
| Dkk3–Tgfβ2 | Upper HH42 | surface epithelium (1.00), oral epithelium (1.00) → mesenchyme (-0.61), bone (-1.00) | Tgfβ2: cytoplasmic → perinuclear-cytoplasmic |
| Ihh–Tgfβ2 | Upper HH42 | surface epithelium (0.91), oral epithelium (1.00) → mesenchyme (-0.61), bone (-1.00) | Tgfβ2: cytoplasmic → perinuclear-cytoplasmic |

**Figure S1.**
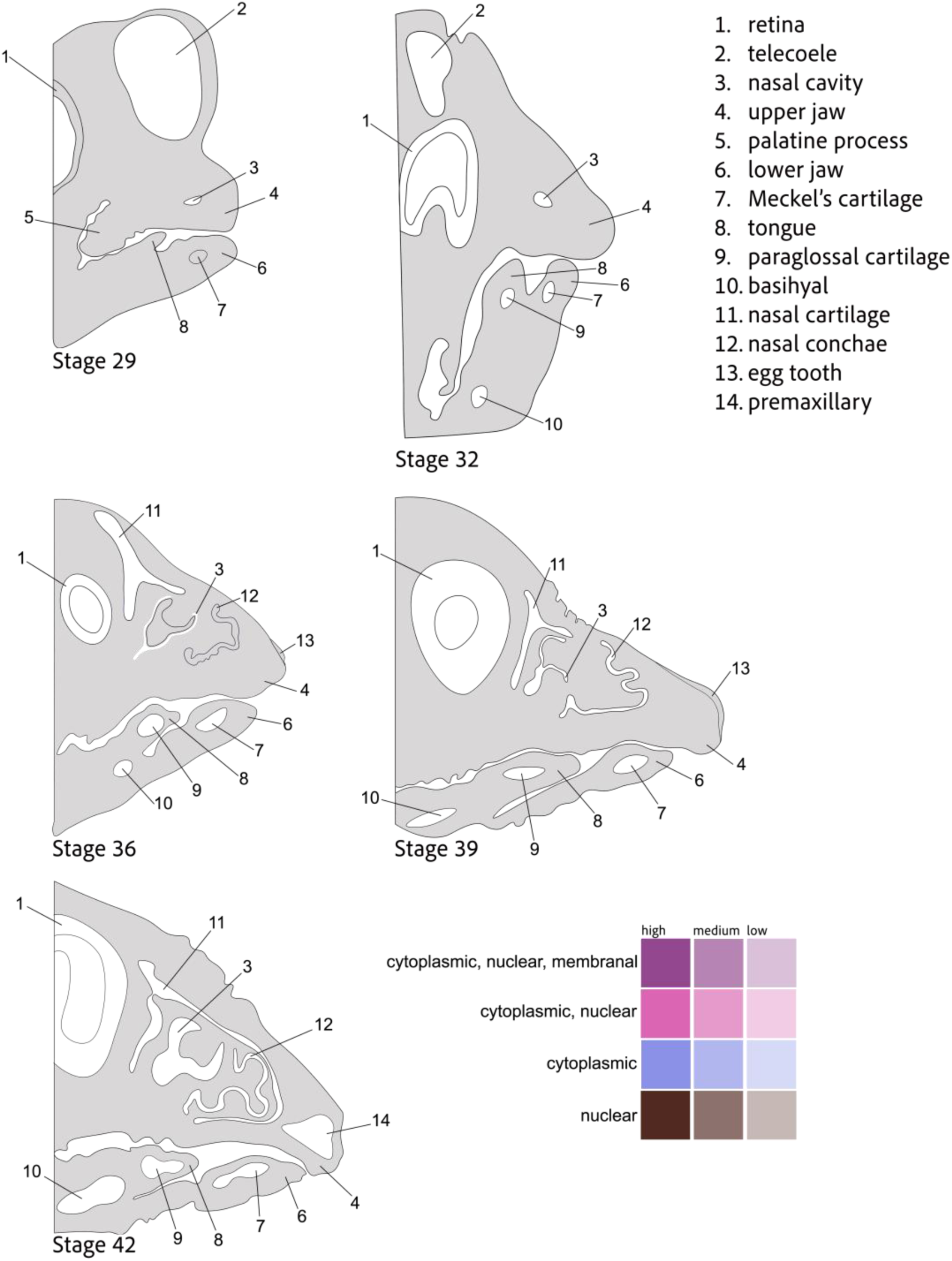
**Illustrated map of beak morphology** of mid-sagittal sections across 5 developmental stages including labels for primary anatomical structures. Color key for cellular location used to annotate protein expression location across Figures S2-S21. Nuclear expression includes perinuclear.

**Figure S2.**
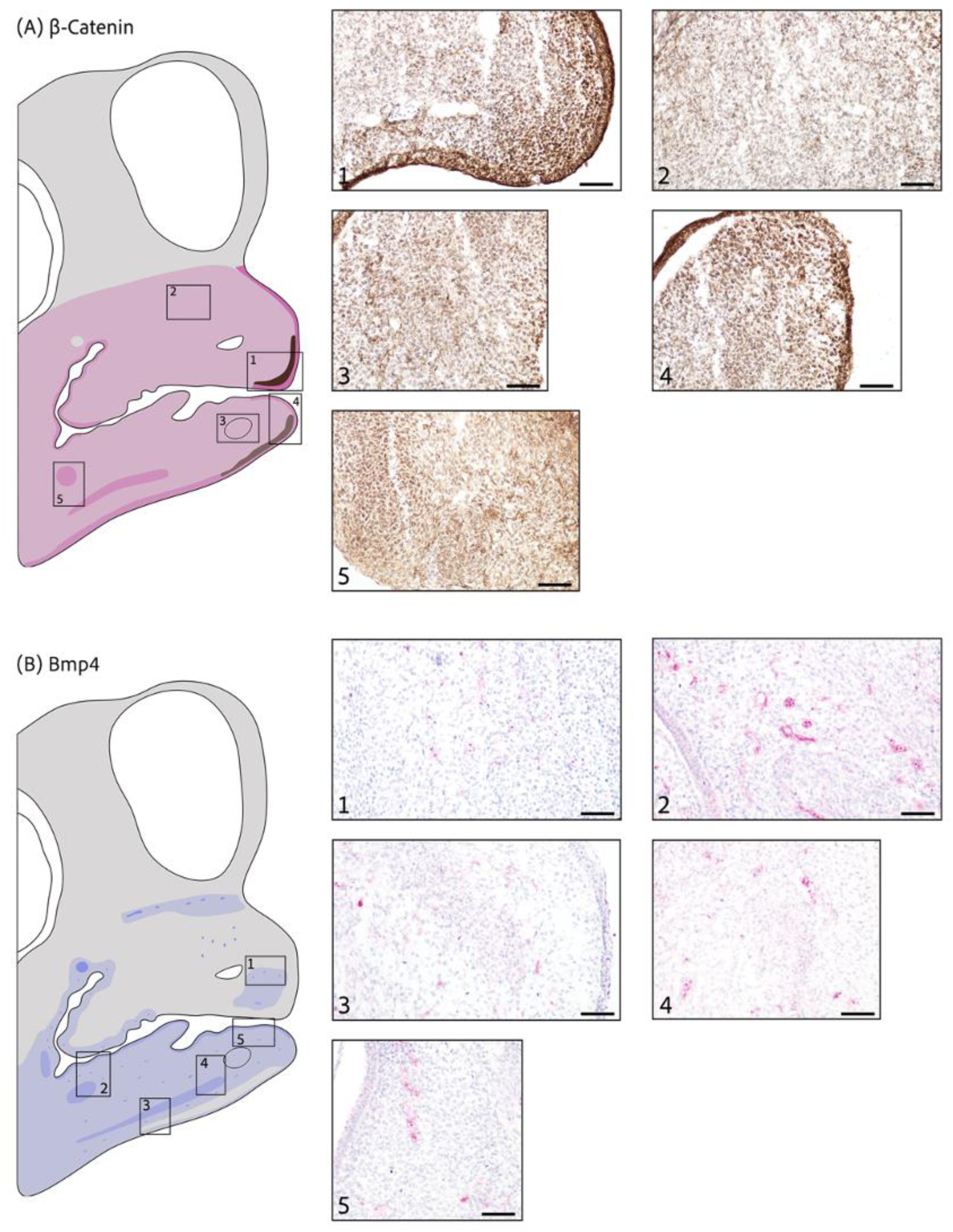
β-catenin and Bmp4 expression at stage HH29/30. (A) **β-catenin** (brown stain) shows overall weak cytoplasmic and nuclear expression across the upper and lower jaws with (1) high nuclear expression at the tip in mesenchymal cells and both high cytoplasmic and nuclear epithelial expression. No expression in blood cells. (2) Low cytoplasmic and nuclear expression in the upper jaw. (3) Low cytoplasmic and nuclear expression in the lower jaw, with Meckel’s cartilage condensation not formed yet. (4) Medium nuclear expression along the outer edge of the tip in the mesenchyme cells and medium cytoplasmic and nuclear epithelial expression. (5) Medium cytoplasmic and nuclear expression in condensations that will become future myogenic areas. (B) **Bmp4** (red stain) is rarely present in the upper beak except for some areas (1) that show very weak cytoplasmic expression. Low cytoplasmic expression across the lower beak with (2, 3, 4) medium cytoplasmic expression in cell condensations that will form future myogenic areas. (5) Weak cytoplasmic expression in the mesenchyme while the oral epithelium shows medium cytoplasmic expression. Medium and high cytoplasmic expression in blood cells. Scale bars are 50*μ*m. (Back to Table S1)

**Figure S3.**
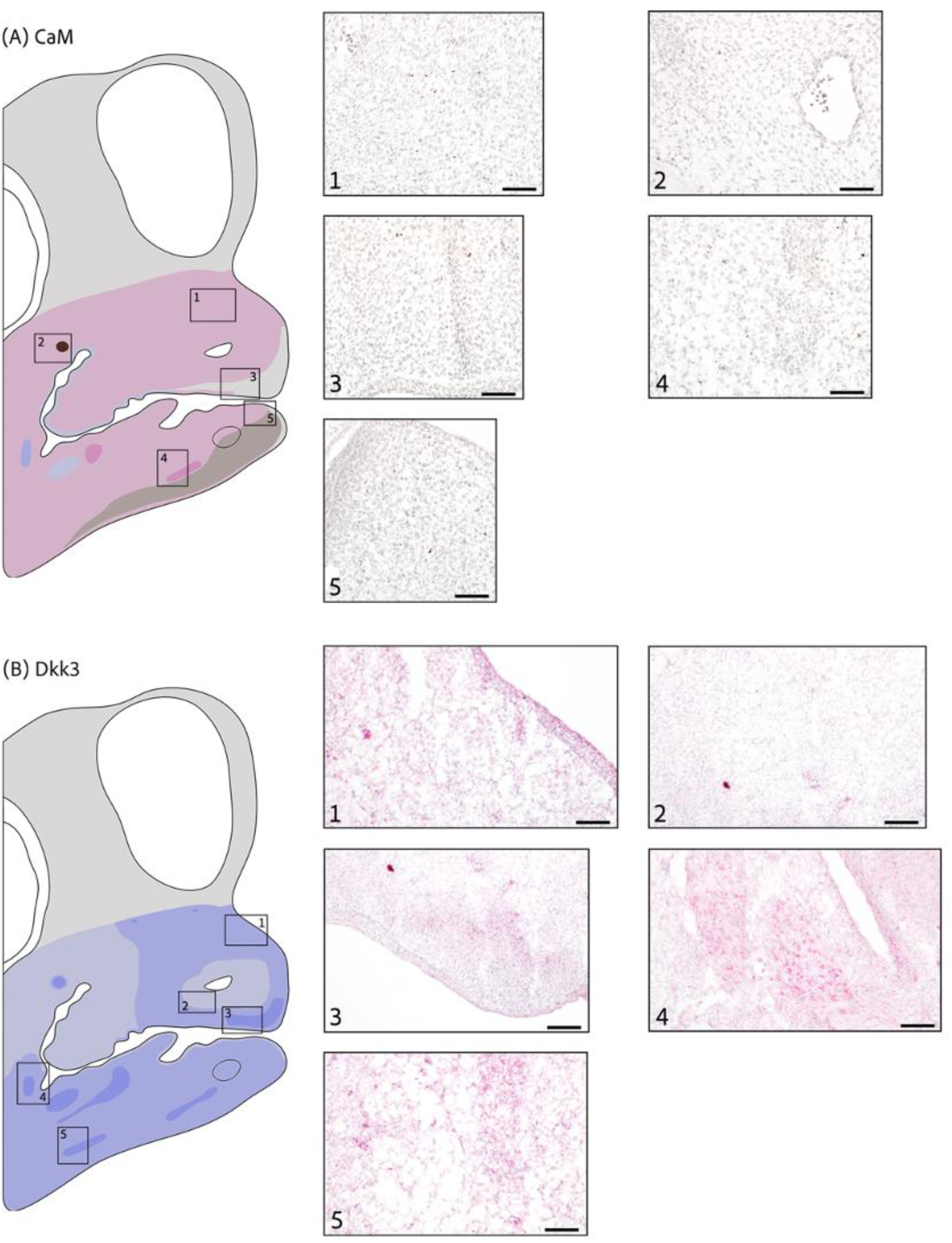
CaM and Dkk3 expression at stage HH29/30. (A) **CaM** (brown stain) shows weak cytoplasmic and nuclear expression across both the upper (1) and lower jaw. (2) Blood cells show high nuclear expression. (3) No expression at the tip of the upper jaw. (4) Medium cytoplasmic and nuclear expression in myogenic condensation. (5) Weak cytoplasmic and nuclear expression in contact with low nuclear expression that is present at the mesenchyme of the outer edge in the lower jaw. (B) **Dkk3** (red stain) shows medium cytoplasmic expression in the upper jaw (1) and especially in the lower jaw. (2) Weak cytoplasmic expression in some areas of the upper jaw with (3) high cytoplasmic expression at the tip. (4,5) High cytoplasmic expression in potential myogenic areas. Scale bars are 50*μ*m. (Back to Table S1)

**Figure S4.**
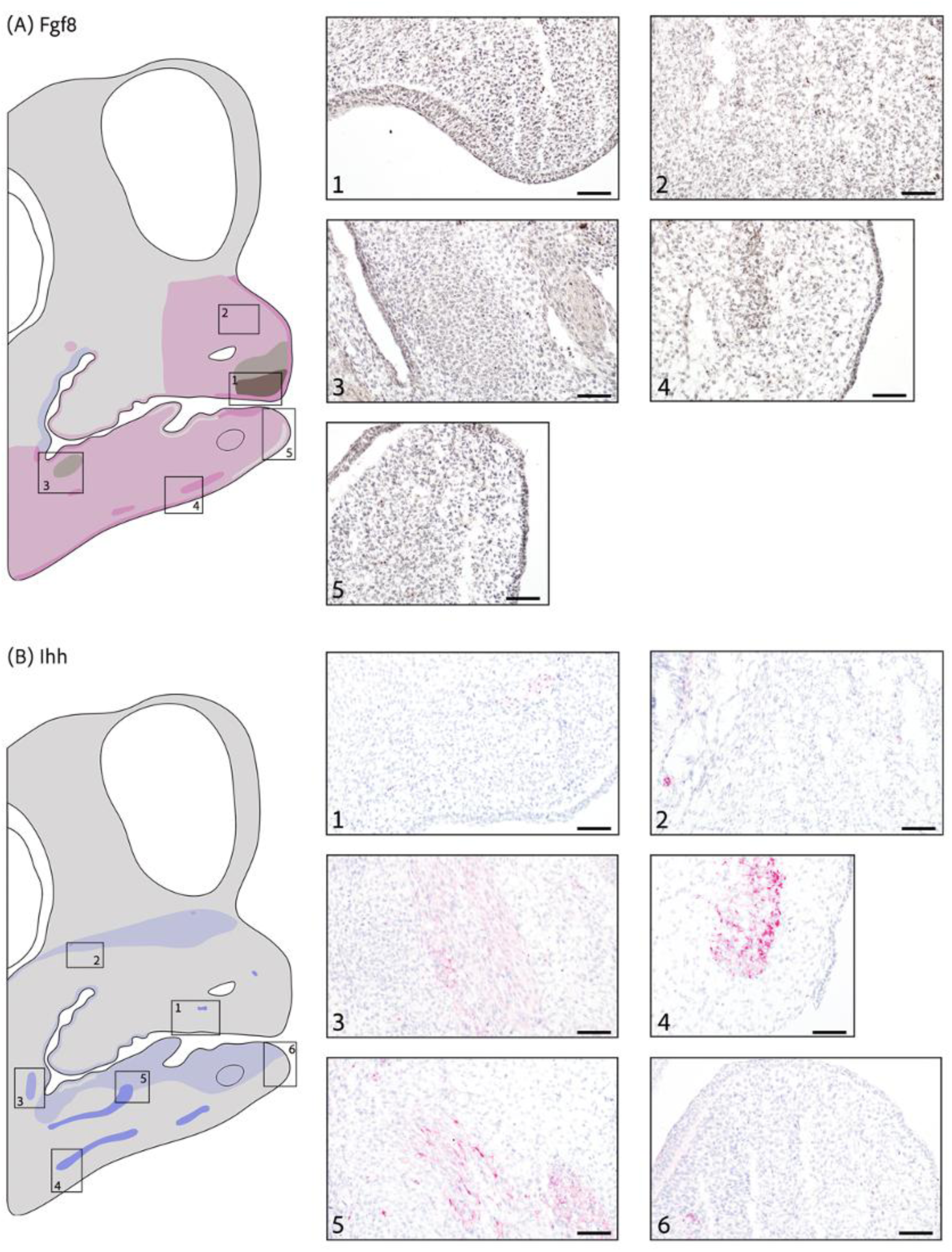
Fgf8 and Ihh expression at stage HH29/30. (A) **Fgf8** (brown stain) is weakly expressed in (2) both the cytoplasm and nucleus at half of the upper jaw and across the whole lower jaw with (1) medium and low nuclear expression in the mesenchyme and medium cytoplasmic and nuclear expression in the epithelium at the tip of the upper jaw and (5) no expression at the outer edge of the lower jaw. (3) Low nuclear expression in cell condensation and (4) medium cytoplasmic and nuclear expression in potential myogenic area. (B) **Ihh** (red stain) expression is localized with weak cytoplasmic expression in both (2) the upper jaw and (6) the lower jaw. (1) Blood cells show medium cytoplasmic expression. Most expression is localized in potential myogenic areas that show (3) medium and (4, 5) high cytoplasmic expression. Scale bars are 50*μ*m. (Back to Table S1)

**Figure S5.**
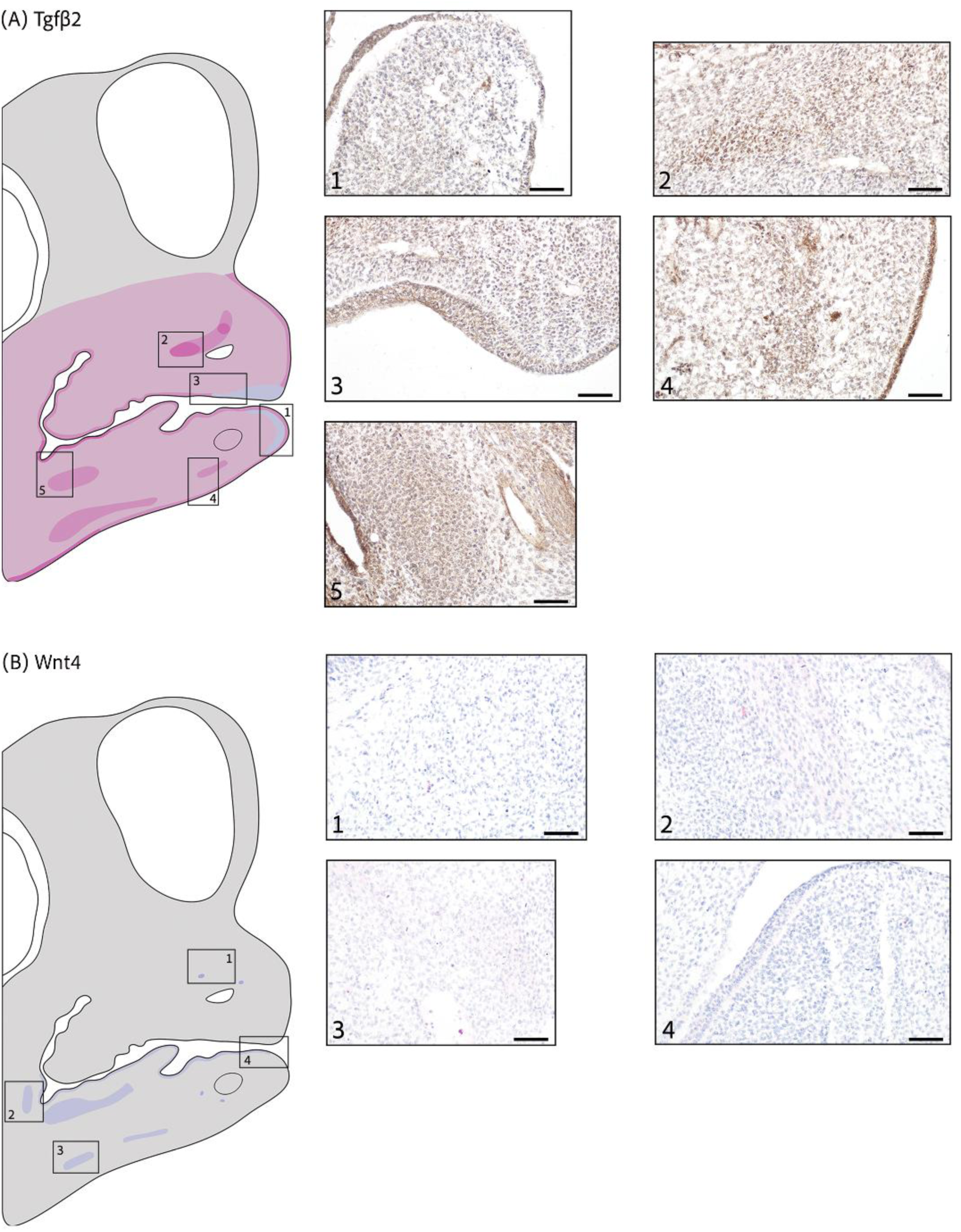
Tgfβ2 and Wnt4 expression at stage HH29/30. (A) **Tgfβ2** shows low cytoplasmic and perinuclear expression across the upper and lower jaw with weak cytoplasmic expression at the tip of the (3) upper and (1) lower jaw. (2) High cytoplasmic and perinuclear expression in the upper jaw and medium expression in (4) potential muscle tissue and (5) cell condensation in the lower jaw. (B) **Wnt4** (red stain) is rarely present in the upper jaw and highly localized in the lower jaw. (1) blood cells show weak cytoplasmic expression. Low cytoplasmic expression in (2, 3) future myogenic areas and (4) oral epithelium in the lower jaw. Scale bars are 50*μ*m. (Back to Table S1)

**Figure S6.**
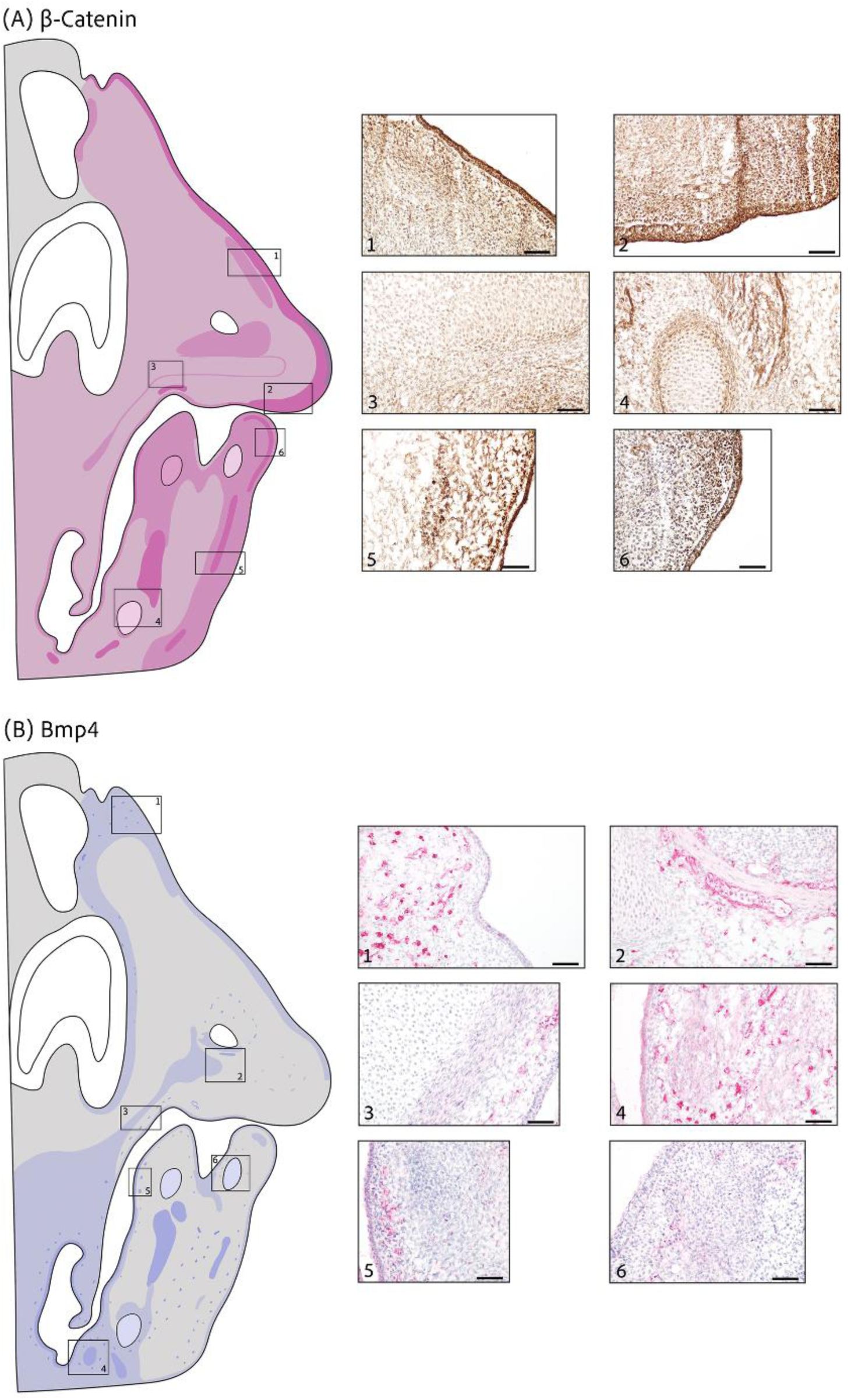
β-catenin and Bmp4 expression at stage HH32. (A) **β-catenin** (brown stain) is weakly present in the cytoplasm and nucleus across both the upper and lower jaw with (1, 2) high expression across the outer edge of the upper jaw and (1) medium expression in potential muscle condensation. (3) Low cytoplasmic and nuclear expression in a large condensation of the maxillary prominence with medium expression at the borderline perichondrogenic area in the upper jaw. Medium expression across the outer edge of the lower jaw. (4) Low cytoplasmic and nuclear expression in the basihyal cartilage of the mandibular prominence with medium expression in the perichondrogenic area. High cytoplasmic and nuclear expression in (4, 5) myogenic areas and (6) in the mesenchyme at the tip of the lower jaw. (B) **Bmp4** (red stain) shows weak localized cytoplasmic expression that is predominantly present in the epithelium. Hematopoietic cells show (1) high and (2) medium cytoplasmic expression. (3) Low cytoplasmic expression in the developing myogenic area next to the cartilage. Lower jaw (4) muscle tissue and (5) inner epithelium shows medium cytoplasmic expression. (6) Weak cytoplasmic expression showed in the early formed condensation that will give rise to Meckel’s cartilage. Scale bars are 50*μ*m. (Back to Table S1)

**Figure S7.**
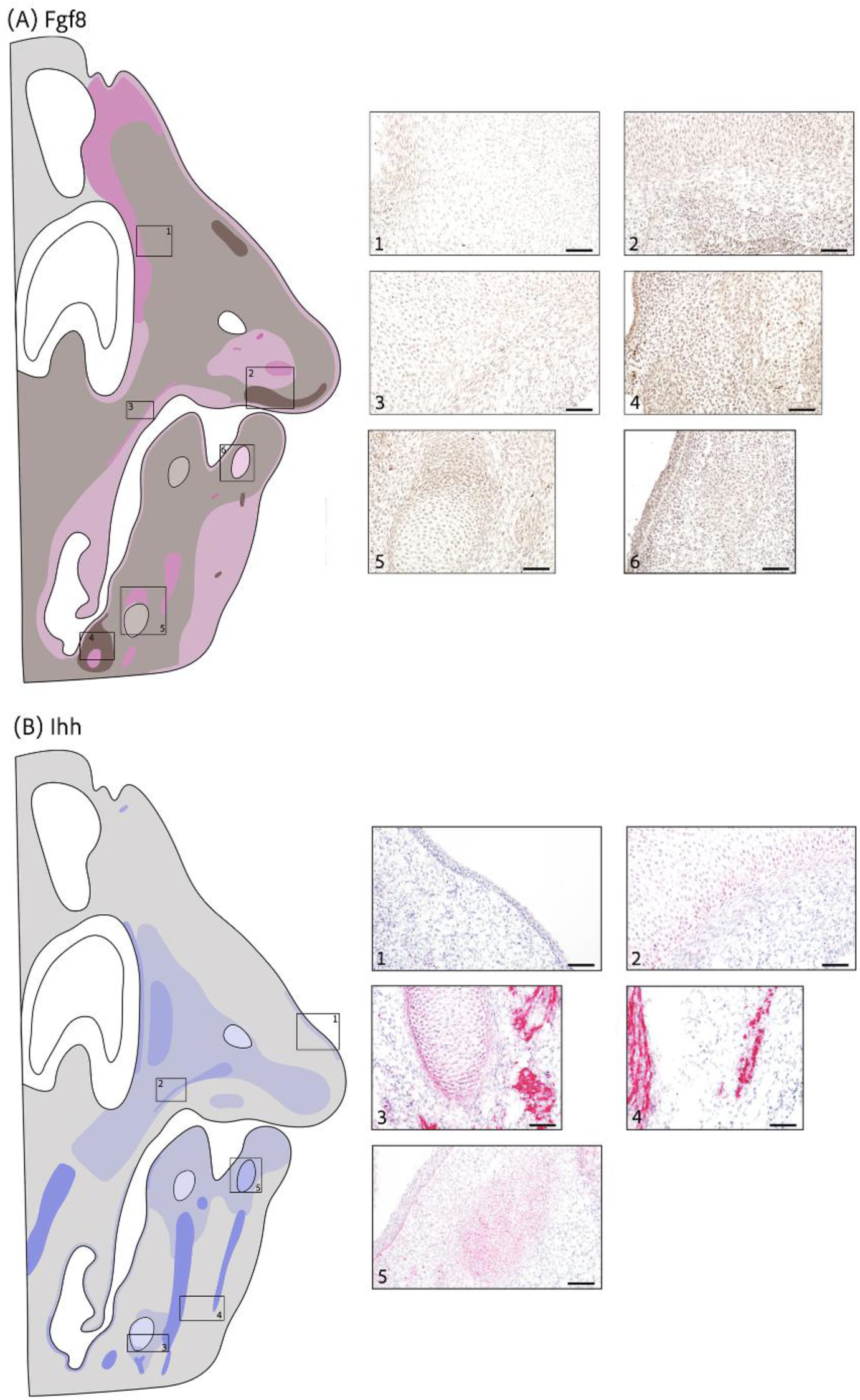
Fgf8 and Ihh expression at stage HH32. (A) **Fgf8** (brown stain) mainly shows weak nuclear expression in the mesenchyme while both cytoplasmic and nuclear expression in the epithelium. Medium cytoplasmic and nuclear expression in (1) mesenchyme next to the retina, (2) terminus of the large condensation at the maxillary prominence and (3) developing muscle next to cartilage area. (2) Medium nuclear expression at the tip of the upper jaw. (4) Medium nuclear expression surrounding cell condensation with additional cytoplasmic expression. (5) Basihyal cartilage with low nuclear expression, perichondrogenic area with low cytoplasmic and nuclear expression. Developing muscle above with medium cytoplasmic and nuclear expression. (6) Cell condensation preceding Meckel’s cartilage with low cytoplasmic and nuclear expression. (B) **Ihh** (red stain) shows scarce epithelium expression with (1) some areas of low cytoplasmic expression. (2, 3) Cartilage contains weak cytoplasmic expression, but perichondrogenic area shows stronger expression. (3, 4) Very strong expression in muscle tissue. (5) Medium cytoplasmic expression in cell condensation that will form Meckel’s cartilage. Scale bars are 50*μ*m. (Back to Table S1)

**Figure S8.**
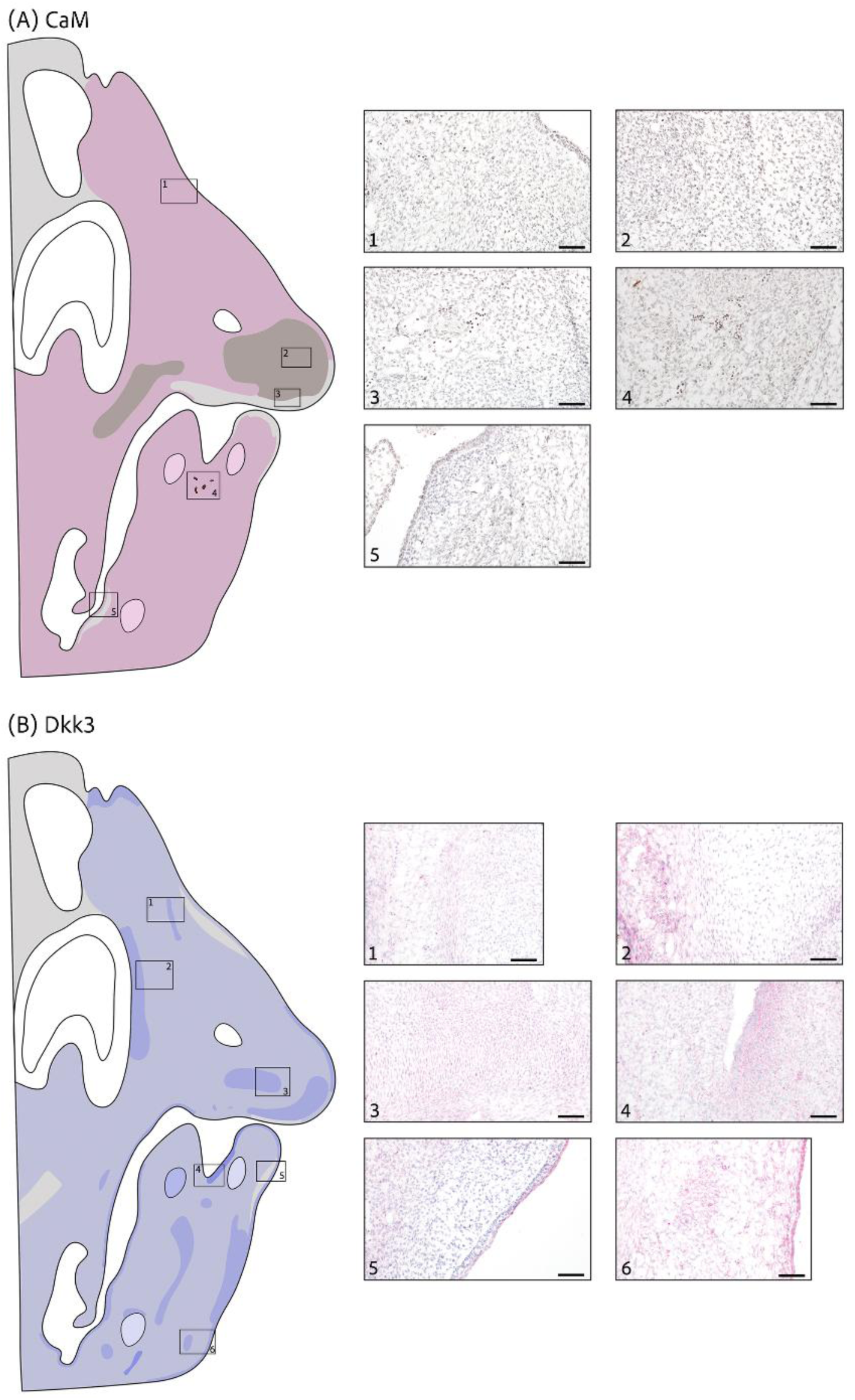
CaM and Dkk3 expression at stage HH32. (A) **CaM** (brown stain) overall (1) weak cytoplasmic and nuclear expression with some areas of (2) low nuclear expression in the upper jaw and (3, 5) no expression. (4) High nuclear expression in blood cells. (B) **Dkk3** (red stain) shows widely distributed low cytoplasmic expression with areas of medium cytoplasmic expression in both upper and lower jaw. In the upper jaw, medium cytoplasmic expression (1) in potential muscle next to developing cartilage (2) next to the retina and (3) in a large condensation of the maxillary prominence. In the lower jaw, (4) contrast between high and low cytoplasmic expression in mesenchyme and epithelium and (6) medium cytoplasmic expression in potential myogenic areas. (5) Scattered areas with no expression. Scale bars are 50*μ*m. (Back to Table S1)

**Figure S9.**
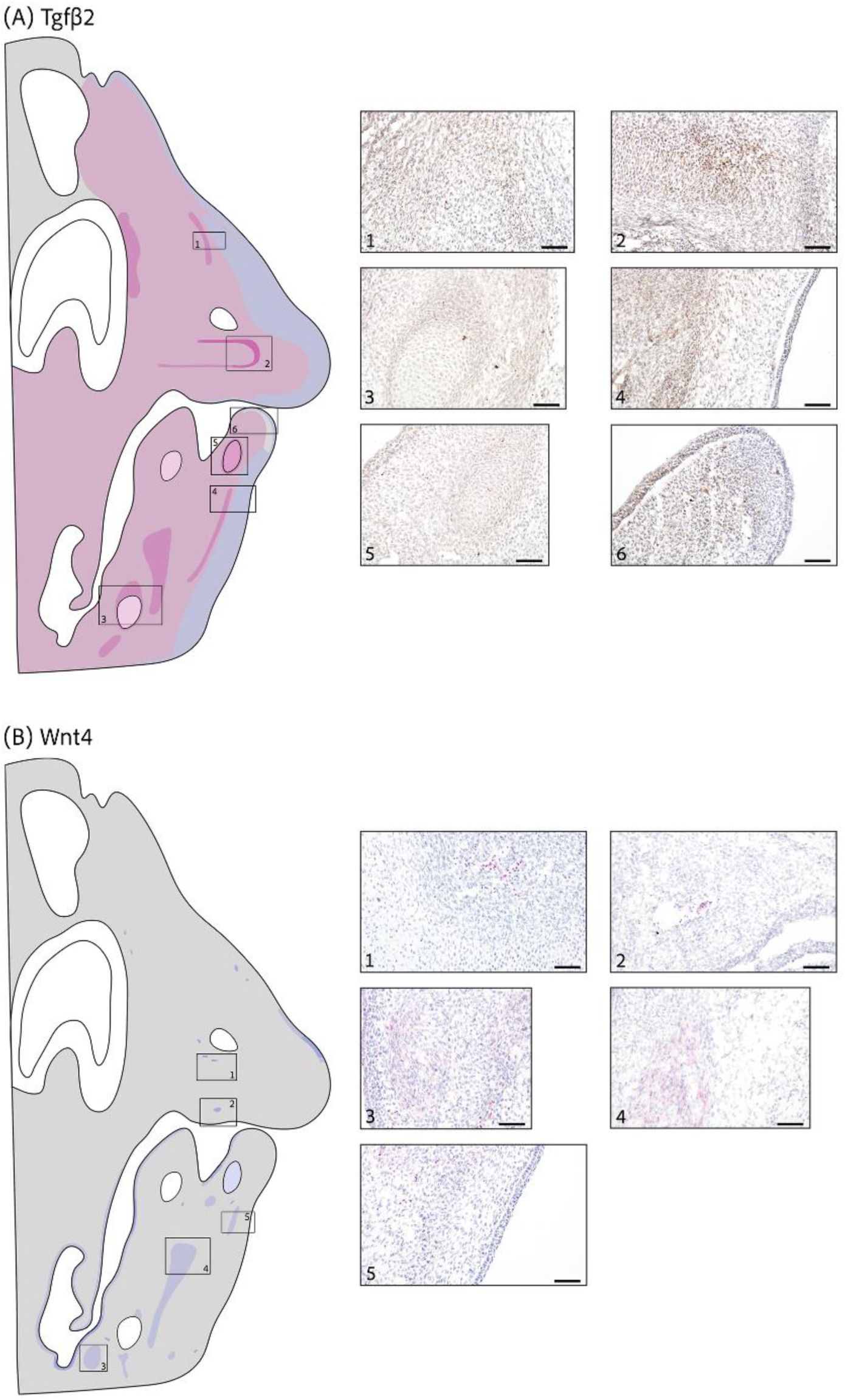
Tgfβ2 and Wnt4 expression at stage HH32. (A) **Tgfβ2** (brown stain) is widely and weakly expressed in the cytoplasm and perinuclearly. Only cytoplasmic expression is maintained in the outer edge of the (1) upper and (4) lower jaw. (2) High cytoplasmic and perinuclear expression surrounding the large condensation at the maxillary prominence. (3) Basihyal cartilage shows low cytoplasmic and nuclear expression while surrounding muscle and perinuclear area show medium expression. (5) Medium cytoplasmic and nuclear expression in cell condensation that will develop into Meckel’s cartilage. (6) No expression at the tip of the lower jaw. (B) **Wnt4** (red stain) is locally expressed mainly in the lower jaw. (1, 2) Blood cells show medium cytoplasmic expression. (3, 4) Muscle tissue and (5) potential myogenic area shows low cytoplasmic expression. Scale bars are 50*μ*m. (Back to Table S1)

**Figure S10.**
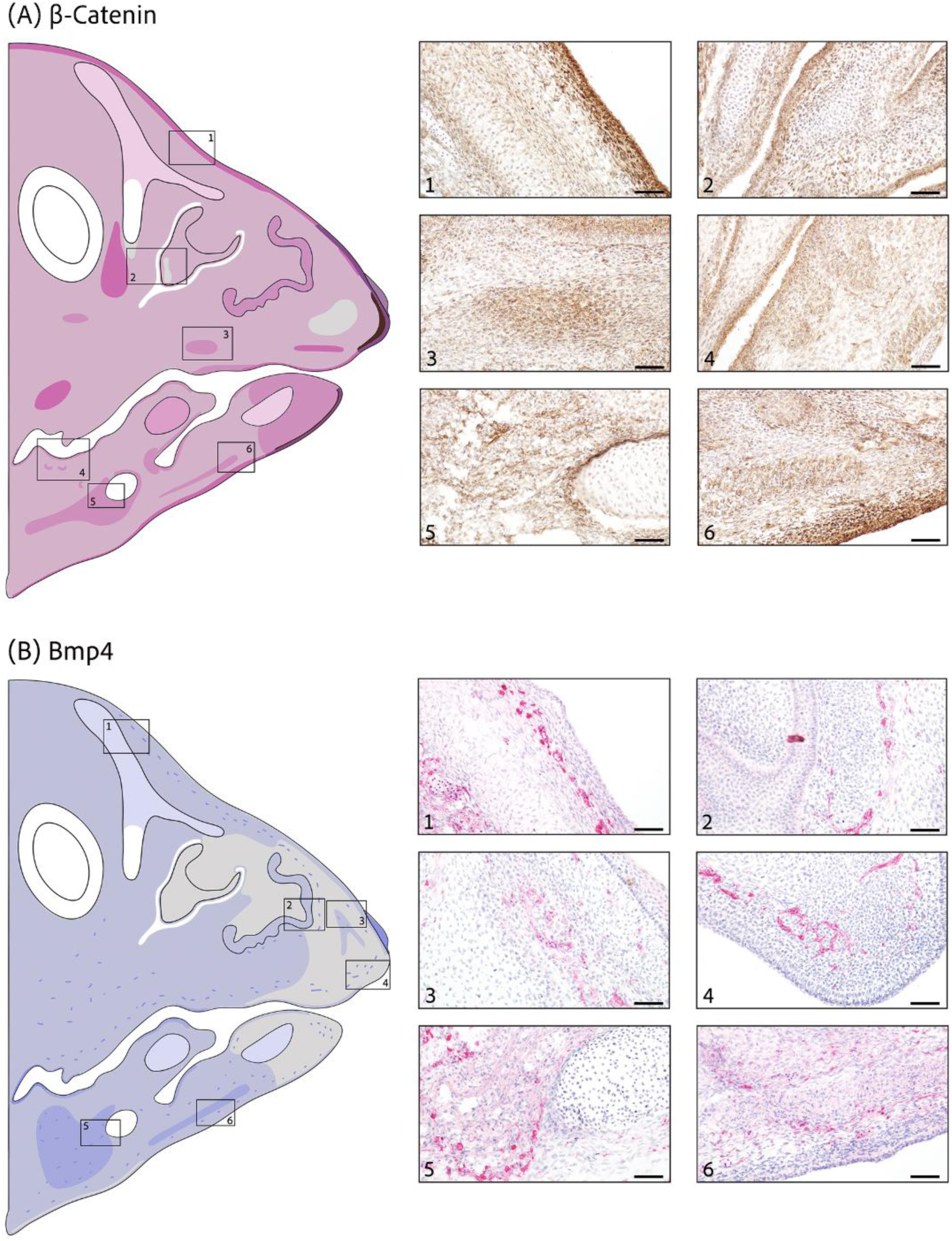
β-catenin and Bmp4 expression at stage HH36. (A) **β-catenin** (brown stain) overall weak cytoplasmic and nuclear expression. (1, 6) High cytoplasmic and nuclear expression in epithelium and nearby mesenchyme across the outer edge of the upper and lower jaw. (2) No expression in some areas of the developing nasal cartilage. Medium cytoplasmic and nuclear expression in (3) cell condensation and (4) epithelial buds. Absence of expression in (5) basihyal cartilage and medium cytoplasmic and nuclear expression in (5, 6) myogenic areas. (B) **Bmp4** (red stain) shows weak cytoplasmic expression across the beak but is absent at the tip. (1) Weak cytoplasmic expression in developing nasal cartilage and (2) nasal conchae, but no expression in surrounding mesenchyme. (3) Premaxillary condensation with low cytoplasmic expression. (4) High cytoplasmic expression in blood cells. (5) No expression in basihyal cartilage and medium expression in (5, 6) muscle tissue. Scale bars are 50*μ*m. (Back to Table S1)

**Figure S11.**
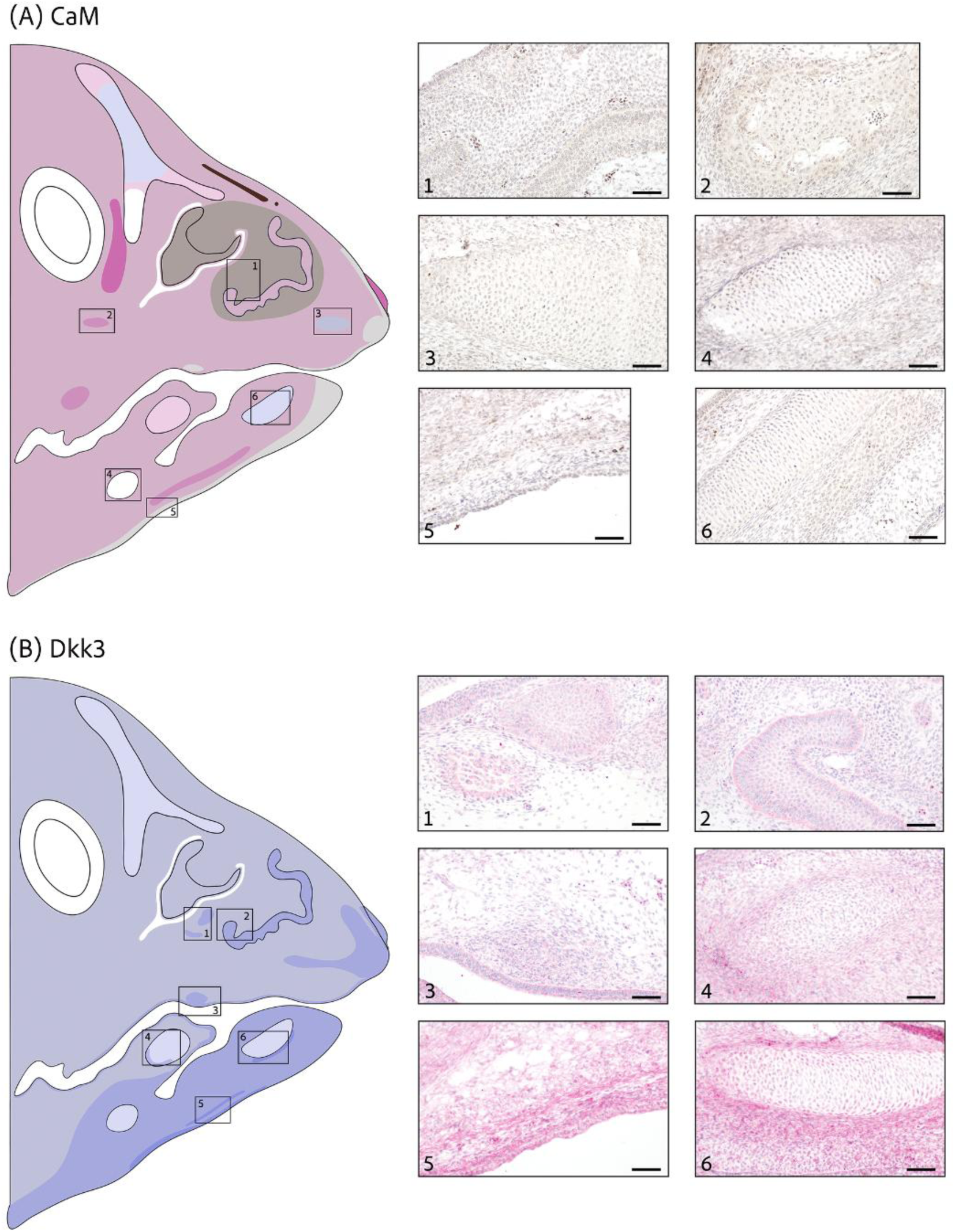
CaM and Dkk3 expression at stage HH36. (A) **CaM** (brown stain) shows overall weak cytoplasmic and nuclear expression, (1) including the nasal conchae but with low nuclear expression in the surrounding mesenchyme and developing nasal cartilage. (2) Medium cytoplasmic and nuclear expression in emerging osteogenic region. Weak cytoplasmic expression in developing (3) premaxillary and (6) Meckel’s cartilage. No expression in (4) basihyal cartilage or tip of upper and (5) outer edge of lower jaw. Strong nuclear expression in blood cells. (B) **Dkk3** (red stain) shows overall weak cytoplasmic expression in upper jaw and medium in lower jaw. (1) Emerging osteogenic region, (2) nasal conchae and (3) cell condensation in upper jaw shows medium cytoplasmic expression. (4) Paraglossal condensation with low expression and moderate expression in surrounding mesenchyme. (5) Contrast of medium and high cytoplasmic expression in both mesenchyme and epithelium. (6) Meckel’s cartilage with weak cytoplasmic expression and moderate to high expression in surrounding mesenchyme. Scale bars are 50*μ*m. (Back to Table S1)

**Figure S12.**
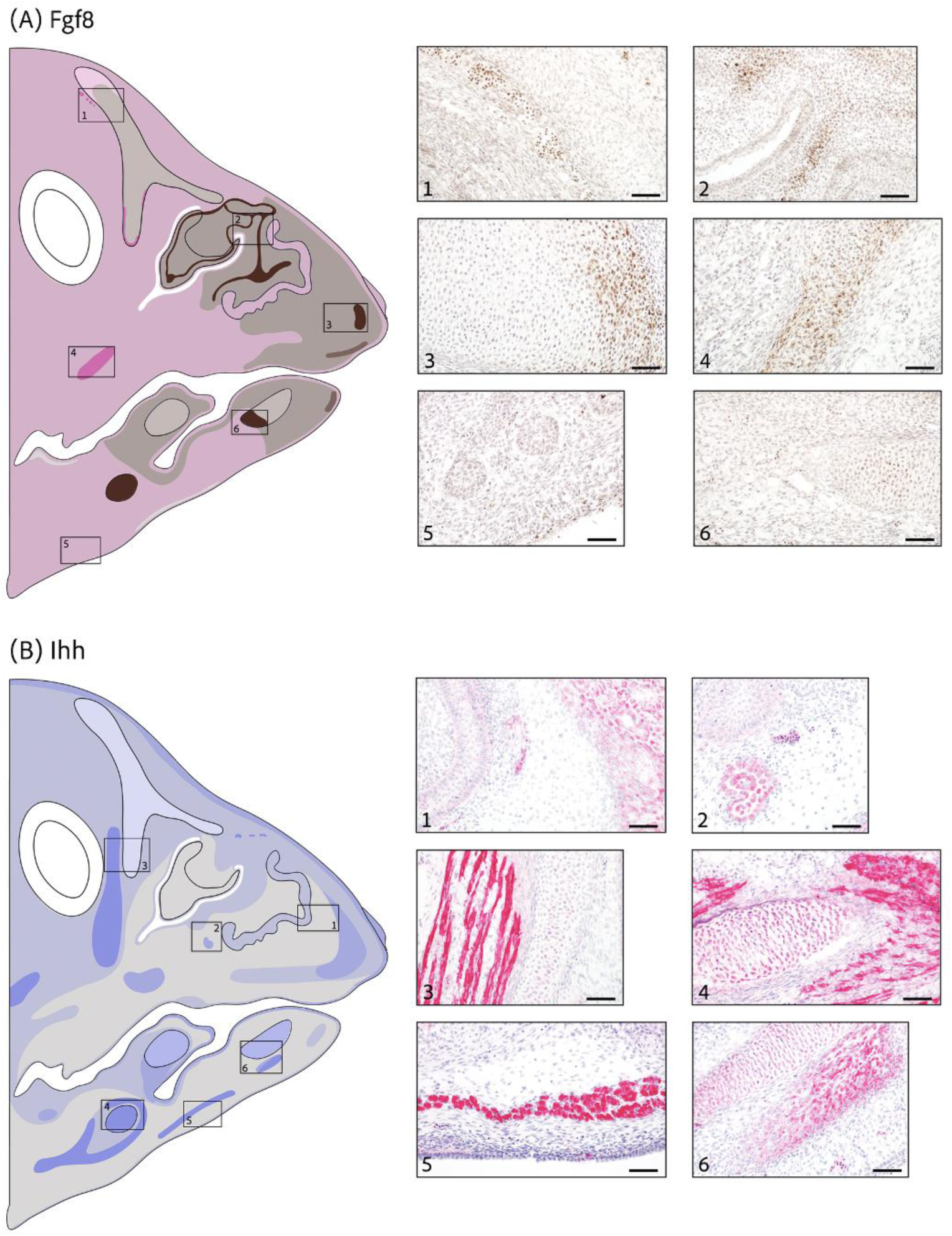
Fgf8 and Ihh expression at stage HH36. (A) **Fgf8** (brown stain) shows overall weak cytoplasmic and nuclear expression while showing only nuclear expression at the tip. (1) Low nuclear expression in developing nasal cartilage and blood cells with high cytoplasmic and nuclear expression. (2) Weak cytoplasmic and nuclear expression in epithelium surrounding nasal cavity and nasal epithelium with combination of weak and strong nuclear expression in adjacent developing cartilage. (3) Low and high nuclear expression in developing premaxillary cartilage. (4) Developing cartilage with strong cytoplasmic and nuclear expression while surrounding muscle and mesenchyme show weak expression. (5) Developing salivary glands. (6) Strong nuclear expression in Meckel’s cartilage with weak nuclear to nuclear and cytoplasmic expression in surrounding mesenchyme. (B) **Ihh** (red stain) shows weak cytoplasmic expression in top region of the upper jaw with more localized expression in the lower jaw. (1) Low expression in nasal conchae and medium expression in developing premaxillary cartilage with no expression in adjacent mesenchyme. (2) Moderate cytoplasmic expression in emerging osteogenic region and weak expression in epithelium surrounding nasal cavity but no expression in mesenchyme. Strong cytoplasmic expression in muscle (3) adjacent to nasal cartilage, which shows weak expression, (4) surrounding basihyal cartilage, with the latter showing moderate expression and (5) next to epithelium. (6) Moderate cytoplasmic expression in developing Meckel’s cartilage with absent to moderate to high expression in surrounding mesenchyme. Scale bars are 50*μ*m. (Back to Table S1)

**Figure S13.**
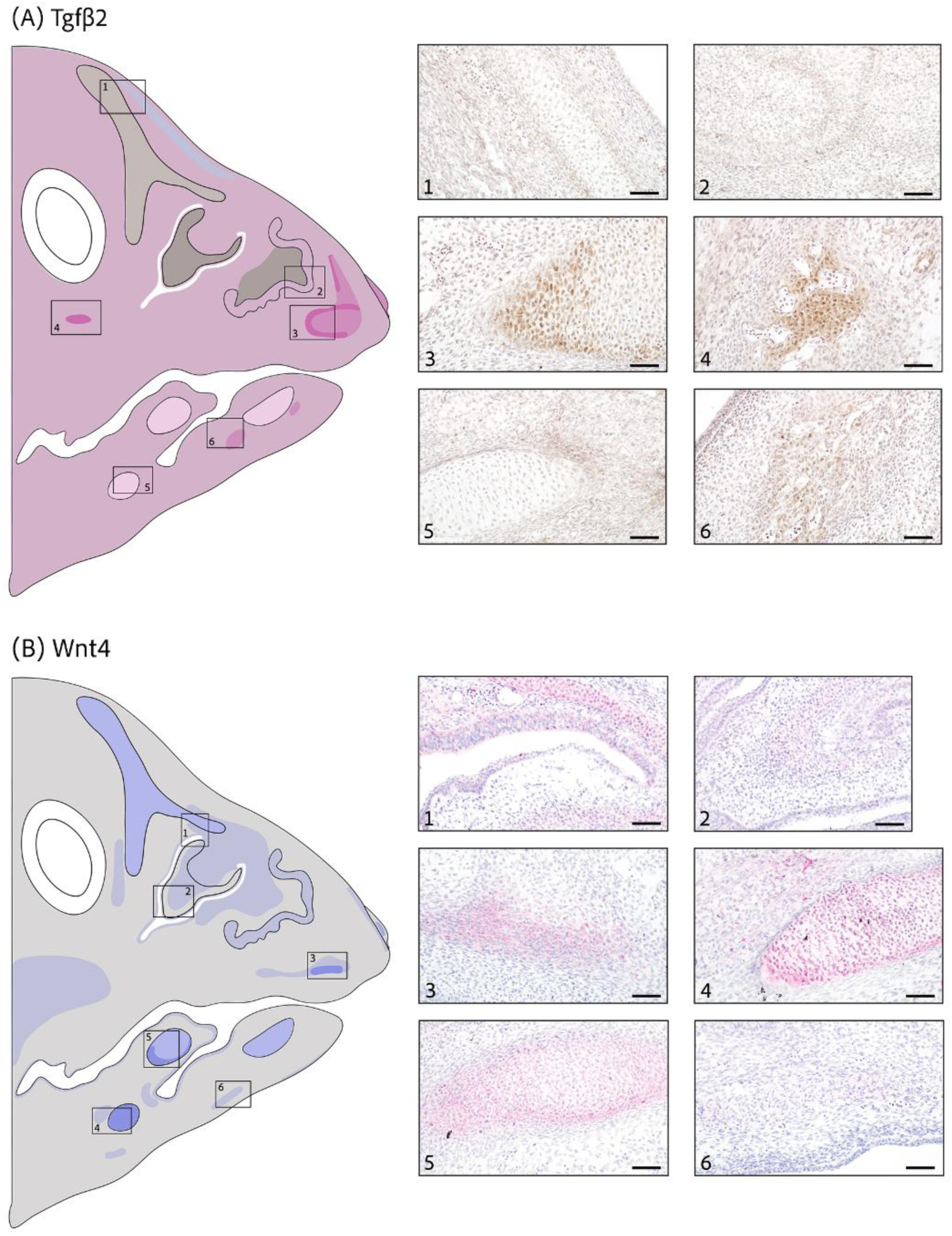
Tgfβ2 and Wnt4 expression at stage HH36. (A) **Tgfβ2** (brown stain) shows overall weak cytoplasmic and perinuclear expression. (1) Weak perinuclear expression in nasal cartilage while cytoplasmic and cytoplasmic and perinuclear expression in surrounding mesenchyme. (2) Low cytoplasmic and perinuclear expression in nasal conchae and low perinuclear expression in adjacent mesenchyme. (3) Moderate to high cytoplasmic and perinuclear expression in developing premaxillary cartilage. Emerging osteogenic region with (4) high and (6) moderate cytoplasmic and perinuclear expression. (5) Low expression in basihyal cartilage and surrounding mesenchyme. (B) **Wnt4** (red stain) is locally expressed with (1) moderate cytoplasmic expression in intermediate region of the nasal cartilage and weak expression in surrounding mesenchyme and nasal epithelium. (2) Nasal cartilage and nasal epithelium show low expression while expression is absent in adjacent mesenchyme. Strong cytoplasmic expression in (3) edge of developing premaxillary cartilage and (4) basihyal cartilage with low expression in adjacent mesenchyme. (5) Paraglossal cartilage with medium expression and high expression in perichondrogenic area. (6) Weak to no expression in mesenchyme. Scale bars are 50*μ*m. (Back to Table S1)

**Figure S14.**
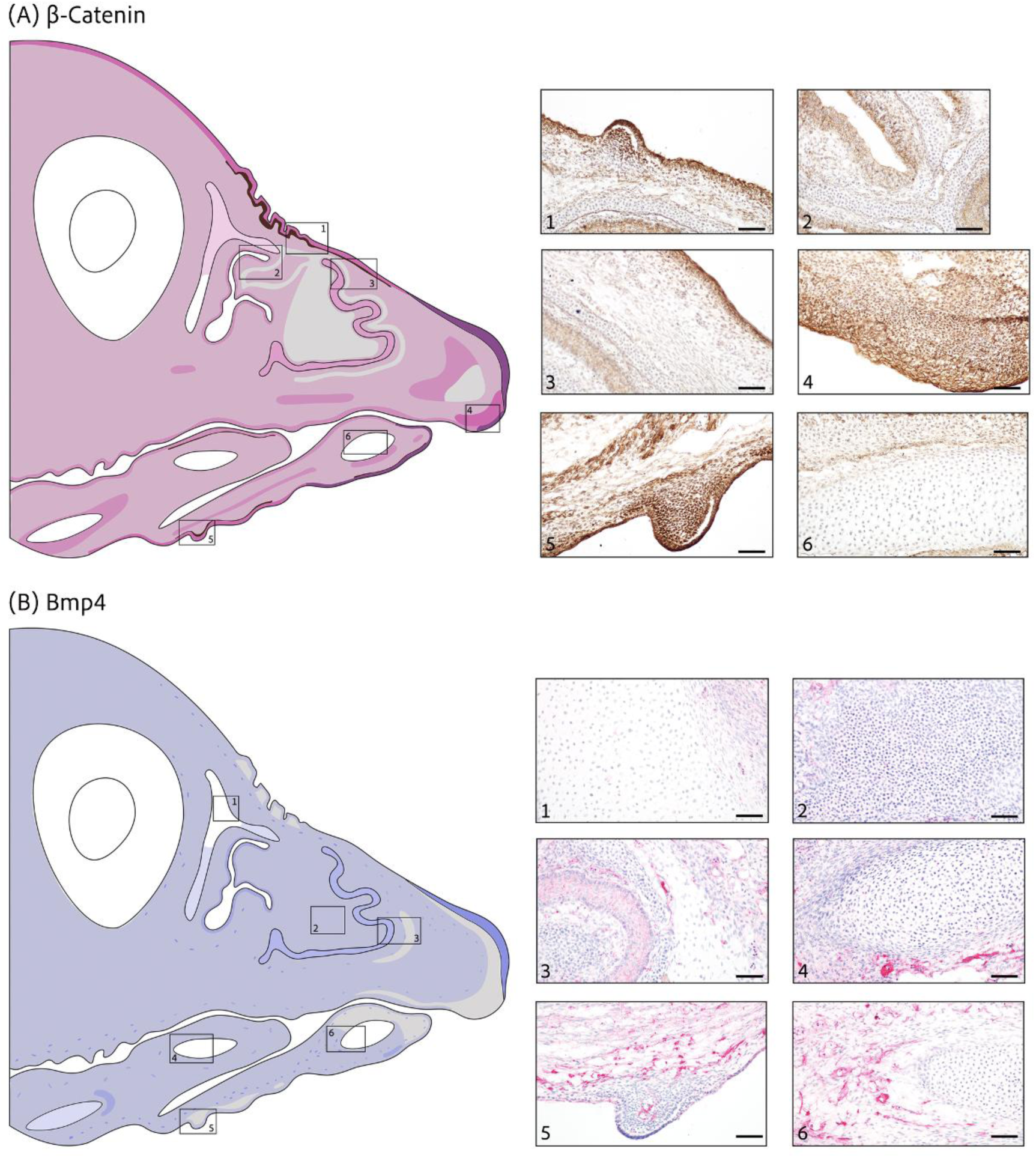
β-catenin and Bmp4 expression at stage HH39. (A) **β-catenin** (brown stain) shows overall weak cytoplasmic and nuclear expression in the mesenchyme and moderate to strong expression in the epithelium. (1, 5) Enriched nuclear expression in mesenchyme subjacent to epithelial prominences. (2, 3) Moderate expression in nasal epithelium and conchae with no expression in nasal cartilage and weak expression in surrounding mesenchyme. (4) Strong cytoplasmic, membranal and nuclear expression in epithelium at the tip with strong cytoplasmic and nuclear expression in overlaying mesenchyme. (6) Meckel’s cartilage shows no expression. (B) **Bmp4** (red stain) widely distributed weak cytoplasmic expression except for absence of expression at the tip. No expression in (1) upper region of nasal, (4) paraglossal or (6) Meckel’s cartilage, while (2) nasal cartilage and surrounding mesenchyme shows weak cytoplasmic expression. (3) Nasal conchae with moderate cytoplasmic expression and weak cytoplasmic expression in adjacent developing cartilage. (5) No expression in mesenchyme accumulated in the epithelial prominence. Blood cells show moderate to high cytoplasmic expression. Scale bars are 50*μ*m. (Back to Table S1)

**Figure S15.**
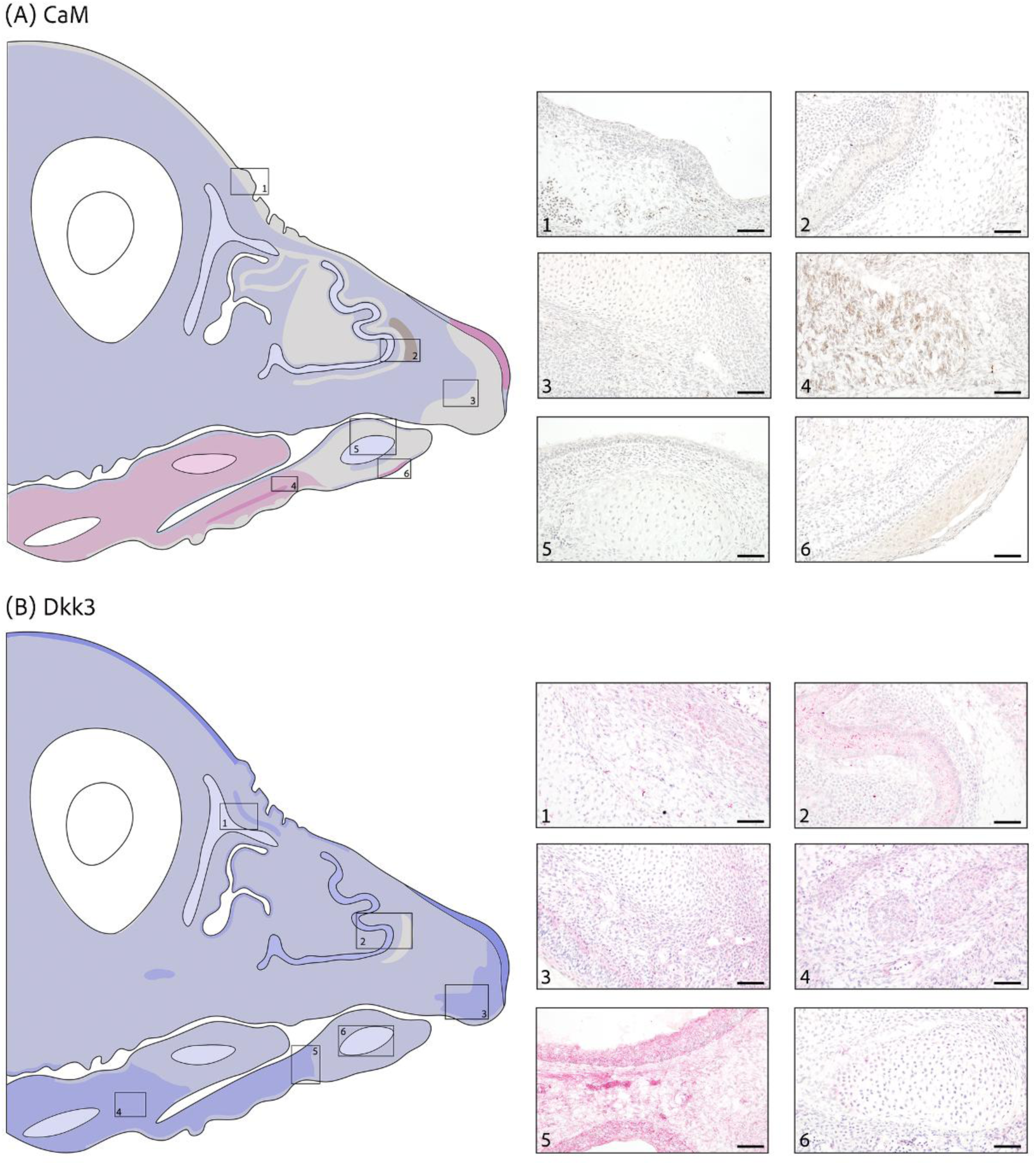
CaM and Dkk3 expression at stage HH39. (A) **CaM** (brown stain) shows weak cytoplasmic expression in the upper jaw while weak cytoplasmic and nuclear expression in the lower jaw with no expression at the tip. Difference in expression between surface epithelium and oral epithelium, with surface epithelium lacking expression and oral epithelium showing weak cytoplasmic expression. (1) No expression in mesenchyme subjacent to epithelial prominences. (2) Weak cytoplasmic expression in nasal conchae and adjacent cartilage, but weak nuclear expression in nearby mesenchyme. (3) Developing premaxillary cartilage and (5) Meckel’s cartilage shows weak cytoplasmic expression with no expression in adjoining mesenchyme. (4) Myogenic area and (6) epithelium with high cytoplasmic and nuclear expression. (B) **Dkk3** (red stain) shows weak cytoplasmic expression in upper jaw and moderate cytoplasmic expression at the lower jaw. (1) Low cytoplasmic expression in nasal cartilage and adjacent mesenchyme with moderate expression in emerging osteogenic region. (2) Medium cytoplasmic expression in nasal conchae with weak expression in adjacent cartilage and no expression in nearby mesenchyme. Weak cytoplasmic expression in (3) premaxillary and (4) primordia epithelial cords with moderate expression in adjacent mesenchyme. (5) Medium to low expression in mesenchyme and epithelium. (6) Meckel’s cartilage shows low cytoplasmic expression. Scale bars are 50*μ*m. (Back to Table S1)

**Figure S16.**
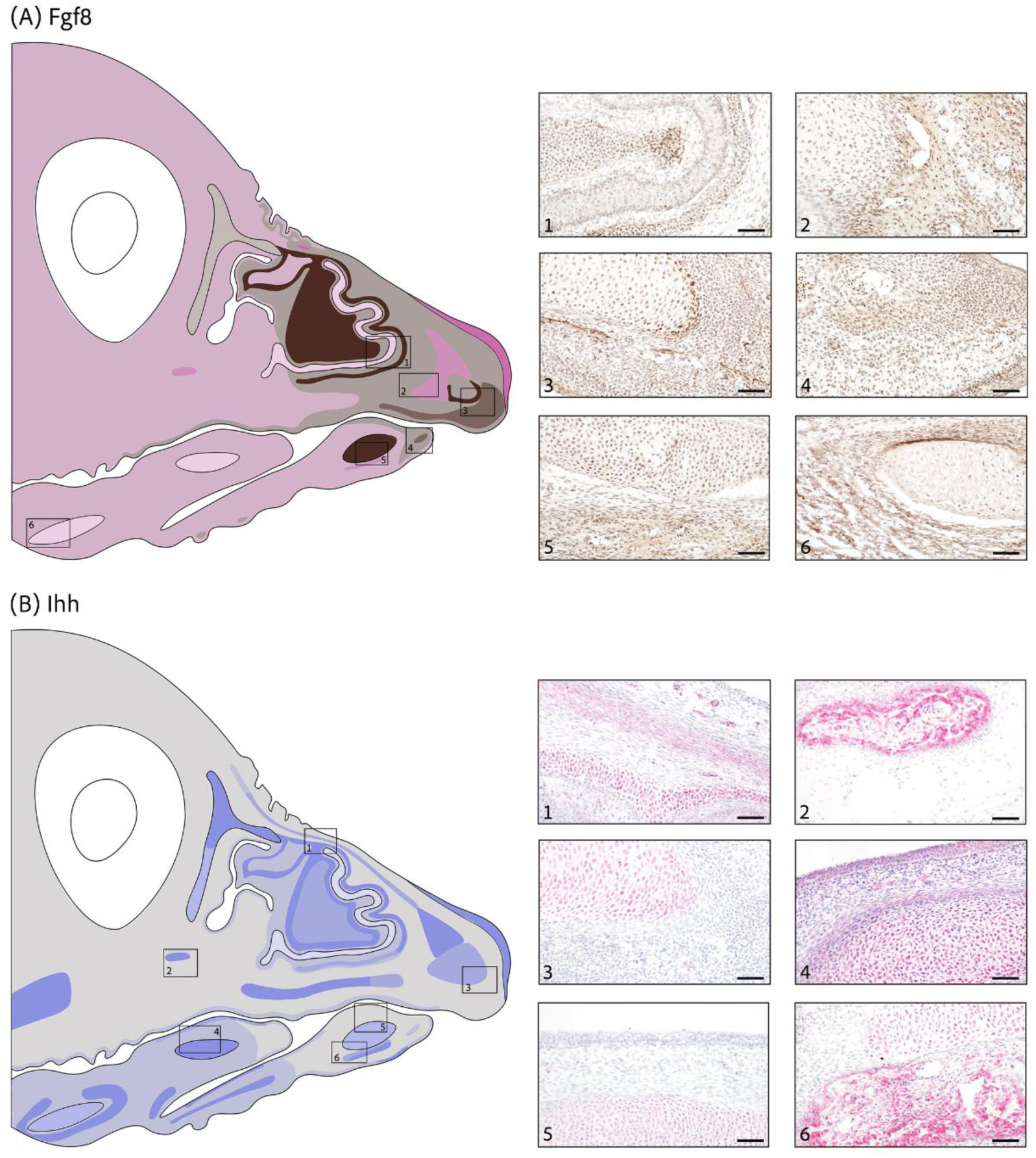
Fgf8 and Ihh expression at stage HH39. (A) **Fgf8** (brown stain) shows weak cytoplasmic and nuclear expression with only nuclear expression at the tip. (1) Nasal conchae with weak cytoplasmic and nuclear expression, strong nuclear expression in nasal cartilage and low nuclear expression in surrounding mesenchyme. (2) Moderate cytoplasmic and nuclear expression in emerging osteogenic region and low nuclear expression in adjacent mesenchyme. (3) Premaxillary cartilage with low nuclear expression, perichondrogenic area shows strong expression and moderate expression in subjacent mesenchyme. (4) Weak to moderate expression in mesenchyme. (5) Strong nuclear expression in Meckel’s cartilage and moderate cytoplasmic and nuclear expression in emerging osteogenic region. (6) Basihyal cartilage and surrounding mesenchyme shows weak cytoplasmic and nuclear expression. (B) **Ihh** (red stain) is locally expressed in upper jaw with more widely distributed weak cytoplasmic expression in the lower jaw. (1) Nasal cartilage with high cytoplasmic expression, no expression in adjacent mesenchyme but moderate to (2) high expression in emerging osteogenic region. (3) Premaxillary and (5) Meckel’s cartilage with moderate cytoplasmic expression and no expression in surrounding mesenchyme or epithelium. (4) Developing paraglossal cartilage with high expression, low expression in mesenchymal cells and moderate expression in epithelium. (6) Moderate to high expression in developing osteogenic region adjacent to Meckel’s cartilage with medium cytoplasmic expression. Scale bars are 50*μ*m. (Back to Table S1)

**Figure S17.**
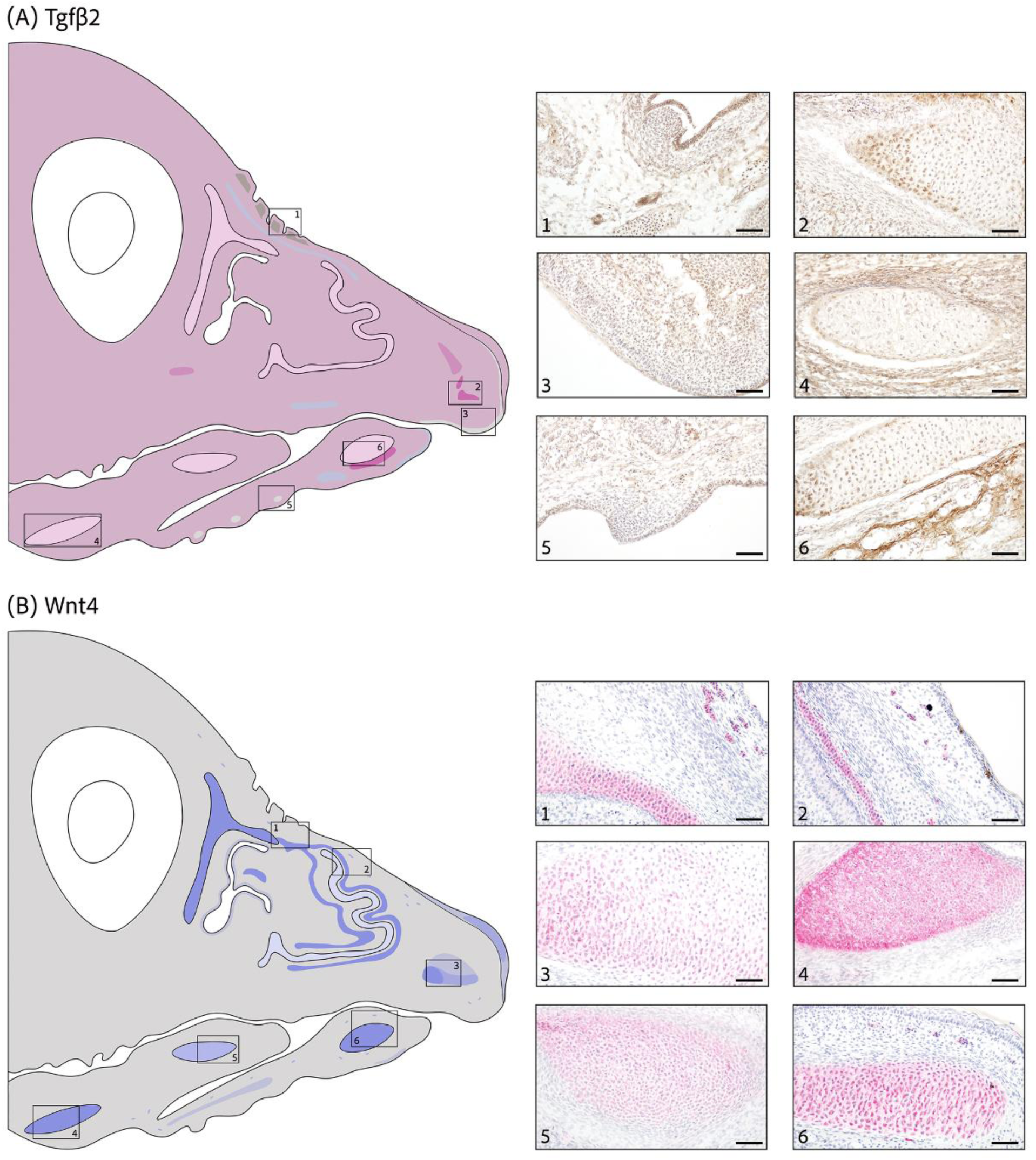
Tgfβ2 and Wnt4 expression at stage HH39. (A) **Tgfβ2** (brown stain) shows overall weak cytoplasmic and nuclear expression. Weak nuclear expression in mesenchyme subjacent to epithelial prominences in (1) upper beak and (5) no expression in lower beak. (2) Premaxillary cartilage and surrounding mesenchyme with weak cytoplasmic and nuclear expression, but moderate expression in perichondrogenic area. (3) Epithelium shows no expression and adjacent mesenchyme shows weak cytoplasmic and nuclear expression. Low expression in (4) basihyal and (6) Meckel’s cartilage and adjacent mesenchyme with moderate expression in emerging osteogenic region. (B) **Wnt4** (red stain) is highly localized. High cytoplasmic expression in (1, 2) nasal, (4) basihyal and (6) Meckel’s cartilage with no expression in surrounding mesenchyme. (3) Weak to moderate to strong cytoplasmic expression in premaxillary cartilage. (5) Developing paraglossal shows medium cytoplasmic expression. Blood cells with medium cytoplasmic expression. Scale bars are 50*μ*m. (Back to Table S1)

**Figure S18.**
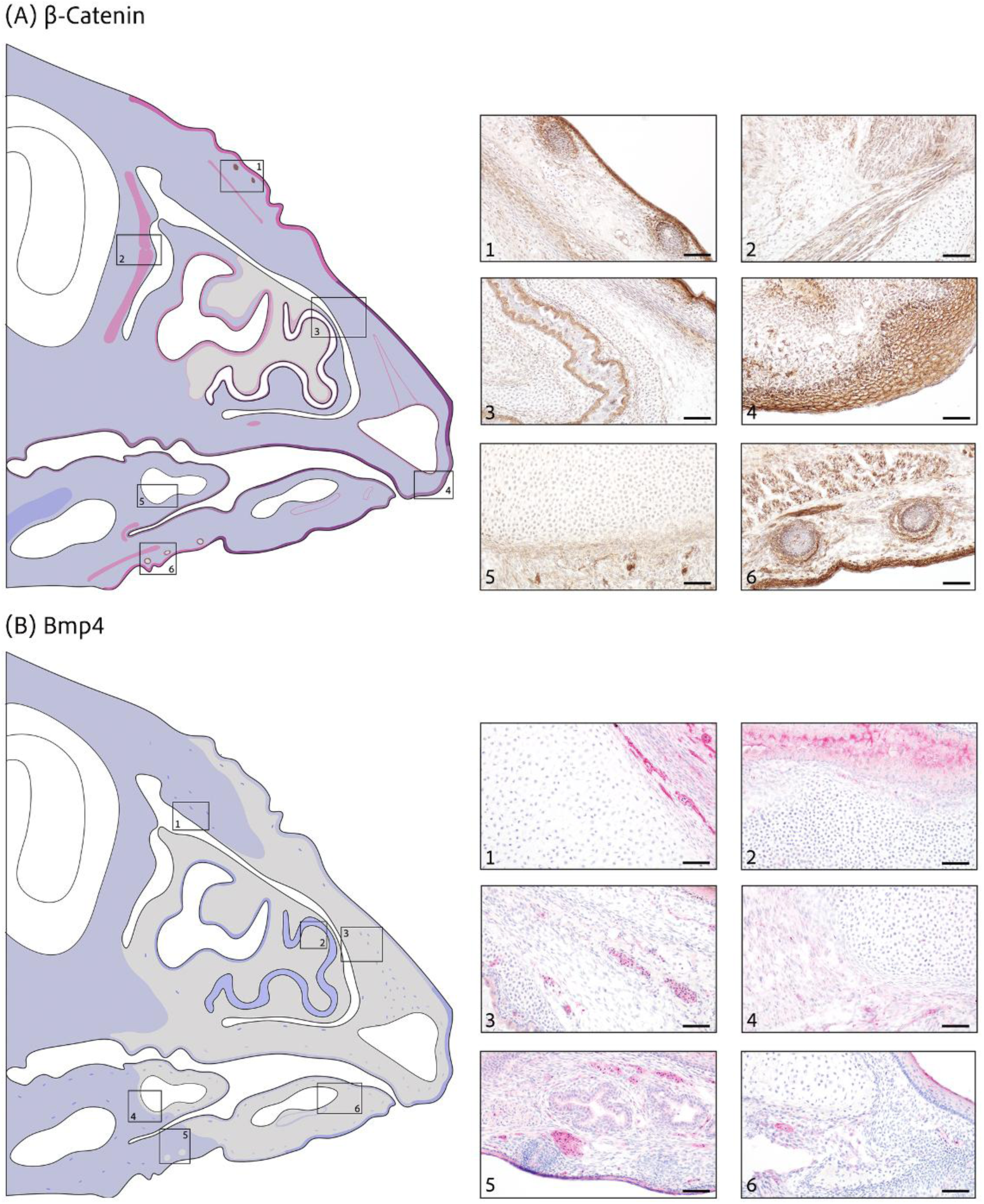
β-catenin and Bmp4 expression at stage HH42. (A) **β-catenin** (brown stain) shows overall weak cytoplasmic expression with moderate to high cytoplasmic and nuclear or cytoplasmic, nuclear and membranal epithelial expression. (1) Medium cytoplasmic and nuclear expression in emerging osteogenic region and border of dermal condensation, epithelium with strong expression. Moderate nuclear expression in dermal condensation. (2) No expression in nasal cartilage and medium cytoplasmic and nuclear expression in adjacent myogenic area. (3) Cartilage with no expression, weak cytoplasmic and nuclear expression in perichondrogenic area and strong cytoplasmic, nuclear and membranal expression in epithelium surrounding nasal conchae. (4) Medium cytoplasmic, nuclear and membranal expression in epithelium at the tip, weak cytoplasmic and nuclear expression in perichondrogenic area and only weak cytoplasmic expression in surrounding mesenchyme. (5) Paraglossal cartilage with no expression and weak cytoplasmic expression in mesenchymal cells. (6) Moderate cytoplasmic and nuclear expression in muscle and outer layer of dermal condensation followed by moderate nuclear and no expression. (B) **Bmp4** (red stain) shows overall weak cytoplasmic expression and no expression at the tip. Blood cells with weak to strong cytoplasmic expression. (1) Nasal cartilage with no expression and weak cytoplasmic expression in surrounding mesenchyme. (2) Moderate expression in nasal conchae but no expression in surrounding cartilage. (3) Emerging osteogenic region and mesenchyme show no expression, but pockets of blood cells show medium cytoplasmic expression. (4) Paraglossal cartilage with no expression and weak cytoplasmic expression in myogenic area. (5) Low cytoplasmic expression in mesenchyme and salivary glands but no expression in dermal condensations. (6) Meckel’s cartilage and surrounding mesenchyme shows no expression, epithelium and periosteogenic area shows weak cytoplasmic expression. Scale bars are 50*μ*m. (Back to Table S1)

**Figure S19.**
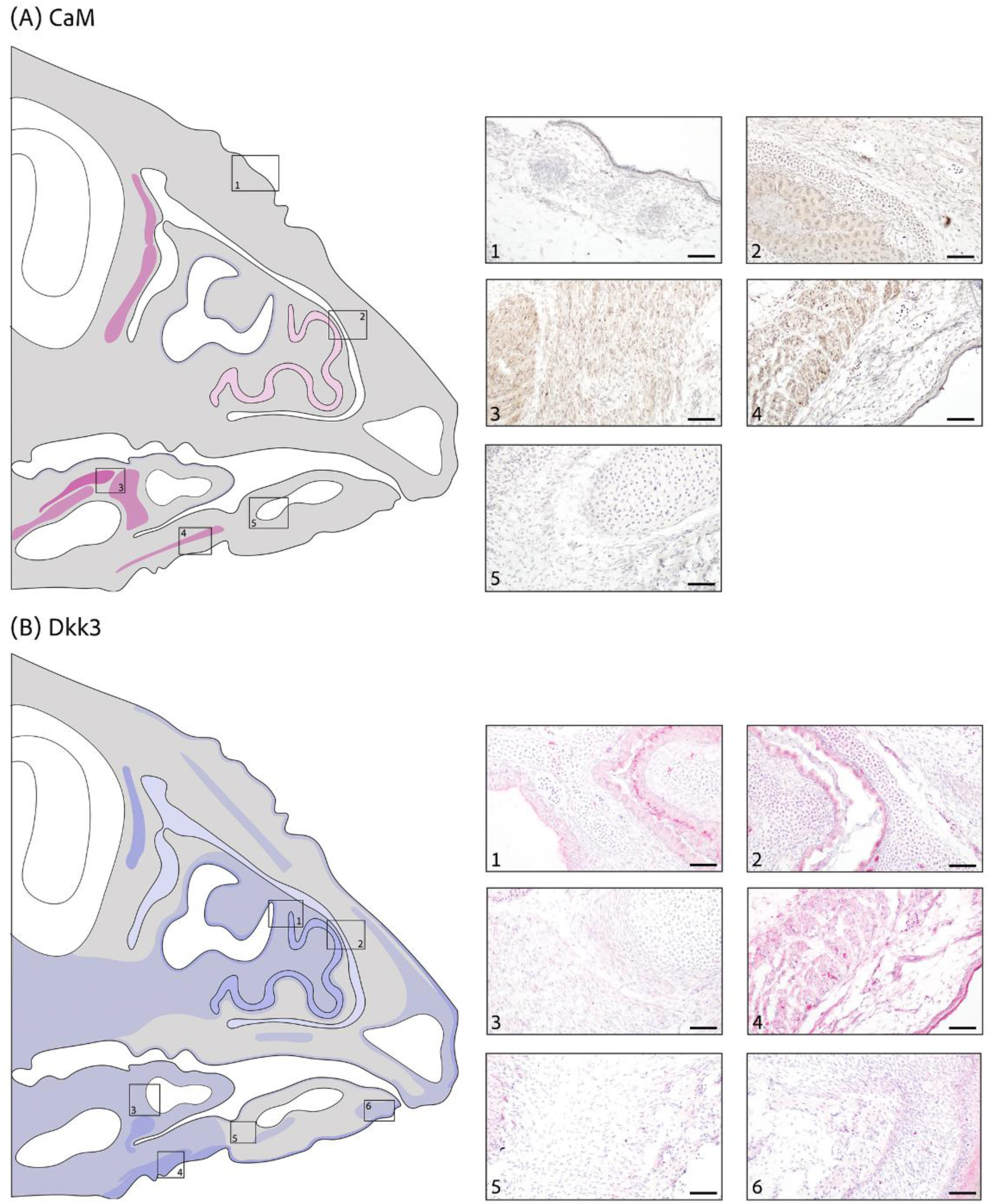
CaM and Dkk3 expression at stage HH42. (A) **CaM** (brown stain) is locally expressed but shows no expression in mesenchyme of upper and lower jaw. (1) Dermal condensations, surrounding mesenchyme and epithelium show no expression. (2) Weak cytoplasmic and nuclear expression in nasal conchae. (3, 4) Moderate to strong cytoplasmic and nuclear expression in myogenic area. (5) Meckel’s cartilage and mesenchyme with no expression. (B) **Dkk3** (red stain) shows moderate cytoplasmic expression in (1) nasal epithelium and (1, 2) nasal conchae with weak expression in adjacent cartilage and no expression in mesenchyme. (3) Paraglossal cartilage with no expression and myogenic area with weak cytoplasmic expression. (4) Muscle and adjacent epithelium with medium cytoplasmic expression. (5, 6) Weak to moderate cytoplasmic expression in epithelium, weak expression in emerging osteogenic region, weak to no expression in surrounding mesenchyme. Scale bars are 50*μ*m. (Back to Table S1)

**Figure S20.**
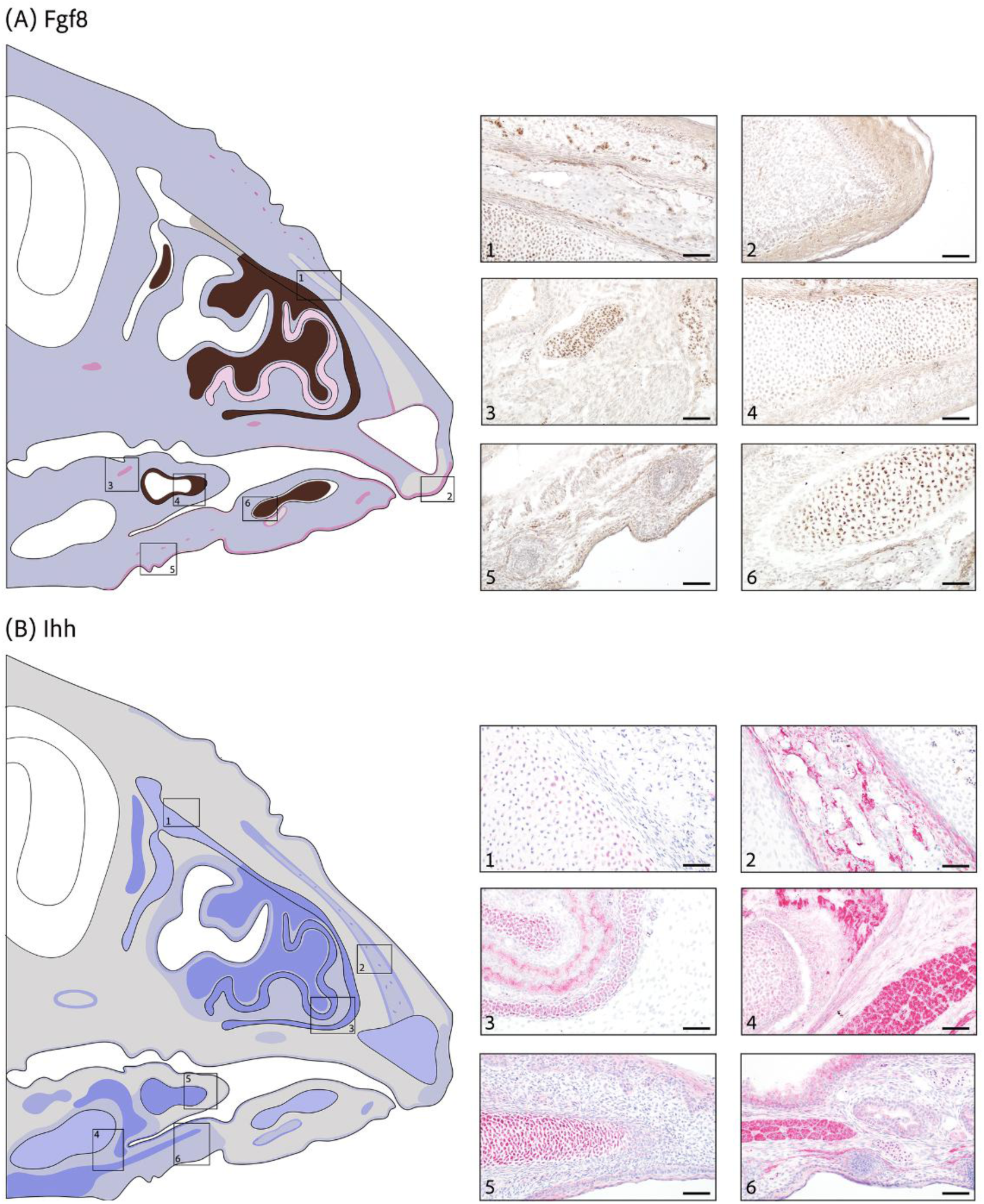
Fgf8 and Ihh expression at stage HH42. (A) **Fgf8** (brown stain) is weakly expressed in the cytoplasm across the upper and lower jaw. (1) Strong nuclear expression in cartilage with no expression in osteogenic area and weak cytoplasmic expression in periosteogenic area and mesenchyme. (2) Moderate cytoplasmic and nuclear expression in epithelium but no expression in overlaying mesenchyme. (3) Blood cells with high cytoplasmic and nuclear expression. (4) No expression in paraglossal cartilage with strong nuclear expression in cartilage edges. (5) Weak cytoplasmic expression in dermal condensations, mesenchyme and muscle with medium cytoplasmic and nuclear expression in epithelium. (6) Meckel’s cartilage shows strong nuclear expression but no expression in cartilage edges and osteogenic region. Periosteogenic region with moderate cytoplasmic and nuclear expression. (B) **Ihh** (red stain) shows almost no expression in mesenchyme of upper and lower jaw. (1) Nasal cartilage with medium cytoplasmic expression. (2) Weak cytoplasmic expression in osteogenic region with moderate expression in surrounding periosteogenic areas. (3) Nasal conchae and surrounding cartilage show strong cytoplasmic expression and weak cytoplasmic expression in adjacent mesenchyme. (4) Basihyal cartilage with moderate cytoplasmic expression, weak cytoplasmic expression in surrounding mesenchyme and strong expression in muscle tissue. (5) Paraglossal cartilage with strong cytoplasmic expression and weak expression in epithelium and perichondrogenic area. (6) Strong cytoplasmic expression in muscle and inner epithelium. Weak expression in dermal condensations, mesenchyme and outer epithelium. Scale bars are 50*μ*m. (Back to Table S1)

**Figure S21.**
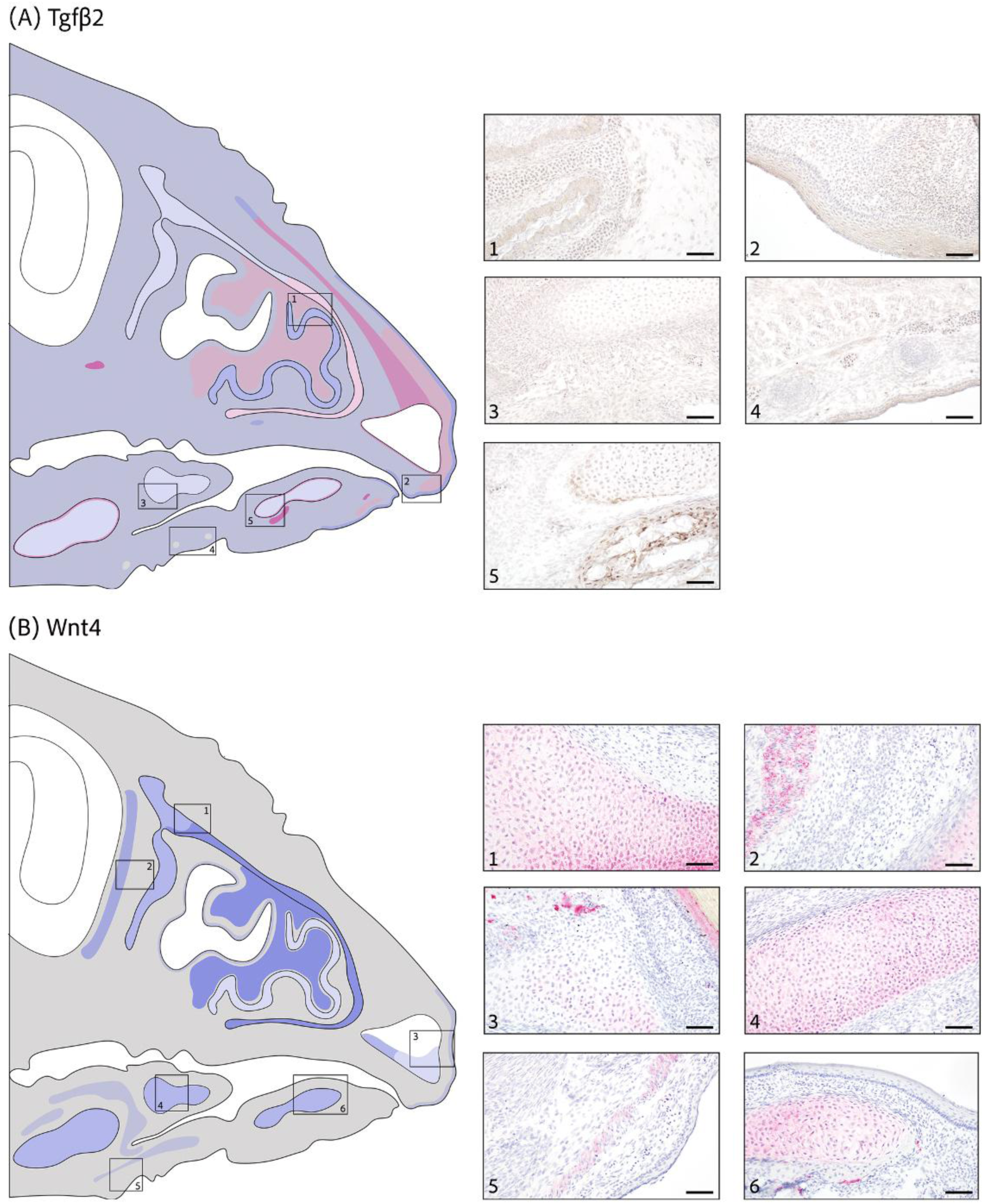
Tgfβ2 and Wnt4 expression at stage HH42. (A) **Tgfβ2** (brown stain) shows overall weak cytoplasmic expression with weak cytoplasmic and perinuclear expression in upper jaw tip. (1) Moderate cytoplasmic expression in epithelium surrounding nasal conchae. Weak cytoplasmic and perinuclear expression in adjacent cartilage. (2) Mesenchyme with weak cytoplasmic or cytoplasmic and perinuclear expression. Low expression in (3,4) paraglossal cartilage, muscle and adjacent mesenchyme and epithelium. (4) Dermal condensations show no expression. (5) Meckel’s cartilage and surrounding mesenchyme show weak cytoplasmic expression. Moderate cytoplasmic and nuclear expression in perichondrogenic area and strong expression in adjacent osteogenic region. (B) **Wnt4** (red stain) shows limited expression with no expression in mesenchyme. (1, 2) Nasal cartilage with moderate to strong cytoplasmic expression and (2) moderate expression in adjacent muscle. (3) Premaxillary cartilage with weak to no cytoplasmic expression and weak cytoplasmic expression in adjacent epithelium. Moderate cytoplasmic expression in (4) paraglossal and (6) Meckel’s cartilage. (5) Muscle shows weak cytoplasmic expression. Scale bars are 50*μ*m. (Back to Table S1)

**Figure S22.**
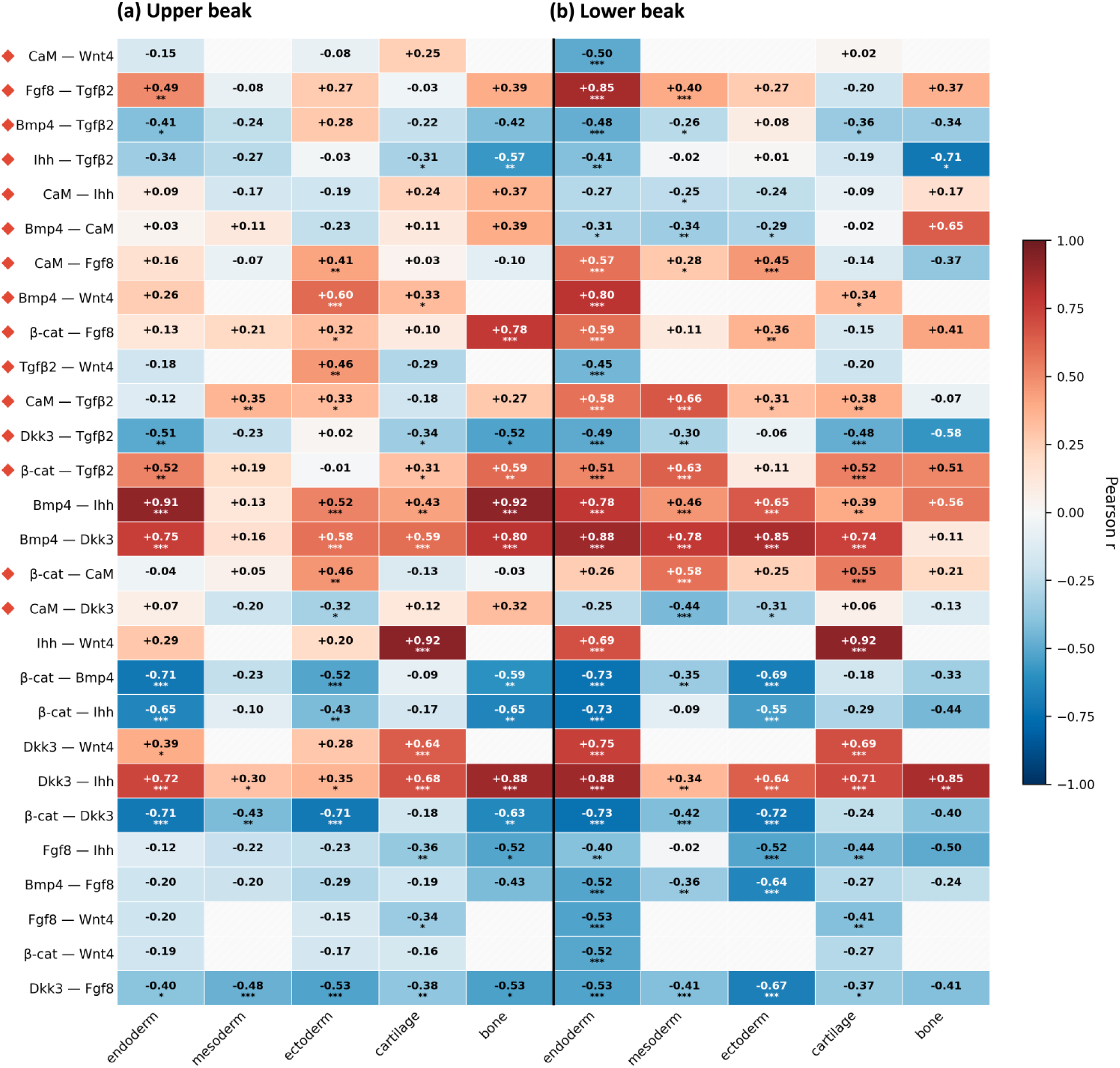
**Non-overlapping tissue-specific regulatory states of the core network (Fig. 2)** in **(a)** upper and **(b)** lower beak. Protein pairs are sorted from top to bottom by the range of their tissue-type variability (indicated by red squares). Shown are tissue types shared between jaws and Pearson *r*’s (\**P* < 0.1, \*\**P* < 0.05, \*\*\**P* < 0.01). Grid cells with missing values have <5 observations.

**Figure S23.**
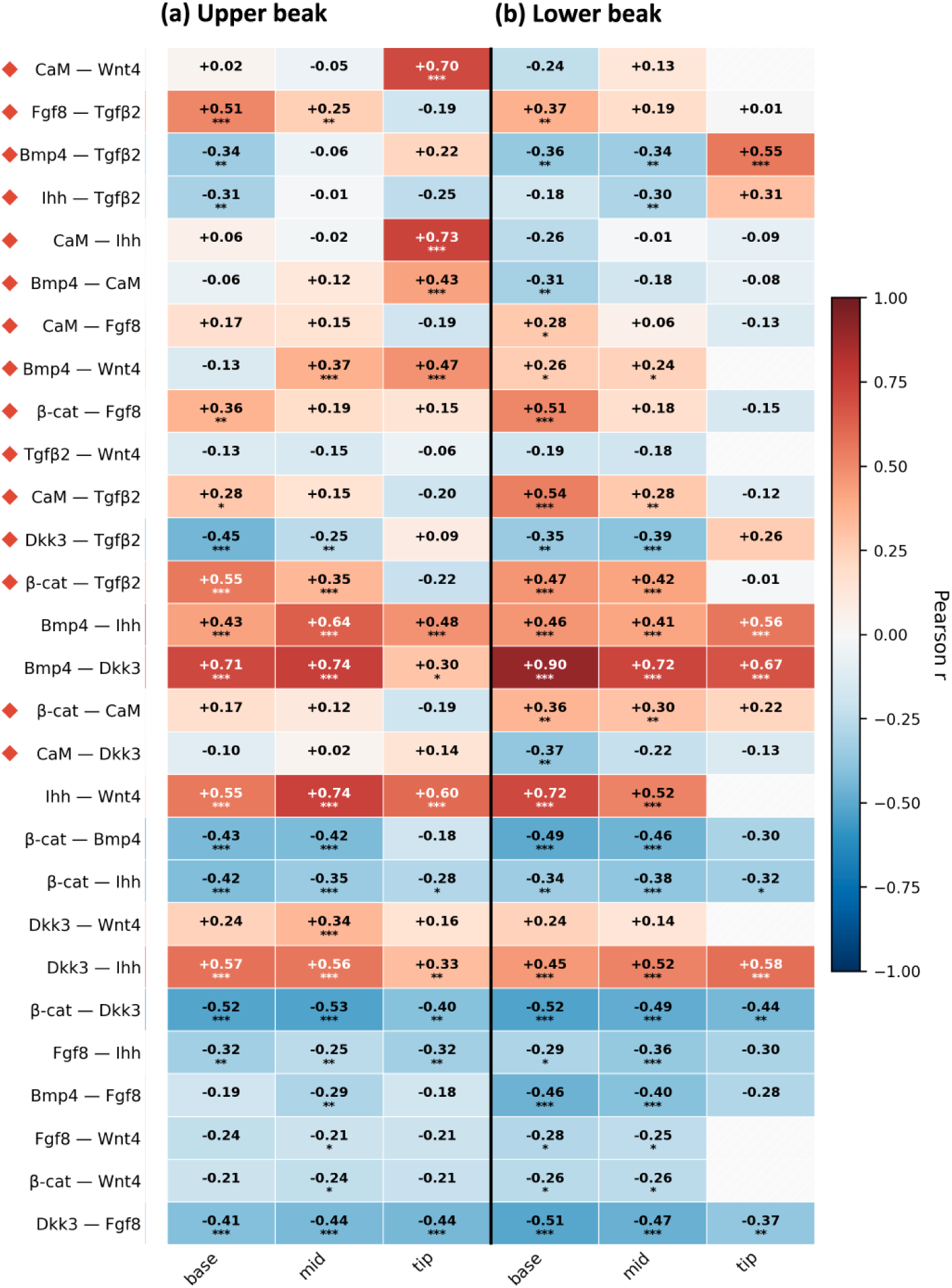
**Non-overlapping location-specific regulatory states of the core network (Fig. 2)** in **(a)** upper and **(b)** lower beak. Protein pairs are sorted from top to bottom by the range of their location-specific variability (red squares). Shown are the locations shared between jaws and Pearson *r*’s (\**P* < 0.1, \*\**P* < 0.05, \*\*\**P* < 0.01). Grid cells with missing values have <5 observations.

**Figure S24.**
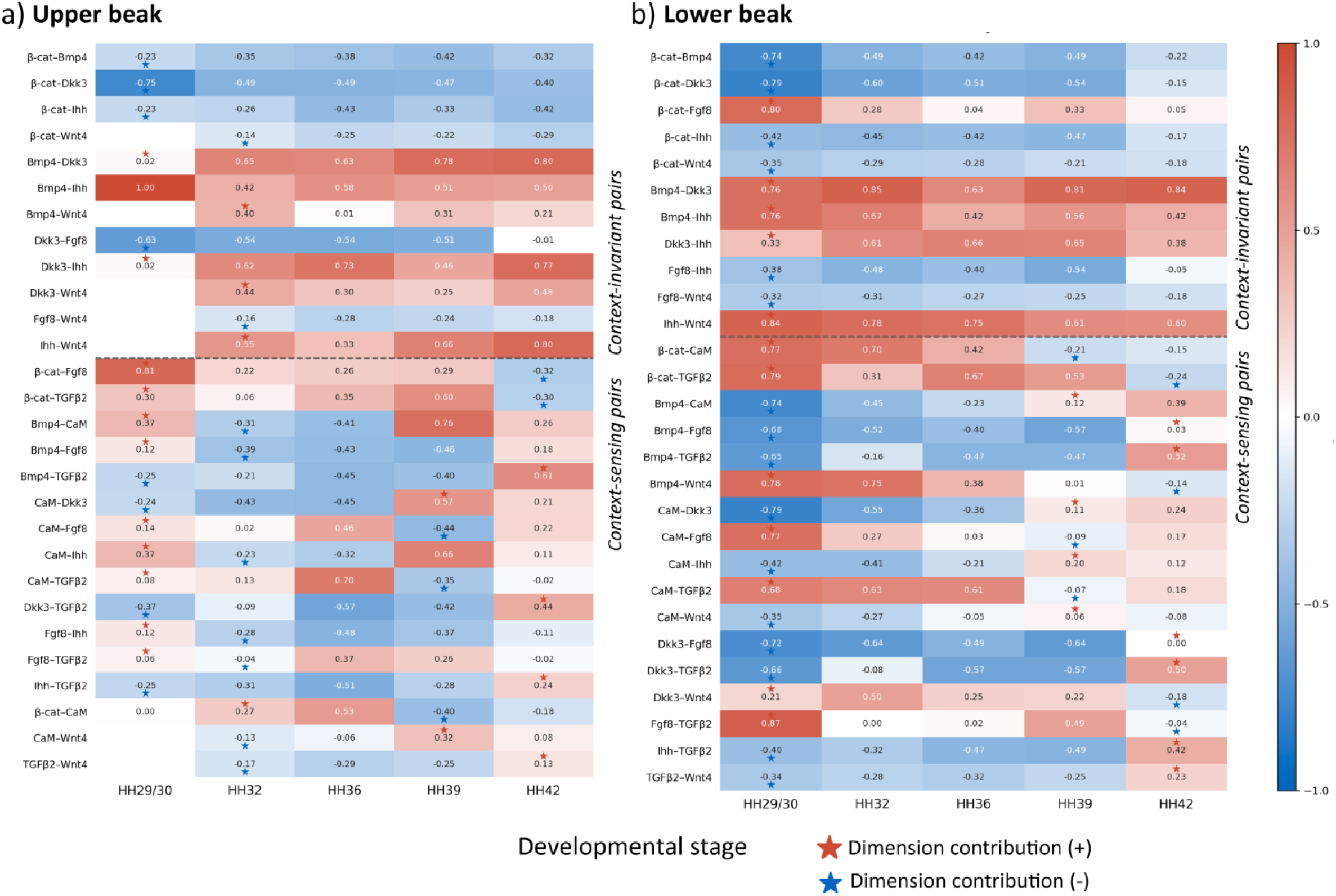
Context-sensing proteins drive non-redundant regulatory space expansion in bow-tie network. Temporal patterns of exploration of controllable dimensions in the network. Heatmaps show stage-specific Pearson *r*’s by developmental stage for **(a)** upper and **(b)** lower beak. Within the context-sensing protein group, rows are ordered by the stage at which the first **new (non-redundant)** controllable dimension is being added to the cumulative reachable space (indicated by a star). Red star shows the earliest stage at which a pair’s positive-sign dimension enters the controllable space; blue star – where negative-sign dimension enters. Context-sensing protein pairs (especially involving CaM and Tgfβ2) drive both positive and negative dimensions across stages. HH42 stage marks the largest expansion of controllable dimensions, reflecting a late-stage network reconfiguration centered on Tgfβ2 (Bmp4–Tgfβ2 and β-cat– Tgfβ2 both reverse signs at HH42).

**Figure S25.**
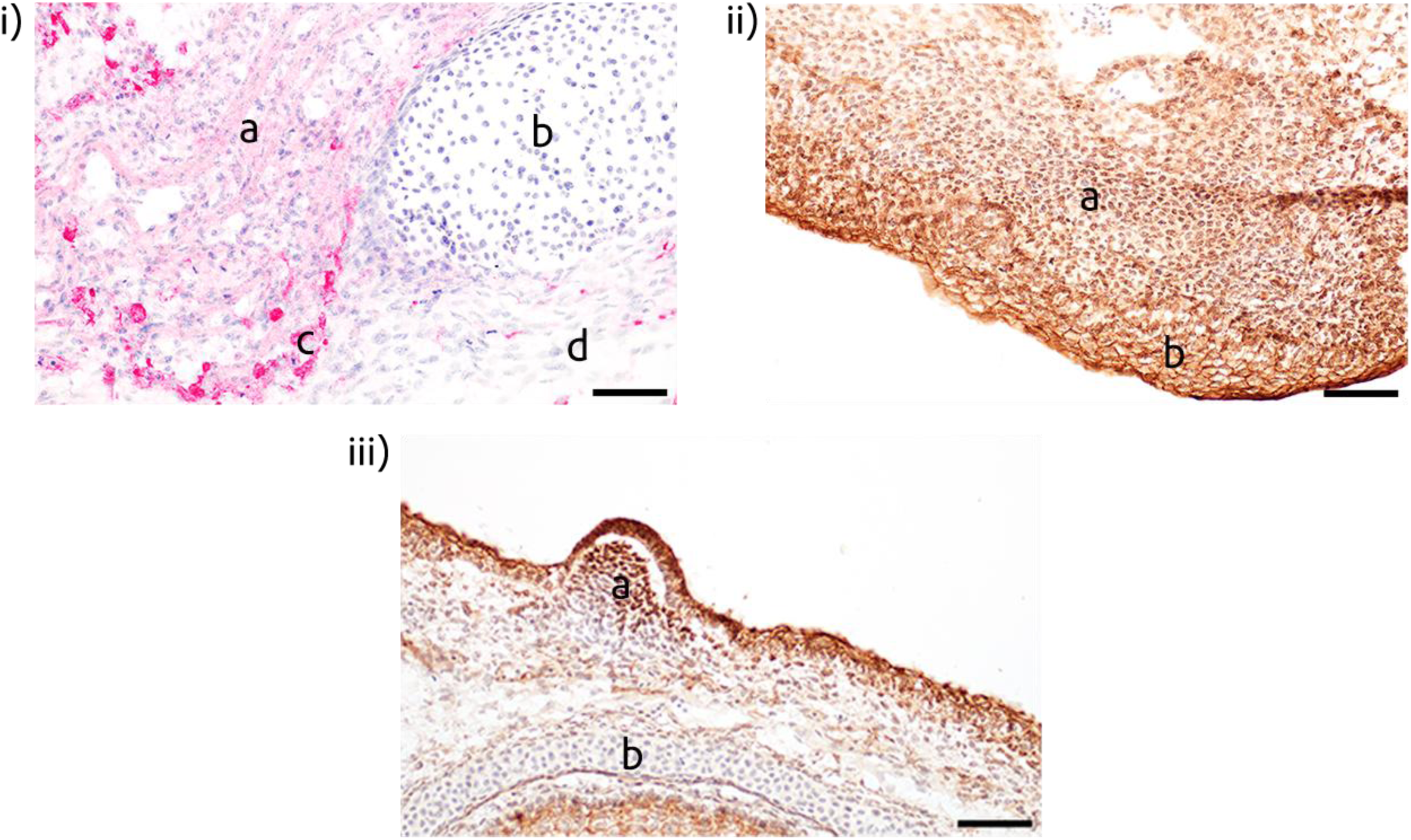
Examples of protein expression categorization in different subcellular locations. i) Vector Red staining for Bmp4 at (d) low, (a) medium and (c) high cytoplasmic expression and (b) absence of expression. ii) DAB staining for β-catenin at (a) strong nuclear and cytoplasmic expression and (b) strong nuclear, cytoplasmic and membranal expression. iii) DAB staining for β-catenin at (a) strong nuclear expression and (b) absence of expression. Scale bars are 50*μ*m.

**Figure S26.**
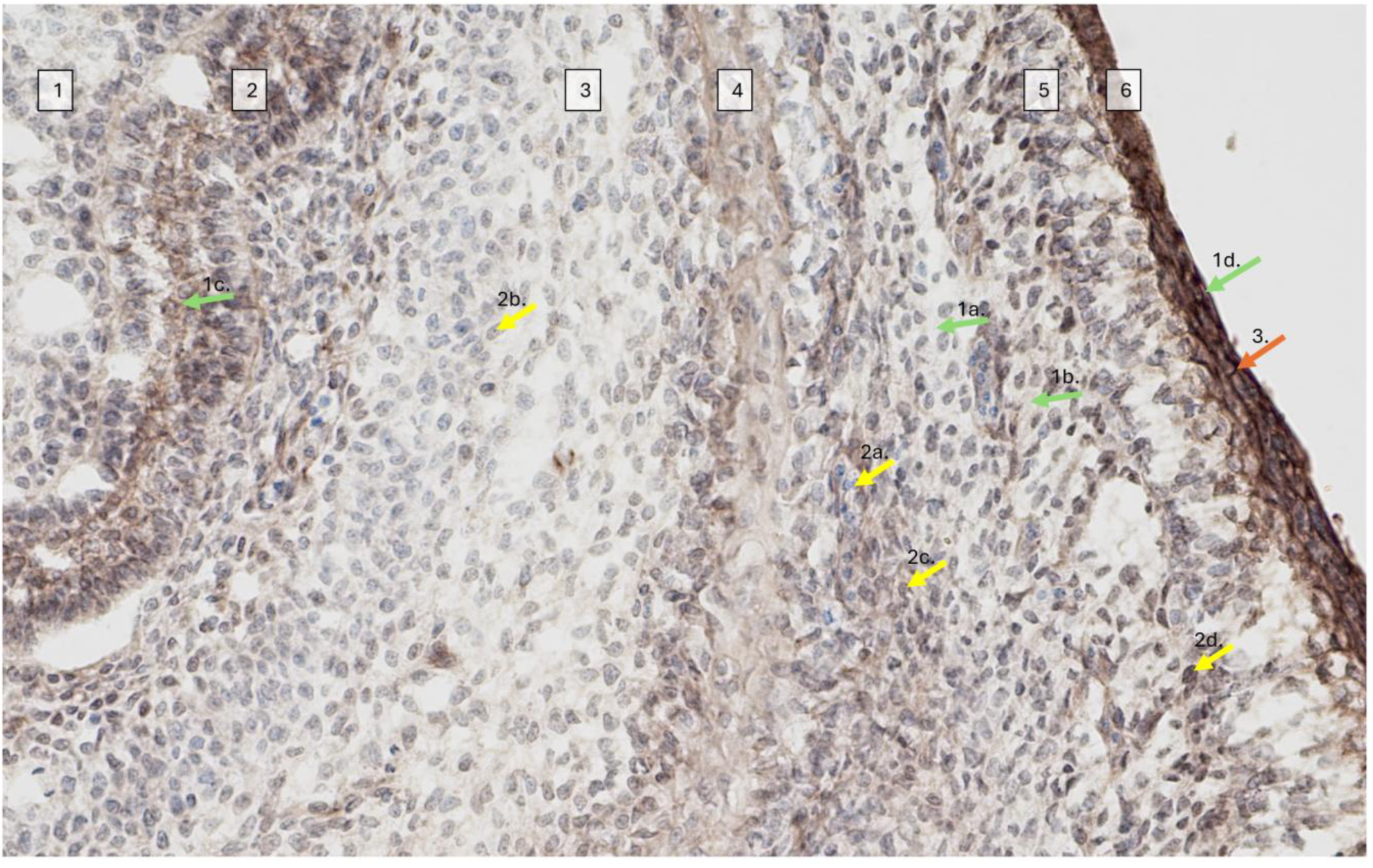
Example illustrating protein expression intensity measures and subcellular location using β-catenin in the upper beak at HH 36. White boxes overlaying tissue types: Box 1: Conchae; Box 2: Nasal epithelium; Box 3: Cartilage; Box 4: Osteogenic region (bone); Box 5: Mesenchyme; Box 6: Surface epithelium. **<u>1) Green arrows showing cytoplasmic staining</u>** 1a. Clear, no staining (0) 1b. Light tan, light staining (1) 1c. Medium brown, medium staining (2) 1d. Dark brown, strong staining (3) **<u>2. Yellow arrows showing nuclear staining</u>** 2a. Bright blue, no staining (0) 2b. Dull blue-gray, light staining (1) 2c. Uniform gray, medium staining (2) 2d. Dark nearly black, strong staining (3) <u>3.</u> Orange arrow showing strong (3) membranal as black lines

## References

1. Hallgrímsson B., Aponte J.D., Vidal-García M., Richbourg H., Green R., Young N.M., Cheverud J.M., Calof A.L., Lander A.D., Marcucio R.S. 2023 The developmental basis for evolvability. In Evolvability: A Unifying Concept in Evolutionary Biology? (eds. Hansen T.F., Houle D., Pavličev M., Pélabon C.), pp. 171–198, The MIT Press.

2. Kauffman S.A. 1993 The Origins of Order: Self Organization and Selection in Evolution. Oxford, UK, Oxford University Press.

3. Davidson E.H., Erwin D.H. 2006 Gene regulatory networks and the evolution of animal body plans. Science 311, 796–800.

4. Erwin D.H., Davidson E.H. 2009 The evolution of hierarchical gene regulatory networks. Nature Reviews Genetics 10, 141–148.

5. Davidson E.H. 2006 The Regulatory Genome: Gene Regulatory Networks in Development and Evolution. San Diego, Academic Press; 304 p.

6. Csete M.E., Doyle J.C. 2002 Reverse engineering of biological complexity. Science 295(5560), 1664–1669. (doi:10.1126/science.1069981).

7. Friedlander T., Mayo A.E., Tlusty T., Alon U. 2015 Evolution of bow-tie architectures in biology. PLoS Comput Biol 11(3), e1004055. (doi:10.1371/journal.pcbi.1004055).

8. Itoh T., Kondo Y., Aoki K., Saito N. 2024 Revisiting the evolution of bow-tie architecture in signaling networks. NPJ Syst Biol Appl 10(1), 70. (doi:10.1038/s41540-024-00396-8).

9. Csete M., Doyle J. 2004 Bow ties, metabolism and disease. Trends Biotechnol 22(9), 446–450. (doi:10.1016/j.tibtech.2004.07.007).

10. Matni N., Ames A.D., Doyle J.C. 2024 A Quantitative Framework for Layered Multirate Control: Toward a Theory of Control Architecture. IEEE Control Systems 44(3), 52–94. (doi:10.1109/mcs.2024.3382388).

11. Ghosh Roy G., He S., Geard N., Verspoor K. 2021 Bow-tie architecture of gene regulatory networks in species of varying complexity. Journal of the Royal Society, Interface / the Royal Society 18(179), 20210069. (doi:10.1098/rsif.2021.0069).

12. Hilliard S., Mosoyan K., Branciamore S., Gogoshin G., Zhang A., Simons D.L., Rockne R.C., Lee P.P., Rodin A.S. 2023 Bow-tie architectures in biological and artificial neural networks: Implications for network evolution and assay design. iScience 26(2), 106041. (doi:10.1016/j.isci.2023.106041).

13. Badyaev A.V., Sánchez Moreno C., Lau B.M., Morrison E.S. 2026 Biophysical resetting of specialization drives evolutionary diversification of a metabolic network. bioRxiv. (doi:10.64898/2026.10.14.759191).

14. Verd B., Monk N.A., Jaeger J. 2019 Modularity, criticality, and evolvability of a developmental gene regulatory network. elife 8, e42832. (doi:10.7554/eLife.42832).

15. Jaeger J., Monk N. 2021 Dynamical modules in metabolism, cell and developmental biology. Interface Focus 11(3), 20210011. (doi:10.1098/rsfs.2021.0011).

16. Frank S.A. 2026 Protocol waists and the developmental hourglass. Proc Biol Sci 293(2075). (doi:10.1098/rspb.2026.0426).

17. Scott J.D., Pawson T. 2009 Cell signaling in space and time: where proteins come together and when they’re apart. Science 326, 1220–1224.

18. Casar B., Arozarena I., Sanz-Moreno V., Pinto A., Agudo-Ibanez L., Marais R., Lewis R.E., Berciano M.T., Crespo P. 2009 Ras subcellular localization defines extracellular signal-regulated kinase 1 and 2 substrate specificity through distinct utilization of scaffold proteins. Mol Cell Biol 29(5), 1338–1353. (doi:10.1128/MCB.01359-08).

19. Hyman A.A., Weber C.A., Julicher F. 2014 Liquid-liquid phase separation in biology. Annu Rev Cell Dev Biol 30, 39–58. (doi:10.1146/annurev-cellbio-100913-013325).

20. Lyon A.S., Peeples W.B., Rosen M.K. 2021 A framework for understanding the functions of biomolecular condensates across scales. Nat Rev Mol Cell Biol 22(3), 215–235. (doi:10.1038/s41580-020-00303-z).

21. Good M.C., Zalatan J.G., Lim W.A. 2011 Scaffold proteins: hubs for controlling the flow of cellular information. Science 332(6030), 680–686. (doi:10.1126/science.1198701).

22. Housden B.E., Perrimon N. 2014 Spatial and temporal organization of signaling pathways. Trends Biochem Sci 39(10), 457–464. (doi:10.1016/j.tibs.2014.07.008).

23. Cramer P. 2019 Organization and regulation of gene transcription. Nature 573(7772), 45–54. (doi:10.1038/s41586-019-1517-4).

24. Kilgore H.R., Chinn I., Mikhael P.G., Mitnikov I., Van Dongen C., Zylberberg G., Afeyan L., Banani S.F., Wilson-Hawken S., Lee T.I., et al. 2025 Protein codes promote selective subcellular compartmentalization. Science 387, 1095–1101.

25. Zhang Y., Jiang W., Li T., Xu H., Zhu Y., Fang K., Ren X., Wang S., Chen Y., Zhou Y., et al. 2025 SubCELL: the landscape of subcellular compartment-specific molecular interactions. Nucleic Acids Res 53(D1), D738–D747. (doi:10.1093/nar/gkae863).

26. Ferrell J.E., Jr. 2012 Bistability, bifurcations, and Waddington’s epigenetic landscape. Curr Biol 22(11), R458–466. (doi:10.1016/j.cub.2012.03.045).

27. Banani S.F., Lee H.O., Hyman A.A., Rosen M.K. 2017 Biomolecular condensates: organizers of cellular biochemistry. Nat Rev Mol Cell Biol 18, 285–298

28. Wang N., Tytell J.D., Ingber D.E. 2009 Mechanotransduction at a distance: mechanically coupling the extracellular matrix with the nucleus. Nat Rev Mol Cell Biol 10, 75–82.

29. DuFort C.C., Paszek M.J., Weaver V.M. 2011 Balancing forces: architectural control of mechanotransduction. Nat Rev Mol Cell Biol 12(5), 308–319. (doi:10.1038/nrm3112).

30. Badyaev A.V., Lee C.A., Gleason M.J., Semenov G.A., Britton S.E., Sánchez Moreno C., Duckworth R.A. 2025 Cell jamming transitions can affect regulatory protein gradients and prime evolutionary divergence. Journal of the Royal Society, Interface / the Royal Society 22, 20250186. (doi:10.1098/rsif.2025.0186).

31. Wright P.E., Dyson H.J. 2015 Intrinsically disordered proteins in cellular signalling and regulation. Nat Rev Mol Cell Biol 16(1), 18–29. (doi:10.1038/nrm3920).

32. Schneider R.A. 2024 Cellular, molecular, and genetic mechanisms of avian beak development and evolution. Annu Rev Genet 58(1), 433–454. (doi:10.1146/annurev-genet-111523-101929).

33. Rubin C.-J., Enbody E.D., Dobreva M.P., Abzhanov A., Davis B.W., Lamichhaney S., Pettersson M., Sendell-Price A.T., Sprehn C.G., Valle C.A., et al. 2022 Rapid adaptive radiation of Darwin’s finches depends on ancestral genetic modules. Sci Adv 8, eabm5982.

34. Fritz J.A., Brancale J., Tokita M., Burns K.J., Hawkins M.B., Abzhanov A., Brenner M.P. 2014 Shared developmental programme strongly constrains beak shape diversity in songbirds. Nat Commun 5, 3700. (doi:10.1038/ncomms4700).

35. Mallarino R., Campàs O., Fritz J.A., Burns K.J., Weeks O.G., Brenner M.I., Abzhanov A. 2012 Closely related bird species demonstrate flexibility between beak morphology and underlying developmental programs. Proceedings of the National Academy of Sciences of the United States of America 109, 16222–16227.

36. Abzhanov A., Kuo W.P., Hartmann C., Grant B.R., Grant P.R., Tabin C.J. 2006 The calmodulin pathway and evolution of elongated beak morphology in Darwin’s finches. Nature 442, 563–567.

37. Abzhanov A., Protas M., Grant B.R., Grant P.R., Tabin C.J. 2004 *Bmp4* and morphological variation of beaks in Darwin’s finches. Science 305, 1462–1464.

38. Badyaev A.V., Young R.L., Oh K.P., Addison C. 2008 Evolution on a local scale: Developmental, functional, and genetic bases of divergence in bill form and associated changes in song structure between adjacent habitats. Evolution 62, 1951–1964.

39. Duckworth R.A., Britton S.E., Lee C.A., Chenard K.C., Badyaev A.V. 2026 Spatial and temporal coordination of signaling pathways in tissue differentiation: developmental atlas of protein expression during zebra finch beak maturation. Developmental Dynamics 2026, 1–26. (doi:DOI: 10.1002/dvdy.70156).

40. Sánchez Moreno C., Duckworth R.A., Badyaev A.V. 2026 Craniofacial developmental atlas of protein expression of the house finch (*Haemorhous mexicanus*). FaceBase Consortium. (doi:10.25550/BN-G7QM).

41. Parsons K.J., Son Y.H., Crespel A., Thambithurai D., Killen S., Harris M.P., Albertson R.C. 2018 Conserved but flexible modularity in the zebrafish skull: implications for craniofacial evolvability. Proc Biol Sci 285(1877). (doi:10.1098/rspb.2017.2671).

42. Singh P., Ahi E.P., Sturmbauer C. 2021 Gene coexpression networks reveal molecular interactions underlying cichlid jaw modularity. BMC Ecol Evol 21(1), 62. (doi:10.1186/s12862-021-01787-9).

43. Jordan D.J., Miska E.A. 2023 Canalisation and plasticity on the developmental manifold of Caenorhabditis elegans. Mol Syst Biol 19(11), e11835. (doi:10.15252/msb.202311835).

44. Fischer E.K., Song Y., Zhou W., Hoke K.L. 2025 Flexibility in gene coexpression at developmental and evolutionary timescales. Mol Biol Evol 42(10). (doi:10.1093/molbev/msaf194).

45. Chin D., Means A.R. 2000 Calmodulin: a prototypical calcium sensor. Trends Cell Biol 10(8), 322–328.

46. Berridge M.J., Bootman M.D., Roderick H.L. 2003 Calcium signalling: dynamics, homeostasis and remodelling. Nat Rev Mol Cell Biol 4(7), 517–529. (doi:10.1038/nrm1155).

47. Liao B., Paschal B.M., Luby-Phelps K. 1999 Mechanism of Ca2+-dependent nuclear accumulation of calmodulin. Proc Natl Acad Sci USA 96, 6217–6222.

48. Tidow H., Nissen P. 2013 Structural diversity of calmodulin binding to its target sites. FEBS J 280(21), 5551–5565. (doi:10.1111/febs.12296).

49. Rajeev P., Singh N., Kechkar A., Butler C., Ramanan N., Sibarita J.B., Jose M., Nair D. 2022 Nanoscale regulation of Ca(2+) dependent phase transitions and real-time dynamics of SAP97/hDLG. Nat Commun 13(1), 4236. (doi:10.1038/s41467-022-31912-1).

50. Li Y., Ahrens M.J., Wu A., Liu J., Dudley A.T. 2011 Calcium/calmodulin-dependent protein kinase II activity regulates the proliferative potential of growth plate chondrocytes. Development 138(2), 359–370.

51. Li A., Cornelius S.P., Liu Y.Y., Wang L., Barabasi A.L. 2017 The fundamental advantages of temporal networks. Science 358(6366), 1042–1046. (doi:10.1126/science.aai7488).

52. Sunmonu N.A., Li K., Li J.Y. 2011 Numerous isoforms of Fgf8 reflect its multiple roles in the developing brain. J Cell Physiol 226(7), 1722–1726. (doi:10.1002/jcp.22587).

53. MacArthur C.A., Lawshé A., Xu J., Santos-Ocampo S., Heikinheimo M., Chellaiah A.T., Ornitz D.M. 1995 FGF-8 isoforms activate receptor splice forms that are expressed in mesenchymal regions of mouse development. Development 121(11), 3603–3613.

54. Rengarajan C., Matzke A., Reiner L., Orian-Rousseau V., Scholpp S. 2014 Endocytosis of Fgf8 is a double-stage process and regulates spreading and signaling. PLoS One 9(1), e86373. (doi:10.1371/journal.pone.0086373).

55. Scholpp S., Brand M. 2004 Endocytosis controls spreading and effective signaling range of Fgf8 protein. Curr Biol 14(20), 1834–1841. (doi:10.1016/j.cub.2004.09.084).

56. Blunt A.G., Lawshe A., Cunningham M.L., Seto M.L., Ornitz D.M., MacArthur C.A. 1997 Overlapping expression and redundant activation of mesenchymal fibroblast growth factor (FGF) receptors by alternatively spliced FGF-8 ligands. J Biol Chem 272(6), 3733–3738. (doi:10.1074/jbc.272.6.3733).

57. Le V.Q., Zhao B., Ramesh S., Toohey C., DeCosta A., Mintseris J., Liu X., Gygi S., Springer T.A. 2023 A specialized integrin-binding motif enables proTGF-beta2 activation by integrin alphaVbeta6 but not alphaVbeta8. Proc Natl Acad Sci U S A 120(24), e2304874120. (doi:10.1073/pnas.2304874120).

58. Jin M., Seed R.I., Shing T., Wang L., Li J., Cheng Y., Nishimura S.L. 2025 Latent-TGF-beta has a domain swapped architecture. Nat Commun 16(1), 10469. (doi:10.1038/s41467-025-65465-w).

59. Cockerill M., Rigozzi M.K., Terentjev E.M. 2015 Mechanosensitivity of the 2nd Kind: TGF-beta Mechanism of Cell Sensing the Substrate Stiffness. PLoS One 10(10), e0139959. (doi:10.1371/journal.pone.0139959).

60. Robertson I.B., Rifkin D.B. 2013 Unchaining the beast; insights from structural and evolutionary studies on TGFbeta secretion, sequestration, and activation. Cytokine Growth Factor Rev 24(4), 355–372. (doi:10.1016/j.cytogfr.2013.06.003).

61. Pezzulo G., Levin M. 2016 Top-down models in biology: explanation and control of complex living systems above the molecular level. Journal of the Royal Society, Interface / the Royal Society 13(124). (doi:10.1098/rsif.2016.0555).

62. Liu Y.Y., Slotine J.J., Barabasi A.L. 2011 Controllability of complex networks. Nature 473(7346), 167–173. (doi:10.1038/nature10011).

63. Watson R.A., Wagner G.P., Pavlicev M., Weinreich D.M., Mills R. 2014 The evolution of phenotypic correlations and “developmental memory”. Evolution 68(4), 1124–1138. (doi:10.1111/evo.12337).

64. Huang S., Eichler G., Bar-Yam Y., Ingber D.E. 2005 Cell fates as high-dimensional attractor states of a complex gene regulatory network. Phys Rev Lett 94(12), 128701. (doi:10.1103/PhysRevLett.94.128701).

65. Oster G.F., Murray J.D., Harris A.K. 1983 Mechanical aspects of mesenchymal morphogenesis. Journal of Embryology and Experimental Morphology 78(1), 83–125. (doi:10.1242/dev.78.1.83).

66. Beloussov L.V. 2008 Mechanically based generative laws of morphogenesis. Phys Biol 5(1), 015009. (doi:10.1088/1478-3975/5/1/015009).

67. Waddington C.H. 1956 The Principles of Embryology. New York, The MacMillan Co.

68. Wilkins A.S. 2001 The Evolution of Developmental Pathways. Sunderland, MA, Sinauer Associates; 603 p.

69. Bargaje R., Trachana K., Shelton M.N., McGinnis C.S., Zhou J.X., Chadick C., Cook S., Cavanaugh C., Huang S., Hood L. 2017 Cell population structure prior to bifurcation predicts efficiency of directed differentiation in human induced pluripotent cells. Proc Natl Acad Sci U S A 114(9), 2271–2276. (doi:10.1073/pnas.1621412114).

70. Huang S. 2012 The molecular and mathematical basis of Waddington’s epigenetic landscape: a framework for post-Darwinian biology? BioEssays 34(2), 149–157. (doi:10.1002/bies.201100031).

71. Selleri L., Rijli F.M. 2023 Shaping faces: genetic and epigenetic control of craniofacial morphogenesis. Nat Rev Genet 24(9), 610–626. (doi:10.1038/s41576-023-00594-w).

72. Foppiano S., Hu D., Marcucio R.S. 2007 Signaling by bone morphogenetic proteins directs formation of an ectodermal signaling center that regulates craniofacial development. Dev Biol 312(1), 103–114. (doi:10.1016/j.ydbio.2007.09.016).

73. Sánchez Moreno C., Badyaev A.V. 2026 Transient epithelial mimicry by neural crest mesenchyme anchors cell condensations across avian beaks. Evolution & Development 28, e70044. (doi:10.1111/ede.70044).

74. Lee C.A., Sánchez Moreno C., Badyaev A.V. 2024 FInCH: FIJI plugin for automated and scalable whole-image analysis of protein expression and cell morphology. MethodsX 13, 102855.

75. Specht H., Peterziel H., Bajohrs M., Gerdes H.H., Krieglstein K., Unsicker K. 2003 Transforming growth factor beta2 is released from PC12 cells via the regulated pathway of secretion. Mol Cell Neurosci 22(1), 75–86. (doi:10.1016/s1044-7431(02)00023-4).

76. Suzuki A., Harada H., Nakamura H. 2012 Nuclear translocation of FGF8 and its implication to induce Sprouty2. Dev Growth Differ 54(4), 463–473. (doi:10.1111/j.1440-169X.2012.01332.x).

77. Dash S., Trainor P.A. 2023 Correction: Nucleolin loss of function leads to aberrant Fibroblast Growth Factor signaling and craniofacial anomalies. Development 150(6). (doi:10.1242/dev.201584).

78. Xu J., Huang Z., Wang W., Tan X., Li H., Zhang Y., Tian W., Hu T., Chen Y.P. 2018 FGF8 Signaling Alters the Osteogenic Cell Fate in the Hard Palate. J Dent Res 97(5), 589–596. (doi:10.1177/0022034517750141).

79. Badyaev A.V., Sanchez Moreno C., Lee C.A., Britton S.E., Johnstone L.M., Duckworth R.A. 2026 Ultimate paths of least resistance: intrinsically disordered proteins as developmental resets in regulatory networks. Proc Biol Sci 293(2062). (doi:10.1098/rspb.2025.2393).

80. van Aert R.C.M. 2023 Meta-analyzing partial correlation coefficients using Fisher’s z transformation. Res Synth Methods 14(5), 768–773. (doi:10.1002/jrsm.1654).

81. Farahbod M., Pavlidis P. 2019 Differential coexpression in human tissues and the confounding effect of mean expression levels. Bioinformatics 35(1), 55–61. (doi:10.1093/bioinformatics/bty538).

